# Space-Time Mapping Identifies Concerted Multicellular Patterns and Gene Programs in Healing Wounds and their Conservation in Cancers

**DOI:** 10.1101/2022.05.25.493500

**Authors:** Kenneth H. Hu, Nicholas F. Kuhn, Tristan Courau, Matthew F. Krummel

## Abstract

Tissue repair responses in metazoans are highly coordinated by different cell types over space and time. However, comprehensive single-cell based characterization covering this coordination is lacking. Here, we captured transcriptional states of single cells over space and time during skin wound closure, revealing choreographed gene expression profiles. We identified shared and prominent space-time patterns of cellular and gene expression enrichment: which we call multicellular ‘movements’ and which spanned multiple cell types. We validated some of the discovered space-time movements using large volume imaging of cleared wounds and demonstrated the value of this analysis to predict gene products made by macrophages or fibroblasts, which activated gene programs in the opposite cell type. Finally, using two different tumor models, we tested the hypothesis that tumors are like ‘wounds that never heal’ finding conserved wound healing movements in the tumor space, wherein some movements were preferentially used in one tumor versus another.

**Graphical Abstract:** 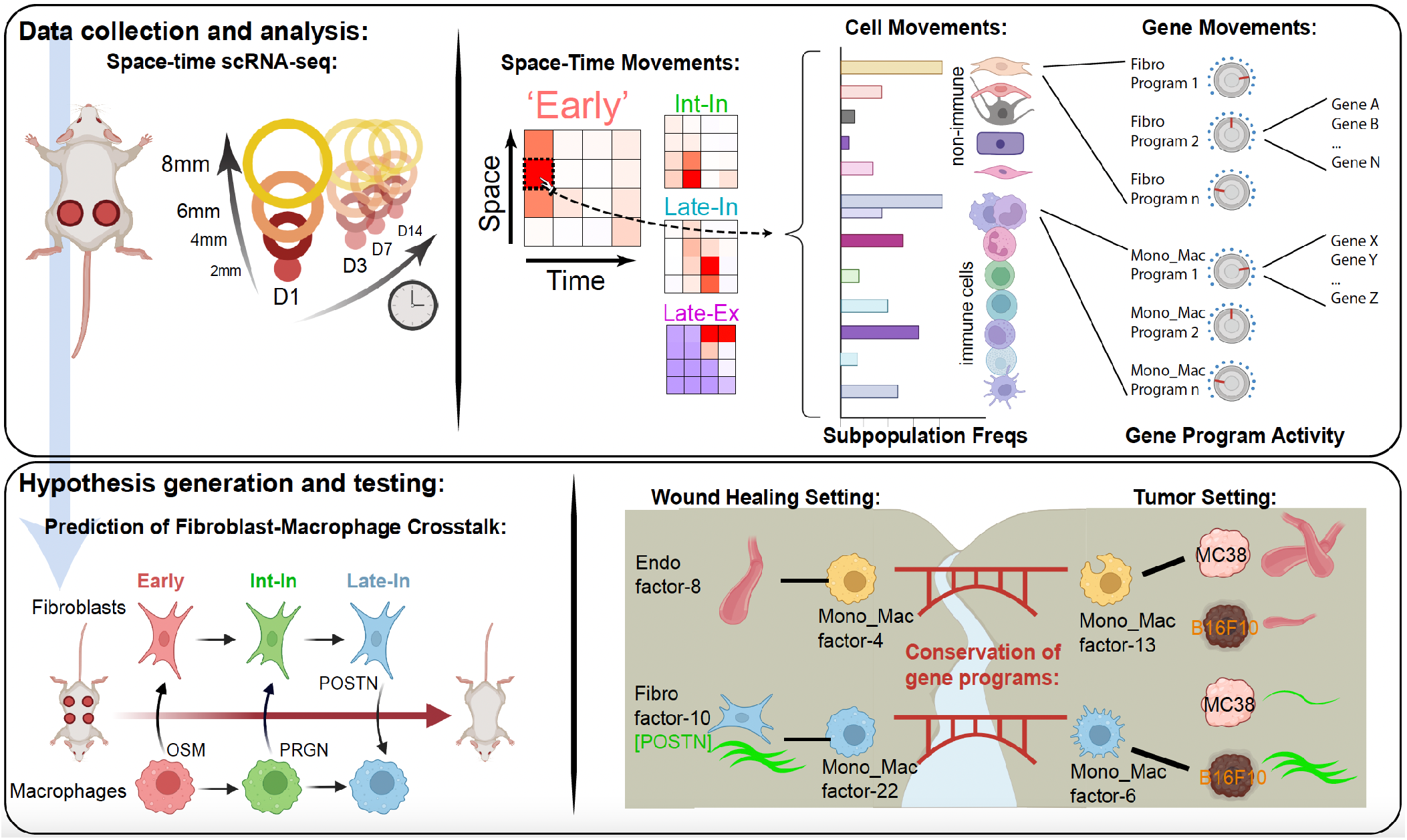

## Introduction

Metazoans rely on intricate networks of cell-cell crosstalk for the maintenance of tissue homeostasis and during repair and regenerative processes following perturbations such as damage^1–3^. Given the diversity of cell types within a tissue and all possible ligand-receptor pairings and their signaling dynamics, a formalized method for interrogating crosstalk over space and time in the tissue remains a daunting task^4^. Even a minimal two-actor system can exhibit robustness and return to a stable state following perturbation^5^. This same adaptation to perturbation can be seen when increasing the number of cellular actors and thus the number of possible ‘edges’ (i.e. cell-cell crosstalk axes), such as the combination of stellate cells, hepatocytes, endothelial cells, and Kupffer cells in a liver-specific niche^6^. Identification of these cell-cell crosstalk mechanisms and effects on gene expression across a multitude of cells at once requires the ability to measure expression patterns within single cells isolated from the tissue, during the process.

To that end, the advent of single-cell technologies has allowed profiling of cells on the transcriptional level at resolutions previously unattainable, generating rich datasets identifying highly-resolved cell subsets and subtle variations in gene expression and activation states^7–9^. A number of computational approaches have sought to infer cell-cell communication via paired ligand-receptor expression^10,11^ or ligand and downstream target gene expression^12^. These inferences can be further strengthened by applying spatial and temporal context to single cell transcriptomics. Spatially resolved single cell datasets, such as those generated along the length of the intestinal villus, have revealed groupings of genes with similar spatiotemporal profiles, shedding light on the spatial segregation of cell functions, the dynamics of cell migration, and cell-cell crosstalk that enforced zonation^13,14^. Similarly, describing gene expression in terms of spatiotemporal patterns revealed signaling pathways and co-regulation of genes in the liver^15^ and in peri-islet acinar cells in the pancreas^16^. Concordantly, we were motivated to describe the healing skin wound in terms of spatiotemporal multicellular patterns and gene expression programs. Skin wound healing naturally displays well-defined spatial and temporal dimensions following the advancing edge of the wound^17^. This process has canonically been segmented into several major phases starting with an initial inflammatory response followed by repair/growth and a resolution phase^18,19^. Interspersed within these phases are coordinated changes in gene expression patterns in diverse cellular players from monocyte/macrophages, neutrophils, fibroblasts, endothelial cells, keratinocytes and beyond^3,20^. Diverse crosstalk mechanisms between these cell types have been identified for regulating the proper duration of and transition between phases^3,20–23^ and disruption of these mechanisms often results in delayed or aberrant healing, demonstrating the highly interdependent structure of the wound healing cellular network. For example, depletion of macrophages at different stages of wound healing results in delayed healing with different phenotypes, again reinforcing the idea that cellular choreography is essential with distinct functional roles during each phase^24,25^. Charting the progression of gene expression in single cells over space and time in the wound in a holistic manner would likely yield information on the coordinated behaviors of myriad cell types in an unbiased manner, leading to identification of new mechanisms of cellular crosstalk (i.e. identification of new edges in our network) and how they drive transitions between different phases of healing.

Additionally, the plasticity of cell types such as macrophages^26^ and fibroblasts^27,28^ and the everchanging milieu of cell signaling factors result in these cells occupying a continuum of expression states, suggesting that clustering approaches may be insufficient to capture the transitions between states. For example, the M1/M2 ‘binning’ of macrophages based on a handful of canonical marker genes (e.g. *Arg1, Il6, Vegfa*, and *Mrc1*) may represent too reductive a model, as wound healing macrophages express combinations of canonical M1/M2 genes all across the timescale of wound healing^29^. Therefore, a method for reframing cellular heterogeneity using overlays of gene programs (collections of co-expressed genes) in the healing wound may better capture the biology underlying the progression of cellular transcriptional heterogeneity.

As such, the healing skin wound represents an ideal testbed for layering spatial and temporal dimensions onto single cell transcriptomics to identify grouped gene program progression across immune and non-immune cell types and thus infer the presence of a coordinated series of cell-cell interactions.

An additional important rational for studying space-time progression of multicellular networks relates to chronic disease, in which healthy resolution is never achieved. Previous observations—largely at the level of tissue morphology—have suggested that tumors are ‘wounds that never heal’, namely that they co-opt wound healing but do not progress from inflammatory or resolution phases and fail to conclude^30–32^. This idea motivated us to develop a framework based on conserved gene programs to identify whether crosstalk elements of the wound healing cellular network are ‘borrowed’ by growing tumors and whether they do so similarly across tumor models. The heterogeneity of a given cell type in different contexts may represent a convolution of conserved differentiation, functional, and tissue-specific expression patterns, as seen in resident immune cells scattered across all tissues^33,34^. Describing the common biology between two single cell datasets may again require going beyond clustering based approaches which may obscure the identification of overlaid gene programs in a continuum of cell states^35^.

Using skin wound healing as a well-defined spatial phenomenon in tissue homeostasis and repair, we mapped changes in cell identity that co-occurred in similar space-time patterns, within the CD45^+^ immune and CD45^-^ non-immune compartment during the wound healing process. Layered on top of cell type identity, we also identified spatiotemporally orchestrated gene programs – or factors – that can be grouped based on their unique space-time profile. Because we found these factors on the basis of their shared space-time patterns across multiple cell types, we refer to these shared factors as multicellular ‘movements’. Informed by spatiotemporal profiles of gene program expression, we focused on a handful of fibroblast-macrophage interactions over the time-course of wound closure, which we then verified using orthogonal approaches such as *in vitro* experiments and whole mount cleared tissue imaging. We studied these groupings, which allowed us to predict cell-cell crosstalk and validate a handful of pathways that had effects on target gene expression in paired cell types. Finally, we derived a framework for how to identify movements across tissue contexts, predicting the conservation of correlated immune and non-immune gene program pairs and validating their variable presence within solid tumor models.

## Results

### Separate waves of immune cell infiltration during wound closure

To establish the compositional changes of immune cells during skin repair, full thickness circular wounds of 4 mm diameter were made on the back of mice (**Figure 1A**). We then sampled the wounds from day 0 to day 14 as they healed (**Figure S1A**). Sampling of CD45^+^ immune cells – collected within a 4 mm radius of the wound center – during and post wound closure via mass cytometry using a 40-marker panel identified waves of major myeloid and lymphoid populations (**Figures 1B and S1B**). At our sampling resolution, we found three waves, dominated by the infiltration/presence of distinct immune cells (**Figure 1C**). *Early*: Up to one day post wounding, wounds were characterized by an influx of Ly6C^hi^ inflammatory monocytes (infl. Monocytes 1-3) and mature Ly6G^hi^ neutrophils (**Figure 1D**). Other immune cell subsets were mostly depleted during these first 24 hours post skin wounding. Starting on day 3, Ly6G^-^ c-Kit^+^ F4/80^+^ CD24^+^ immature pre-neutrophils were found in the wound bed (**Figures 1D and S1C**), which phenotypically match a recently described proliferative neutrophil precursor in the bone marrow^36^. *Mid*: Days 3 to 10 marked the peak emergence of three myeloid cells populations that had increasing levels of MHCII expression and concordant decrease of Ly6C levels (**Figure 1D**), as seen in maturing blood-derived monocytes extravasating into inflamed skin and gut^37–39^. Based on Ly6C, MHCII, CD64, F4/80, and CD11c expression we annotated them as Ly6C^int^ monocytes (Ly6C^int^ MHCII^mid^ CD64^mid^ F4/80^low^ CD11c^low^), macrophages (Ly6C^low^ MHCII^+^ CD64^+^ F4/80^+^ CD11c^mid^), and monocyte-derived dendritic cells (DCs) (Ly6C^-^ MHCII^+^ CD64^-^ F4/80^-^ CD11c^+^) (**Figure S1B**). This highlights a coordinated wave of myeloid cell emergence in the wound bed, with inflammatory monocytes and neutrophils predominantly inhabiting the wound in the first day, transitioning to macrophage and monocyte-derived DC populations with increasing MHCII expression. *Late*: In addition to the most mature macrophages, MHCII-expressing dendritic cells also arrive on day 14 when the wound is closed (**Figure 1D**). A further nuance of the late phase is the accumulation of some cell populations that were immediately lost upon wounding, many to the levels well beyond the unwounded state. These include Macrophage_3 populations as well as T cell populations, such as CD4^+^, CD8^+^, γδ, and dermal γδ T cells^40^ (**Figures 1D and S1D**). All these T cell populations reach pre-wound levels, except CD4^+^ T cells, which rebounded to a higher level than pre-wound. That cluster is composed of Foxp3^+^ regulatory T cells, γδ T cell receptor (TCR)^+^ T cells and conventional CD4^+^ T cells (**Figure S1D**). Among those, the γδTCR^-^ CD4^+^ Foxp3^-^ T cells rebound the earliest and to the highest level (**Figure S1E**).

**Figure 1.**
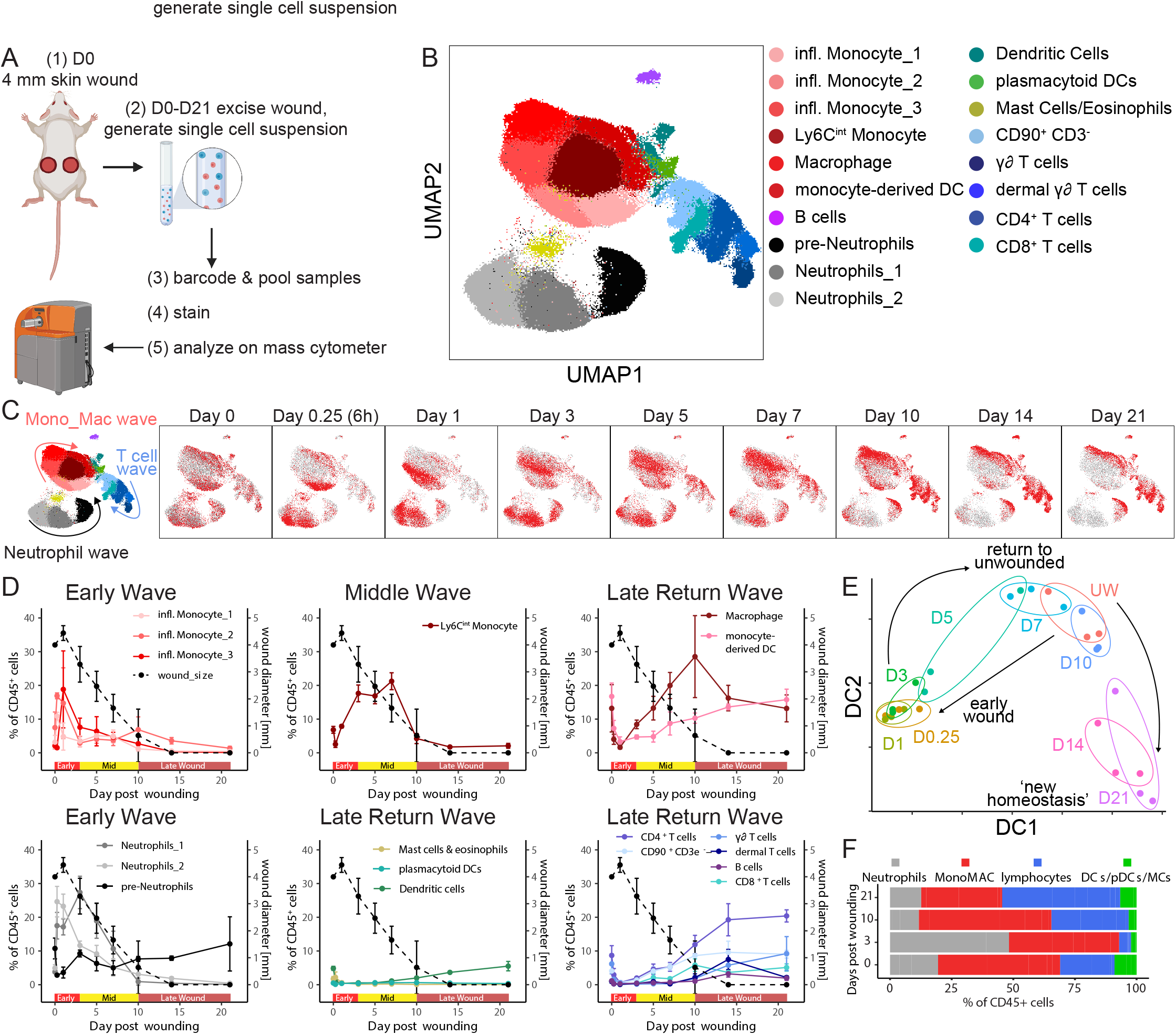
Separate waves of immune cell infiltration during wound closure. (**A**) Experimental layout of immune cell profiling during skin wound repair. Full thickness circular dorsal skin wounds of 4 mm diameter were caused in mice. Wound and surrounding tissue was collected using an 8 mm diameter punch biopsy. A single cell suspension was generated prior to barcoding, staining, and analysis on a mass cytometer. (**B**) UMAP projection of immune cells detected during skin wound repair based on a 40-marker panel. (**C**) UMAP projection of all immune cells from (B) in gray and cells highlighted in red by timepoint of wound sampling. (**D**) Line plots of all immune cell clusters identified in (B) plotted as percentage of all CD45^+^ immune cells per timepoint of wound sampling (left y-axis). Dotted line represents wound diameter (right y-axis). Dots represent mean values of n=3 mice per timepoint. Error bars represent ±SD. (**E**) Phenotypic earth mover’s distance (PhEMD) diffusion map embedding of all samples collected during wound closure. Each dot represents the wound-resident immune cell composition of one mouse. Dots are color-coded by day and n=3 mice were collected per timepoint. UW, unwounded. DC, diffusion coefficient. (**F**) Horizontal stacked bar chart of major immune cell populations plotted as percentage of all CD45^+^ cells during skin wound repair. DCs, dendritic cells. pDCs, plasmacytoid dendritic cells. MCs, mast cells.

To globally describe the dynamic changes in immune cell composition during skin repair, we applied ‘phenotypic earth mover’s distance’ (PhEMD) analysis to our dataset^41^. This dimensionality reduction revealed that day 1 post-wounding samples are very dissimilar to the unwounded state (**Figure 1E**). Day 3 and day 5 samples displayed higher similarity to the unwounded state, with day 7 and day 10 post-wounding samples showing the most similar immune cell composition to unwounded state. The timepoints day 10 and day 14 mark the time of wound closure (**Figure S1A**), suggesting that upon epithelial skin repair, the skin-resident immune cell composition reaches pre-wounding homeostasis. However, day 14 and day 21 samples ‘overshoot’ this state and arrive at a distant state from all other samples (**Figure 1E**). This difference is driven by a decrease in neutrophils and monocyte-macrophage cells, and a corresponding increase in the lymphocyte population (**Figure 1F**). These findings were recapitulated in a second experimental run, where identical cell subpopulations and dynamics were observed (**Figures S1F-S1H**).

In summary, the immune cell population infiltrating the skin wound displays a dynamic turnover, wherein it temporarily reaches the pre-wound composition but eventually assumes a ‘new homeostasis’ of skin-resident immune cell population that is lymphocyte-rich. To gain a more comprehensive understanding on these state changes, we developed a spatial single cell transcriptional analysis of skin wound repair encompassing immune and non-immune cells and, for the present, focused upon the first 14 days which encompassed the majority of resolution.

### Transcriptional space-time analysis of single cells unveils unique cell patterns during skin repair

Involvement of both immune and non-immune cells are crucial for skin repair^42–44^. To understand how gene expression programs in the different phases of skin repair are linked amongst cell types and across the radial dimensions of a wound, wounds were physically and radially partitioned using successively large punches: wound center (2 mm center diameter), wound edge (2-4 mm diameter), wound proximal (4-6 mm diameter), and wound distal (6-8 mm diameter). These different areas were sampled during different phases of wound repair: inflammatory (D1), transition (D3), return-to-unwounded (D7), and ‘overshoot/new homeostasis’ (D14) (**Figures 1E and 2A**). Given the original wound diameter of 4 mm, we thus captured two distinct areas within the wound bed plus two distinct areas in skin beyond the wound and into uninjured tissue. Upon digestion of the individual rings of tissue, CD45^+^ immune and CD45^-^ non-immune cells were sorted separately, barcoded using MULTI-seq^45^ based on their collected timepoint and area, and then their transcriptional state assessed by using the 10x Genomics single-cell RNA sequencing (scRNAseq) platform (**Figure 2A**). This provided us with a space-time scRNAseq dataset that covers the major extent of the skin repair process.

**Figure 2.**
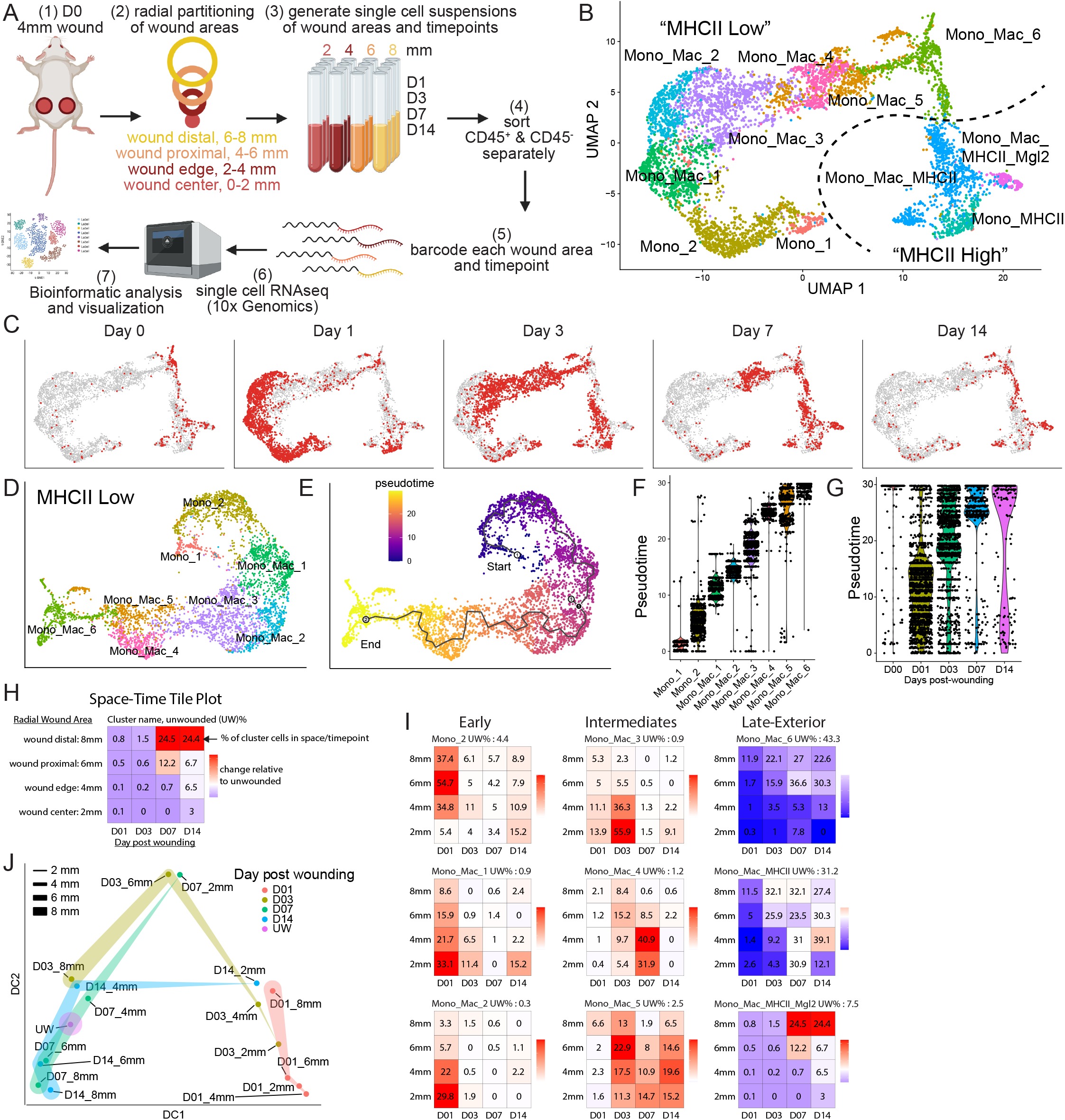
Transcriptional space-time analysis of single cells unveils unique patterns of monocyte/macrophage populations during skin repair. (**A**) Experimental layout of transcriptional space-time analysis in wounded skin, including radial partitioning of wound area into four zones, generation of single cell suspension at four time points, sorting of CD45^+^ and CD45^-^ cells, barcoding, cell capture by droplet-based device, sequencing, and downstream analysis. (**B**) UMAP plot of monocyte/macrophage subset from CD45^+^ object. Dotted line separates eight MHCII^lo^ and three MHCII^hi^ monocyte/macrophage clusters. (**C**) UMAP projection of all monocyte/macrophage cells from (B) in gray and cells highlighted in red by timepoint of wound sampling. (**D**) UMAP of MHCII^lo^ monocyte/macrophage subset with eight distinct clusters. (**E**) Pseudotime trajectory using Monocle 3 on MHCII^lo^ monocyte/macrophage subset starting at cluster Mono_1, progressing through all cell clusters and ending at cluster Mono_Mac_6. (**F**) Violin plot split by the MHCII^lo^ monocyte/macrophage subpopulations and plotted according to their distribution in pseudotime. (**G**) Violin plot of all MHCII^lo^ monocyte/macrophage cells split by day post-wounding and plotted according to their presence in pseudotime. D00 represents unwounded skin. (**H**) Outline of Space-Time tile plot. The 4×4 grid depicts relative abundance of a cell cluster across radial wound area (y-axis) and time post wounding (x-axis). Each tile is one space-timepoint. Number in tile is the percentage of a subpopulation among all Mono_Mac cells at specific space-timepoint. Color indicates relative change compared to unwounded (UW) state. Red indicates increase in subpopulation, blue indicates decrease in subpopulation compared to UW, white indicates no change. Exemplary data is depicted. (**I**) Space-time tile plots representing Mono_Mac subpopulations. (**J**) Phenotypic earth mover’s distance (PhEMD) diffusion map embedding of all space-timepoints. Each dot represents all CD45^+^ immune cells captured by scRNAseq within that space-timepoint. Dots are color-coded by day and width of band corresponds to area sampled. DC, diffusion coefficient.

First, we focused on the CD45^+^ immune cells. Graph-based clustering and differential gene expression analysis of 10,492 cells revealed heterogenous monocyte and macrophage subpopulations, four neutrophil subpopulations, mast cells, dendritic cells, B cells, two T cell, and one Natural Killer (NK) cell cluster (**Figure S2A and S2B**). After re-clustering the monocyte-macrophage (from here on referred to as Mono_Mac) populations as the largest representative of CD45^+^ immune cells, we identified three MHCII^hi^ and eight MHCII^lo^ subpopulations (**Figures 2B and S2C**).

We noted that MHCII^hi^ cells expressed markers associated with monocyte-derived dendritic cells (moDCs), such as *Cd209a*^46^ and the transcription factor *Nr4a3*^47^ (**Figure S2C and S2D**). Additionally, both MHCII^lo^ and MHCI^hi^ Mono_Macs emerged together and early during wound healing (**Figure 2C**, Day1). On these bases, we considered them separately, reclustering them prior to further analysis (**Figures 2D and S2E**, respectively).

The MHCII^lo^ Mono_Mac subset consisted of two subsets expressing high levels of *Ly6c2*, which we annotated as Mono_1 and Mono_2 (**Figure S2C**). Those, together with Mono_Mac_3 also were identified by upregulation of the transcription factor *Klf2*, which regulates pro-inflammatory cues in monocytes^48^ and at least one of the chemokines *Cxcl3, Ccl2, Ccl6, Ccl7*, and *Ccl24*. Finally, Mono_Mac_4, Mono_Mac_5, and Mono_Mac_6 shared *Ms4a7* and *Apoe* expression, two markers associated with microglia and brain-border macrophages^49,50^. Using pseudotime analysis^51^, we anchored a trajectory on Mono_1 and Mono_2 populations, representing cells most similar to Ly6C^hi^ blood-circulating monocytes (**Figure S2C**), which then proposed a linked transcriptional progression through all clusters successively, eventually ending at Mono_Mac_6, putatively the most differentiated Mono_Mac cluster (**Figures 2E and 2F**). Consistent with this interpretation, Mono_Mac MHCII^lo^ cells from different timepoints map along the calculated pseudotime (**Figure 2G**) with the exception that cells collected on day 14 were split between early and late pseudotime, highlighting again the new homeostasis in wounded skin that was also found in the CyTOF data (**Figure 1E**). Leveraging this trajectory, we also found trends of gene expression characteristic of known myeloid cell states: early expression of inflammatory genes *Ifit1, Ly6c2*, and *Plac8*^52^, mid-point expression of *Slpi* and *Mmp12*, with *Mmp12* being linked to antagonizing further monocyte recruitment^53^, and late expression of genes *Mrc1, Retnla*, and *C1qc* associated with tissue-resident macrophages (**Figure S2F**).

When applying the same analysis to MHCII^hi^ Mono_Macs, anchored on the Mono_MHCII subset, we again found that different clusters show distinct positions within pseudotime (**Figures S2G**), with Mono_Mac_MHCII cells emerging at the intermediate to late stage, representing a transition state to Mono_Mac_MHCII_Mgl2 cells, which emerge very late in pseudotime (**Figure S2H-I**).

To search for patterns of space-time distribution of myeloid states, we created tile plots in which each tile shows the prevalence of a cell population—defined from the clustering, above—relative to the unwounded state, both over time (x dimension) and across the wound diameter (y dimension) (**Figure 2H**). These could be created using cell-type frequencies (**Figure 2I**) or relative to pseudotime (**Figure S2J**). The latter demonstrates the finding that MHCII^lo^ cell populations most prevalent early in pseudotime were overrepresented on day 1 across the whole wound, and modestly again at day 14 specifically at the wound center (**Figure S2J**). Applying such an analysis to the MHCII^hi^ grouping shows that early pseudotime MHCII^hi^ Mono_Macs appear most predominantly at the center of the wound (**Figure S2K**, see D01_2mm), highlighting their different space-time trajectory within the healing skin wound and likely different origin compared to MHCII^lo^ Mono_Macs.

When we ordered the tile plots in order of cell type appearance, we discovered greater spatial complexity in how specific immune populations (**Figure S2L-N**), exemplified by the Mono_Macs (**Figure 2I**), emerge through wound healing phases. Thus, three subpopulations of these were characterized as ‘Early’ (i.e. peak at day 1)—represented by Mono_2, Mono_Mac_1 and Mono_Mac_2 and varied from 6 mm to 2 mm in the peak location of the amplified population. Three others had middle-originating waves (centered on D3 or D7): denoted as ‘Intermediates’ and represented by Mono_Mac_3, Mono_Mac_4, Mono_Mac_5, respectively. Each of these again had specific space-time patterns of recruitment, with Mono_Mac_2 exhibiting an ‘Intermediate Interior (Int-In)’ and Mono_Mac_4 a ‘Late Interior (Late-In)’ pattern. Three others had ‘Late Exterior (Late-Ex)’ distributions, namely Mono_Mac_6, Mono_Mac_MHCII, and Mono_Mac_MHCII_Mgl2 and these were noted as ‘return’ populations, namely that the frequency for much of the repair process was lower than in the unwounded state. A few others, notably the earliest progenitor Mono_1, as well Mono_MHCII, did not fit into those patterns (**Figure S2L-M**).

Our spatial scRNAseq dataset also captured the space-time accumulation of additional immune cells beyond monocytes and macrophages (**Figure S2N**). Using PhEMD analysis, the samples when viewed across these additional populations occupied an arc-like trajectory in reduced dimensions (**Figure 2J**), on the face similarly to that seen in bulk analysis in **Figure 1**. In this analysis, late wound outside areas (D07_6mm, D07_8mm, D14_6mm, D14_8mm) are more similar to each other than to the unwounded state, again supporting the previous observation that the immune composition reaches a new state after skin repair, dissimilar to pre-injury (**Figures 1E and 2J**).

### Large volume imaging visualizes spatial distribution of Mono_Mac subsets in wounded skin

We aimed to validate and extend our findings for the Mono_Mac compartments in a setting of intact tissue architecture that did not rely on microdissection. We thus utilized large volume tissue imaging of wounded skin to probe for the location of individual cell states relative to the wounded area (**Figure 3A**). The clearing-enhanced 3D (Ce3D)^54^ protocol allowed detection of immune and non-immune cells within wounded skin, highlighting their relative position to the wound edge as marked by integrin alpha-6 (ITGA6) staining of the re-established epithelial basement membrane (**Figure S3A** and **Movie S1**, white). Of note, ITGA6 staining also highlighted vasculature structures in non-wounded skin, hair follicles, fascia, and severed nerve bundles lying outside of the closing wound (**Figure S3B and Movie S2**, white).

**Figure 3.**
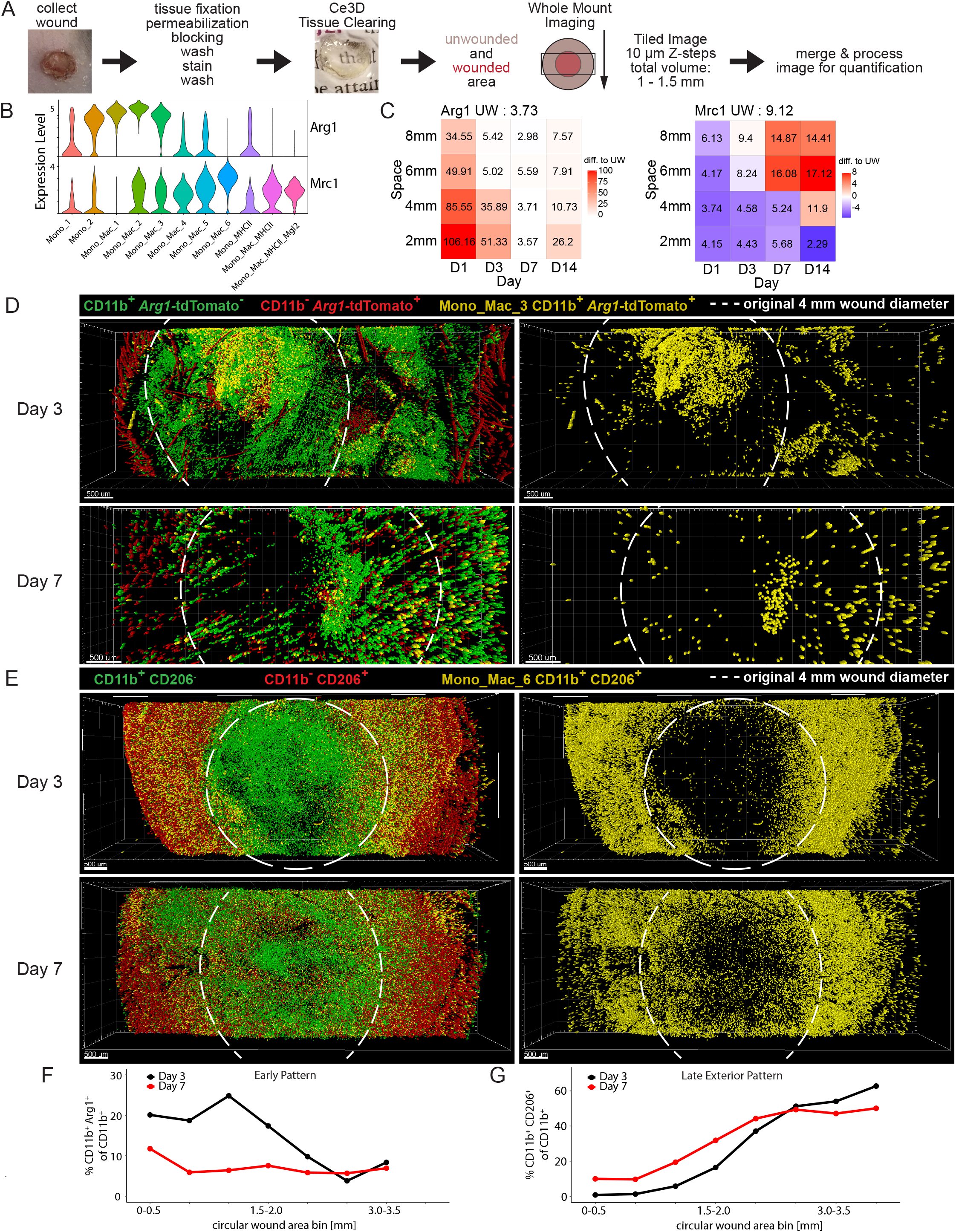
Large volume imaging visualizes spatial distribution of Mono_Mac subsets in whole wounds during skin repair. (**A**) Workflow for large volume imaging. Wounds are collected by skin excision, processed for staining, and then cleared using the Ce3D protocol. A rectangular cuboid covering the wound and surrounding unwounded skin tissue is acquired on a scanning confocal microscope. Acquired images are stitched together and processed for image analysis using Imaris. (**B**) ViolinPlot of (left) *Arg1* and (right) *Mrc1* natural log-normalized mRNA expression level within all Mono_Mac subpopulations. (**C**) Space-time tile plot of (left) *Arg1* and (right) *Mrc1* mRNA expression (normalized to depth) within all Mono_Mac cells. Tiles are color-coded relative to unwounded (UW) state. Red, high. Blue, low. (**D**) Top-down view of processed image from *Arg1*-tdTomato reporter mouse on (top) day 3 and (bottom) day 7 post-wounding. Colored dots indicate cell location of *Arg1*^-^ CD11b^+^ Mono_Mac (green), *Arg1*^+^ CD11b^-^ non-myeloid cells (red), and *Arg1*^+^ CD11b^+^ Mono_Mac_3 (yellow). Dotted line represents original 4 mm wound diameter. Images on the right show only *Arg1*^+^ CD11b^+^ Mono_Mac_3 cells (yellow). Bar, 500 μm. One of two representative images is shown for each day. (**E**) Top-down view of processed image from wild-type C57BL/6 mouse on (top) day 3 and (bottom) day 7 post-wounding. Colored dots indicate cell location of CD206^-^ CD11b^+^ Mono_Macs (green), CD206^+^ CD11b^-^ non-myeloid cells (red), and CD206^+^ CD11b^+^ Mono_Mac_6 (yellow). Dotted line represents original 4 mm wound diameter. Images on the right show only CD206^+^ CD11b^+^ Mono_Mac_6 cells (yellow). Bar, 500 μm. One of two representative images is shown for each day. (**F**) Quantification of CD11b^+^ *Arg1*^+^ cells in (D) relative to distance from the center of the wound. (**G**) Quantification of CD11b^+^ CD206^+^ cells in (E) relative to distance from the center of the wound.

We focused on two gene products—*Arg1* and *CD206/Mrc1* that, though often grouped together as a combined signature of ‘M2’ macrophages *in vitro*^55^, show clear cell subset (**Figure 3B**) and space-time distinct (**Figure 3C**) patterns in wound healing, further confirming results from tumors^56^ that they are not obligately part of the same gene network. By scRNAseq data, ‘early’ inflammatory Mono_Mac subpopulations Mono_Mac_1, Mono_Mac_2, and Mono_Mac_3 express notably high levels of *Arg1* (**Figures 3B and S3C**). We validated this via cleared wound images using an *Arg1* reporter mouse^57^, finding CD11b^+^ *Arg1*-tdTomato^+^ myeloid cells were predominantly found at the center of wounds and only at early but not late timepoints after wounding (**Figure 3D, 3F, S3D and S3E; Movies S3 and S4**), consistent with the space-time tile plot (**Figure 3C**). Confirming previous data^58^, we also detected CD11b^-^ *Arg1*-tdTomato^+^ hair follicles in unwounded skin (**Figure S3E**). Conversely, *Mrc1* gene expression was highest in the Mono_Mac_6 subpopulation and predominated in exterior wound regions late in the response (**Figures 3B, 3C and S3F**). Ce3D-cleared large volume imaging using antibodies against the *Mrc1*-encoded protein CD206 confirmed the absence of CD11b^+^ CD206^+^ cells in the center of the wound on day 3 post-wounding and their increased presence on the exterior non-wounded skin area (**Figures 3E and 3G, S3G, and S3H; Movies S1 and S2**). Together, these data demonstrate the unique spatial and temporal dynamics of two different Mono_Mac subsets during wound repair, found using this analysis.

### Unique space-time patterns of fibroblast subpopulations have matching Mono_Mac patterns

Taking the previously identified patterns of Mono_Macs as an exemplar, we sought to use space-time mapping to determine whether other cell types have subsets with matched patterns. Unsupervised clustering of 6944 CD45^-^ cells identified 19 different clusters, which could broadly be separated into endothelial cells, fibroblasts, melanocytes, muscle cells, keratinocytes, and hair follicle-associated dermal sheath papilla cells (**Figure 4A and S4**). Similar to immune populations, many cell types such as fibroblasts (**Figures 4B-4C**), keratinocytes (**Figures S4B-S4F**), and other skin-resident cells (**Figures S4G-S4O**) displayed distinct space-time distributions. For our major analysis, to seek associated cell states, we primarily focused on fibroblasts as their interaction with macrophages is well-documented^59^. Here, we identified 5 separate fibroblast clusters by distinct gene expression (**Figures 4B and S4P**) and numbered them in accordance with their accumulation in time (**Figure 4C**).

**Figure 4.**
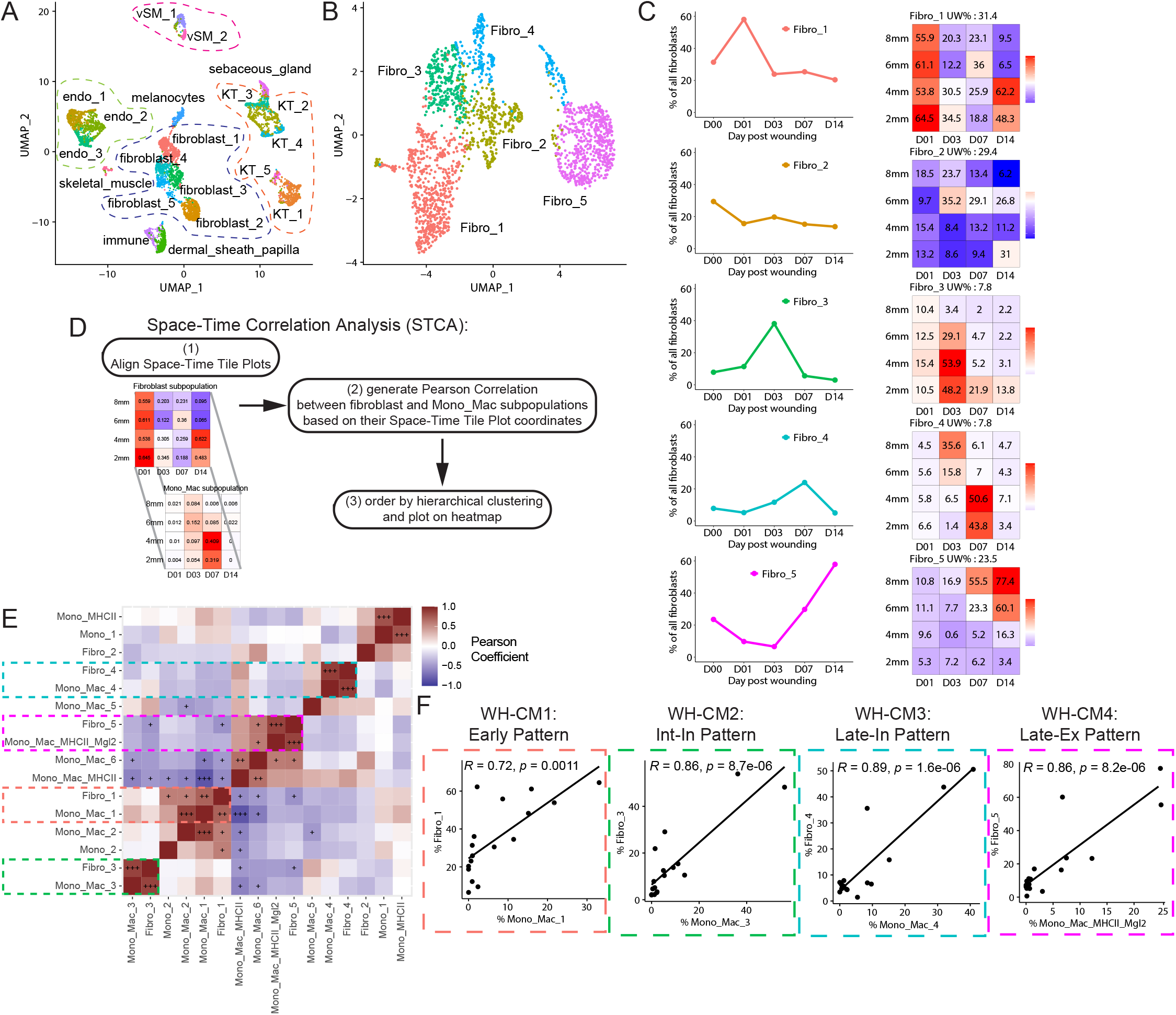
Fibroblast subpopulations have specific patterns in common with particular Mono_Mac subpopulations. (**A**) UMAP plot of CD45^-^ non-immune cells found during skin repair. Endo, endothelial cells. KT, keratinocytes. vSM, vascular smooth muscle. (**B**) UMAP plot of the fibroblast subset found during skin repair. (**C**) Left: Line plots of fibroblast subpopulations identified in the scRNAseq dataset during skin repair. Percentage of each subpopulation within all fibroblasts is plotted by day post-wounding. Right: Spacetime tile plot representing the fibroblast subpopulations. Each tile is one space-timepoint. Number in tile is percentage of subpopulation among all fibroblasts at specific space-timepoint. Color indicates relative change compared to unwounded (UW) state. Red indicates increase in subpopulation, blue indicates decrease in subpopulation compared to UW. (**D**) Outline of Space-Time Correlation Analysis (STCA). (**E**) Pearson correlation matrix output of STCA comparing all fibroblast and Mono_Mac subpopulations. +++p-value < 0.05, ++p-value < 0.005, +++p-value < 0.0005. (**F**) Pearson correlation xy-plots of select fibroblast-Mono_Mac subpopulation pairs displaying high correlation in occurrence in space-time during wound skin repair. WH-CM, wound healing cell movement.

The Fibro_1 cluster exhibited high expression of inflammatory mediators, such as the chemokine *Cxcl5* and the TNFα-induced opsonin *Ptx3* (**Figure S4P**). Jaccard similarity analysis of differentially expressed genes (DEGs) between Fibro_1 and a recent pan-fibroblast atlas^60^ revealed that the Fibro_1 cluster shared the most DEGs with the *Cxcl5*^+^ and *Adamdec1*^+^ clusters (**Figure S4Q**) found previously in settings of acute tissue injury or colitis^60^. Fibro_1 also expressed genes attributed to inflammatory cancer-associated fibroblasts (iCAFs) and antigen-presenting CAFs (apCAFs)^61,62^ (**Figure S4R**). Their emergence on day 1 post-wounding, across the wound diameter, representing more than 50% of all wound fibroblasts at that moment (**Figure 4C**) may be consistent with a general categorization of this program as being ‘acute inflammatory’.

The Fibro_2 cluster remained the steadiest in both space and time (**Figures 4C**) and Jaccard similarity analysis of DEGs with the pan-fibroblast atlas^60^ revealed high similarity to the ‘universal’ *Col15a1*^+^ fibroblast cluster (**Figure S4Q**).^59^

Cells belonging to the Fibro_3 and Fibro_4 clusters were predominantly found in inner diameter regions, where they make up about half of all fibroblasts present on Day 4 and 7, respectively (**Figure 4C**). DEGs from both populations were often associated with the extracellular matrix (ECM), such as *Tnfaip6* and *Timp1* ^63,64^ in Fibro_3 (**Figure S4P**) and *Col7a1, Col11a1, Postn*, as well as *Lrrc15*, in Fibro_4^64^ (**Figure S4P**). Gene expression similarity analysis revealed Fibro_4 as being most similar to those named *Lrrc15*^+^ in the pan-fibroblast atlas^60^ (**Figure S4Q**), which are possibly associated with immune exclusion in pancreatic cancer^65^. Both subpopulations also expressed *Cthrc1* (**Figure S4P**), which was recently described in collagen-producing lung fibroblasts^66^, further highlighting their phenotype as collagen-depositing cells during skin repair.

Finally, the Fibro_5 cluster represented a fibroblast subpopulation that constitutes 23.5% of all fibroblasts in unwounded skin but takes until day 7 and day 14 post-wounding to recover to this baseline level (**Figure 4C**). These expressed *Pi16* and *Dcn* (**Figure S4A**), markers associated with adventitial fibroblasts in the lung^66^ and dominated the fibroblast population outside of the wounded area in the 6 and 8 mm space, making up 55-77% of all fibroblast cells there (**Figure 4C**). Again, this highlights the effect of local wound repair on neighboring unwounded skin areas, where cell compositions are affected, despite the tissue not being subject to direct physical injury.

With the cell states of two key populations (fibroblasts and Mono_Macs) mapped across wound healing space-time (**Figures 2 and 4**), we began to study whether any of these co-occurred in space and time during the skin wound healing process, asking if this could reveal groups of heterotypic cell types putatively engaged in orchestrated and multicellular communication. We performed a Space-Time Correlation Analysis (STCA) by correlating the frequency of the fibroblast and Mono_Mac subpopulations across the tile plot (**Figure 4D**, see Methods for further details). This uncovered four co-occurring Mono_Mac/fibroblast pairs with very high degrees of space-time correlation, namely: Fibro_1 and Mono_Mac_1, Fibro_3 and Mono_Mac_3, Fibro_4 and Mono_Mac_4, Fibro_5 and Mono_Mac_MHCII_Mgl2 (**Figure 4E**). To visualize this in another way, we plotted cell percentages for each of these subpopulations over all samples and found high cross-correlations throughout the wound healing process (**Figure 4F**), with the Fibro_1 and Mono_Mac_1 pair coinciding as an ‘Early’ Wound Healing Cell Movement 1 (henceforth, WH-CM1), Fibro_3 and Mono_Mac_3 representing the ‘Int-In’ pattern (WH-CM2), Fibro_4 and Mono_Mac_4 representing the ‘Late-In’ pattern (WH-CM3), and Fibro_5 and Mono_Mac_MHCII_Mgl2 representing the ‘Late-Ex’ pattern (WH-CM4) (**Figures 2I and 4C**).

A larger STCA analysis, comparing all identified CD45^-^ non-immune and CD45^+^ immune cell subpopulations in our spatiotemporal scRNAseq dataset, demonstrated a 5th block of correlation (**Figures S4S and S4T**). This block, whose space-time distribution suggests it be called ‘Edge’ cell movement (WH-CM5) (**Figure S4S**), is only represented by endothelial cells, DCs, and NK cells. The distinct meanings for some of these additional cell types will await further in-depth analysis as, in this study, we next sought to dive yet deeper into the gene expression programs that might link two candidate cell types and underlie the pattern of their co-development.

### Gene program analysis identifies movements of gene expression across diverse cell types and predicts functional cell-cell interactions between macrophage and fibroblasts

In order to move beyond cluster-based analysis, which identifies the most significantly differentially expressed genes between clusters and not necessarily co-expressed genes, we adopted Non-negative matrix factorization (NMF). This analysis seeks to decompose a cell by gene matrix into the product of two smaller matrices with non-negative components ^35,67–70^ and can reveal layers of heterogeneity beyond clustering, especially in a population of cells without clearly demarcated subpopulations^35,69,35,67,69–71^. Here, collections of genes with similar expression patterns across the cell type contribute to a ‘factor’ with their gene weight denoting the strength of its contribution to that factor. Each cell is also then assigned a value for each factor (e.g. feature plots in **Figures 5A and 5B**). We henceforth use the terms ‘factor’ and ‘gene program’ interchangeably. In our study, these factors can be then visualized and studied based on mean factor levels per cell at each space-time coordinate (tile plots in **Figures 5A and 5B**).

**Figure 5.**
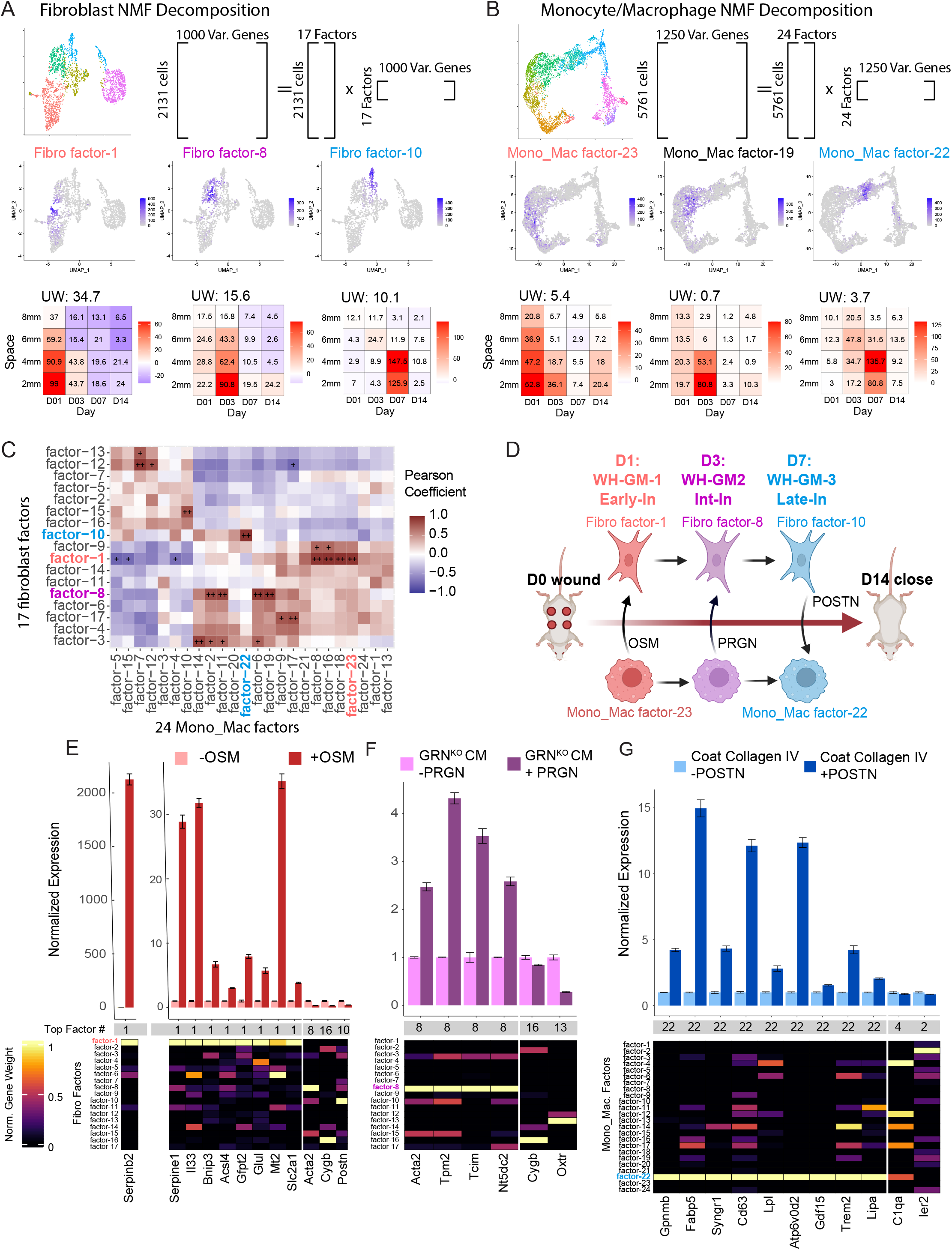
Gene program analysis identifies modules of gene expression across diverse cell types and predicts cell-cell interactions between macrophages and fibroblasts. (**A and B**) Schematic showing strategy for NMF based decomposition of the (**A**) fibroblast and (**B**) monocyte/macrophage populations. The top 1000 or 1250 most variable genes expressed in fibroblasts and monocyte/macrophages, respectively, within at least 2% of cells were used as the starting point for analysis. NMF decomposition of the fibroblast and Monocyte_Macrophage populations yielded 17 and 24 factors respectively, using cophenetic score to select the factor number (see Figure S5C). Shown are three example FeaturePlots for factor ‘expression’ and Tile Plots describing the average expression level of the factor as a function of space-time. (**C**) Space-time correlation matrix for average factor expression profiles. Correlation was calculated using Pearson correlation and significance adjusted for multiple comparisons using Benjamini-Hochberg procedure. + (alpha <0.05), ++ (alpha <0.005). (**D**) Cartoon schematic of hypothetical fibroblast-macrophage crosstalk and progression over the timespan of wound healing. Three putative interactions that we investigate *in vitro* are labeled. WH-GM, wound healing gene movement. OSM, oncostatin M. PRGN, progranulin. POSTN, periostin. Early-In, early interior. Int-In, intermediate interior. Late-In, late interior. (**E**) Primary mouse skin fibroblasts were treated with recombinant oncostatin M (OSM) at 20 ng/mL for 24 h before RNA extraction. Expression values of selected gene targets were assayed using RT-qPCR and normalized to *18s* levels. Error bars denote standard error of the mean from technical triplicates. Data is representative of two independent experiments. Factor with the largest contribution from each gene target denoted. Heatmap displays the normalized gene weight contribution to all 17 identified fibroblast factors for the gene being probed. (**F**) Conditioned 1% serum media from bone marrow-derived macrophages (BMDMs) isolated from granulin-knockout (GRN^KO^) mice was collected and added to primary murine skin fibroblasts (PSFs) with or without 1 ug/mL of recombinant Progranulin (PRGN). Following 24 h, cells were harvested for RT-qPCR for selected gene targets. Expression values of selected gene targets were assayed using RT-qPCR and normalized to *18s* levels. Error bars denote standard error of the mean from technical triplicates. Data is representative of two independent experiments. Factor with the largest contribution from each gene target denoted Heatmap displays the normalized gene weight contribution to all 17 identified fibroblast factors for the gene being probed. CM, conditioned media. (**G**) BMDM’s were plated on collagen IV-coated dishes with or without addition of 5 ug/mL recombinant POSTN in 1% serum media. Following 24 h, cells were harvested for RNA extraction. Expression values of selected gene targets were assayed using RT-qPCR and normalized to *18s* levels. Error bars denote standard error of the mean from technical triplicates. Data is representative of two independent experiments. Factor with the largest contribution from each gene target denoted. Heatmap displays the normalized gene weight contribution to all 24 identified M_M factors for the gene being probed.

For our study, we applied a variation of NMF analysis, termed non-smooth NMF (nsNMF)^71^ to our fibroblast and macrophage subset because of its sparser output during factorization as compared to the base NMF algorithm^72,73^. We also applied a cophenetic metric to identify the optimal number of factors for the most stable result as in **Figure S5C**^74^. This resulted in 24 factors for all Mono_Mac populations and 17 factors for the fibroblasts, each with a largely unique and specific collection of top-contributing genes (**Figures S5A and S5B**). We note that some genes are shared strongly between factors. For example, MHCII associated genes (e.g. *H2-Aa, H2-Ab1*) are shared between Mono_Mac factor-4 and factor-7 (**Figure S5A**), however such strong overlaps remain rare in our decompositions. In addition, top contributing genes to certain factors are also shared with DEGs that define clusters for example *Postn* is a DEG defining Fibro_4 cluster (**Figure S4P**) contributing to WH-CM4, and is also a weighted gene within fibroblast factor-10 (**Figure S5B**).

Similar to the STCA above with cell subpopulation fractions, we asked if the space time distribution of certain factors in the Mono_Mac population were correlated with factors in the fibroblast population. Thus, we constructed a correlation matrix between the profiles (similar approach to **Figure 4E**) for Fibroblast and Mono_Mac factors and identified pairs of strongly correlated factors. We focused on three sets of correlated factors, henceforth gene movements (GM), that center on D1, 3 and 7 of wound healing. These three, WH-GM-1 through 3, respectively exhibit space-time patterns broadly similar to WH-CM1 (‘Early’), WH-CM2 (‘Int-In’) and WH-CM-3 (‘Late-In)’ patterns described above (**Figure 5C and tile plots in Figures 5A and SB**).

Based on the space-time coincidence of these factors, we explored the hypothesis that such correlations could reveal cell-cell signaling between fibroblasts and Mono_Macs that drive the emergence of reciprocal gene programs over the time course of wound closure (**Figure 5D**). We began by examining fibroblast/Mono_Mac movements WH-GM-1 through 3 using CellChatDB^10^ to help identify putative ligands that are products of genes contributing to a ‘sender’ factor. We then experimentally queried if the set of top contributing genes of the correlated ‘target’ factor, in the opposite cell type, might be upregulated by the ligand of interest in the target cell type using a series of *in vitro* experiments. We explored three such relationships between macrophages and fibroblasts below:

#### WH-GM-1/Early-In

The Oncostatin M (OSM) pathway was predicted by CellChat to be most prevalent at D1 (**Figure S5D**) with *Osm* being a major contributing gene to Mono_Mac factor-23 (**Figure S5A**). That, in turn was defined to be in a space-time gene movement with fibroblast factor-1 (**Figure 5C**). We then used RT-qRT-PCR to test how *in vitro* stimulation with OSM would affect gene expression in primary skin fibroblasts (PSFs), finding that genes shown in **Figure S5B** that make up fibroblast factor-1 (*Serpinb2, Serpine1, Il33, Bnip, Gfpt2, Acsl4* among others) were significantly upregulated by OSM treatment (**Figure 5E**). Importantly, top contributing genes to other factors (e.g. *Acta2, Cygb, Postn*) (**Figure S5B**) were not similarly upregulated (**Figure 5E**).

#### WH-GM-2/Int-In

We noted the emergence of fibroblast factor-8 (e.g. *Acta2, Nt5dc2, Tpm2, Tcim*) (**Figure S5B**) within fibroblasts in the wound area on D3 (Int-In pattern) (**Figure 5A**). Previous literature demonstrated that Granulin (PRGN) stimulation could upregulate one of these genes, *Acta2*, in fibroblasts^75^. When we added GRN to fibroblasts, we found that upregulation of *Acta2* occurred and also coincided with upregulation of other top-contributing genes within the identified fibroblast factor-8 (e.g. *Tpm2, Nt5dc2* and *Tcim*) (**Figure 5F**). Because *Grn* does not specifically contribute to a single factor in the Mono_Mac population, it is still unknown how this fibroblast factor-8 is restricted to inner wound on D3; we offer some hypotheses in the discussion below.

#### WH-GM-3/Late-In

CellChat predicted the Periostin (POSTN) pathway for D3/7 could signal from fibroblasts to macrophages (**Figure S5D and S5E**) with predicted signaling from Fibro_3/4 to Mono_Mac_3/4, specifically through interaction with Integrins alpha-V and beta-3 (*Itgav, Itgb3)^76^. Postn* was a top contributing gene to fibroblast factor-10 (**Figure S5B**), which correlated well in space-time with Mono_Mac factor-22 to make up WH-GM-3 (**Figure 5C**). Among the strongest gene contributors to Mono_Mac factor-22 were *Gpnmb, Atp6v0d2, Fabp5, Cd63, Trem2*, and *Lipa* (**Figure S5A**). Using an *in vitro* stimulation of BMDMs, we confirmed that collagen-bound recombinant POSTN induced expression of top contributing members of Mono_Mac factor-22 but not those from other factors (e.g. factor 14 and 2: *C1qa* and *Ier2* respectively) (**Figure 5G**).

Similar to STCA analysis in **Figure S4T**, we correlated the space-time profiles of macrophage factors against the space-time frequencies of the other immune (**Figure S5F**) and non-immune (**Figure S5G**) types. In addition to the already described macrophage-fibroblast pairings, we found extended associations to other cell type frequencies, including association of Mono_Mac factor-4/5 to endothelial cell_3 frequency, Mono_Mac factor-7 to keratinocyte_4 frequency and Mono_Mac factor-14/22 to vascular smooth muscle 1 (vSM_1) frequency (**Figure S5G**). Finally, we extended the NMF approach to all cell types, identifying 114 gene programs scattered across our broad cell type definitions (top contributing genes and weights found in Supplementary Table 1); analysis of their correlated space-time profiles revealed ‘blocks’ of shared space-time patterns including but also extending beyond the Early, Edge, Int-In, Late-In, and Late-Ex patterns described above (**Figures S5H and S5I and Movie S5**). Investigation of the extensive collection of other correlated gene programs across cell types will await confirmation and investigation in a separate and larger study. Rather than defining more of these individually, we sought to understand whether the coincident macrophage-fibroblast patterns identified in wound healing might be conserved and detected in other settings, notably in tumors.

### Identification of conserved gene programs between wound healing and cancer

The paradigm of the tumor as a ‘wound that never heals’ has been put forth by several integrative studies^30,31^ and we sought to determine whether the tumor microenvironment, specifically the Mono_Mac compartment, could be described by factors identified in the wound healing process. We hypothesized that NMF analysis could reveal conserved gene programs between the wound healing and tumor tissue contexts.

We first generated an integrated scRNA-Seq dataset from two different mouse tumor models—B16F10 mouse melanoma and MC38 mouse colorectal—by sorting infiltrating immune cells 14 days after tumor injection (**Figure 6A**). We then examined the Mono_Mac subset similarly to our wound healing dataset, resulting in a 3,859 cell object (**Figure S6A**) and applied our ns-NMF workflow, identifying 25 factors (**Figures S6B and S6C**) We then sought to quantify the degree of factor similarity between the two contexts. Because the vast majority of genes contribute negligibly to a given factor, we decided against correlation metrices on the entire set of genes, to avoid overestimating similarities due to the heavy skewedness towards low values. Instead, we prioritized that a conserved pair of factors exhibits significant overlap in the top contributing genes; therefore, we applied a Jaccard distance metric based on the top 20 genes by weight or J_20_ distance. Using this metric, the vast majority of pairings displayed little to no overlap in their top 20 contributing genes, but a few rose prominently from the background (**Figure 6B**).

**Figure 6.**
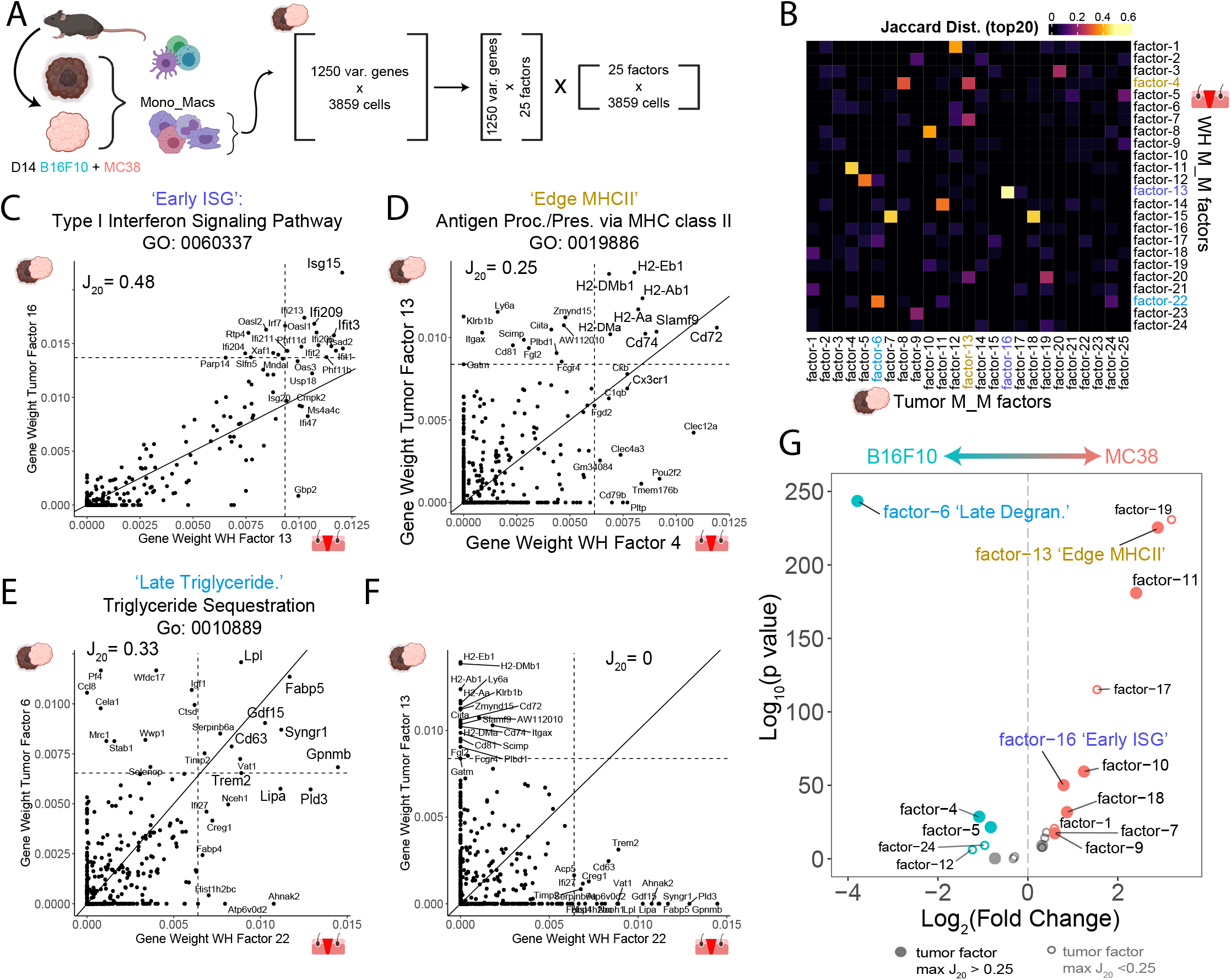
Identification of conserved gene programs in Mono_Mac between wound healing and cancer. (**A**) Strategy for generation of a multi-tumor model Mono_Mac scRNA-Seq dataset. B16F10 and MC38 tumors were harvested, sorted for CD45^+^ cells and used for scRNA-Seq. Following integration, Mono_Mac populations were selected for NMF decomposition, starting with 3859 cells and the top 1250 variable genes expressed in at least 2% of cells. This resulted in 25 factors of interest based on the cophenetic metric (seen in Figure S6C) (**B**) Heatmap showing the Jaccard_20_ distance (defined in Materials & Methods) between all 25 M_M tumor and 24 M_M WH factors based on top contributing gene weights. (**C-F**) Scatter plots for selected tumor/WH factor pairs for (**C**) Tumor factor-16 vs WH factor-13, (**D**) Tumor factor-13 vs WH factor-4, (**E**) Tumor factor-6 vs WH factor-22, and (**F**) Tumor factor-13 and WH factor-22 with the gene weight contributions plotted as calculated from the basis matrix in the NMF output (see Figure S5A for WH factors and S6B for tumor factors). Slope represents x=y line and dotted lines represent the weight for the 20^th^ highest gene contribution in either factor. The Jaccard_20_ index is shown and thus reflects the frequency of points in quadrant I over quadrants I,II and IV. For pairings in **C-E**, top shared genes in the upper right quadrant were put through Enrichr to find overrepresented cellular processes with the top result by p-value listed. Full Enrichr output can be found in the extended data (Supplementary Table 2). (**G**) Volcano plot showing differential expression of factors between MC38 and B16F10 Mono.Mac. datasets for the 25 identified factors. Y-axis denotes log10 of unadjusted p-value. Labelled points have adjusted p-value < 0.05 (Bonferroni correction) and absolute log2 fold-change greater than 0.5. Colored points have absolute log2 fold change greater than 0.5.

We generated scatterplots of gene weights to more closely examine shared gene contributions between pairs of WH and Tumor factors (examples in **Figures 6C-6F**). While the majority of factor pairings resembled **Figure 6F** with little to no overlap in gene contributions, we did identify nine strong pairings (J_20_>0.25) (**Figure 6C-6E** and **Figures S6D-S6I**). Gene Ontology (GO) analysis on the shared genes in the upper right quadrant revealed unique cellular processes associated with each pairing which could, at least partially, reflect the functional output of those gene programs (full list of GO terms in Supplementary Table 2). We focus on three of these pairings below:

#### ’Early ISG’ Program

Of the stronger pairings, one, the WH M_M-13-to-Tumor M_M-16 factors was highly characterized by a collection of well-described Interferon stimulated genes^77^ (ISG) (**Figure 6C**). GO analysis of the top shared genes yielded the term ‘Type I interferon signaling pathway’. In the wound, this WH M/M factor-13 followed an ‘Early’ WH-GM1 spacetime profile (**Figure S5H and S5I**).

#### ’Edge MHCII program

Another factor pairing, (WH M_M-4-to-Tumor M_M-13; **Figure 6D**), involved genes associated with antigen presentation through MHCII. In this case, there were highly correlated expression levels of *H2-Ab1, H2-Eb1* and other genes associated with antigen presentation (e.g. *Cd74*). In this case, there were substantial additional differential genes expressed in the wound healing versus tumor factors, and these may represent context-specific genes that co-express only in those specific setting. GO analysis of the top shared genes indicated antigen processing and presentation via MHCII. WH M_M factor-4 followed an ‘Edge’ WH-GM5 space time profile which seemed to follow the closure of the wound (**Figure S5H and S5I**).

#### ‘Late Triglyercide’

The WH M_M factor-22 to tumor M_M factor-6 pairing, was marked by genes including *Gpnmb, Fabp5, Syngr1, Cd63, Trem2*, and *Lipa* among others (**Figure 6E**). Several of these genes are associated with lipid trafficking, including *Trem2, Fabp5*, and *Lipa*. GO analysis revealed an enrichment for ‘Triglyceride Sequestration’ suggesting a functional output associated with intracellular vesicle trafficking and exocytosis. This particular factor also followed the space time pattern we termed ‘Late-In’ or WH-GM3 (**Figure S5H and S5I**).

Finally, given these few but significant similarities, we sought to determine whether such programs are used equivalently in the two model tumor systems. The MC38 dataset displayed significant enrichment of factors including tumor factors 11 and 13 which had strong (J_20_>0.25) correspondence to a WH factor. Meanwhile, B16F10 tumors were uniquely marked by a separate and very strong enrichment of factor-6, corresponding to WH factor-22 (**Figure 6G**). Other conserved factors such as tumor factor-16 (‘Early ISG’) displayed less dramatic enrichment in either model (<2 fold). Together this suggests that while tumors may indeed borrow factors from wound healing, individual tumors may do so uniquely.

### Application of Movements: Conservation of Mono_Mac factors predicts increased Periostin and decreased Selectin-P^+^ vessel density in B16F10 vs. MC38 tumor models

We finally sought to study the conserved gene programs spatially and confirm the differential usage of wound healing factors in different tumor models. To this end, we used our M_M factor similarity /translation matrix (**Figure 6B**) and our wound healing movement identification, to make and then test predictions about the state of the tumor microenvironment in either tumor model. As observed in **Figure 6G**, the ‘Late Triglyceride’ pairing (M_M tumor factor-6 corresponding to M_M WH factor-22 **Figure 6E**) was more highly expressed in B16F10. We used the observation that the latter factor was paired in a space-time movement with fibroblast factor-10 in our wound healing dataset (**Figure 5C**), to form the prediction that one of the members of fibroblast factor-10, POSTN, would be more prevalent in B16F10 vs. MC38 model (**Figure 7A**). Immunofluorescence staining of both tumors identified indeed profoundly larger density of POSTN fibers within B16F10 tumors as compared to MC38 (**Figures 7B and 7C**). In addition, a considerably larger fraction of CD11b^+^ cells were in close contact with POSTN fibers in the B16F10 model vs. MC38 (**Figures 7D, 7E, and S7A**).

**Figure 7.**
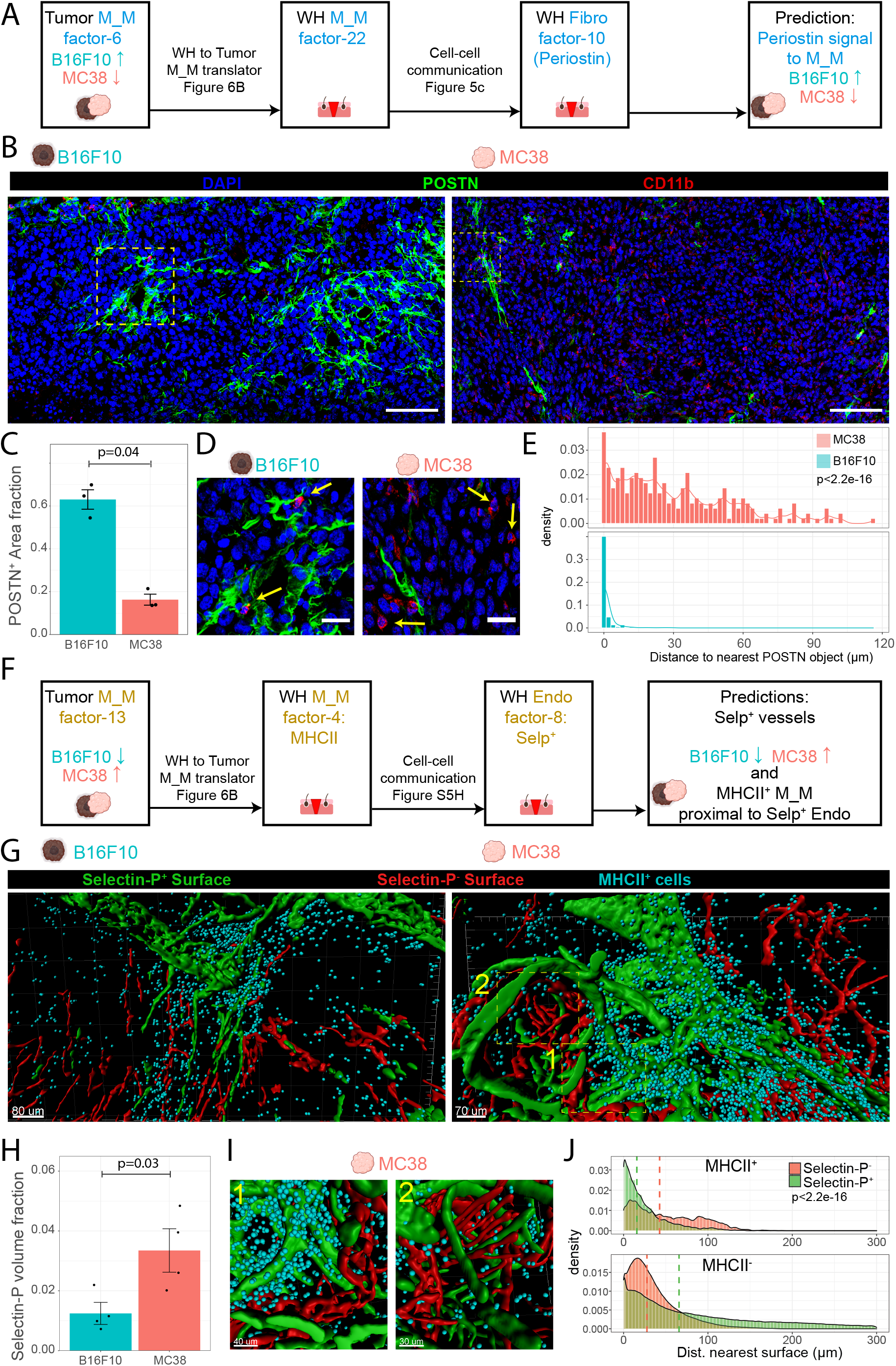
Conservation of Mono_Mac factors predicts increased Periostin and decreased Selectin-P^+^ vessel density in B16F10 vs MC38 tumor models. (**A**) Schematic for hypothesis generation in the tumor setting. Translation of the tumor M_M factor-6 to the WH M_M factor-22 allows prediction that the same stimuli (Periostin from WH Fibro factor-10) might underlie tumor M_M factor-6 and thus be more prevalent in the B16F10 vs. MC38 tumor model. (**B**) Representative immunofluorescent images of 10 μm fresh frozen sections of D14 B16F10 and MC38 tumors stained for DAPI (blue), POSTN (green), and CD11b (red). Scale bar = 100 μm. Representative of two experimental replicates. (**C**) Barchart denoting fraction area of POSTN^+^ surfaces as a fraction of total imaged tissue area. Onesided Wilcoxon rank-sum test used. Each point represents a scanned area from 3 separate tumors. (**D**) Insets from denoted regions of interest in Figure 7B. Arrows denote CD11b^+^ cells in close contact with POSTN^+^ surfaces (B16F10) or not (MC38). Scale bar = 25 μm. (**E**) Histograms showing distribution of distance of CD11b^+^ cells to nearest POSTN surface. Representative of 3 independent replicates (3 tumors). Bin-width = 2 μm. Kolmogorov-Smirnov test used to test against null hypothesis that samples were drawn from the same distribution (**F**) Schematic for hypothesis generation in the tumor setting. Translation of the tumor M_M factor-13 to the WH M_M factor-4 allows prediction that the same association between WH M_M factor-4 with endothelial WH factor-8 might be found in the tumor setting and that endothelial factor-8 might be more prevalent in MC38 vs. B16F10 tumor model (**G**) Processed 3D images of cleared 250 μm thick tumor slices from a (left) B16F10 and (right) MC38 tumor. Generated surfaces based on Selectin-P^+^ staining (green) and CD31^+^Selectin-P^-^ signal (red). Dots (cyan) denote MHCII^+^ cells. (**H**) Bar chart showing comparison of the cumulative Selectin-P^+^ surfaces volume normalized to total imaged tissue volume between MC38 and B16F10. One-sided Wilcoxon rank sum test used. (**I**) Zoomed in and rotated insets from MC38 tumor in Figure 7G exemplifying (1) dense accumulation of MHCII^+^ cells proximal to Selectin-P+ vessels and (2) sparse MHCII^+^ cell accumulation proximal to Selectin-P^-^. (**J**) Histograms indicating the distances of MHCII^+^ spots and MHCII^-^ spots to the nearest Selectin-P^+^ or Selectin-P^-^ surface in the MC38 model. Dashed line indicates the median. Histograms representative of 4 independent replicates (4 separate tumors). P-value calculated via Kolmogorov-Smirnov test.

Conversely, the shared ‘Edge MHCII’ factor (tumor M_M factor-13 corresponding to WH M_M factor-4 **Figure 6D**) was more dominant in MC38 (**Figure 6G**). Going back to the space-time correlations in **Figure S5H**, WH M_M factor-4, as an ‘Edge’ pattern WH-GM5, grouped with WH endothelial factor-8, which comprised genes including *Selp, Vwf, Sele* and *Ackr1*. Following a similar line of inference as before with POSTN signaling to macrophages, we predicted a higher density of Selectin-P^+^ vasculature in MC38 vs. B16F10 tumors (**Figure 7F**).

Using 3D imaging of cleared thick tumor sections (250 μm) (**Figures 7G, S7B**), we found a markedly increased density of Selectin-P^+^ vasculature in the MC38 tumor relative to B16F10 tumor (**Figure 7H**), consistent with our prediction in **Figure 7F**. Interestingly, analysis also revealed clear physical proximity of MHCII^+^ cells (cyan points) with Selectin-P^+^ vasculature (green surfaces) vs Selectin-P^-^ vasculature (red surfaces) in both tumor models (**Figures 7I, 7J, and S7C**), with dense accumulations of MHCII^+^ cells ringing Selectin-P^+^ vessels. We also note this preference for Selectin-P vessels was not found in MHCII^-^ cells (**Figure 7J, and S7C**). These lines of evidence provide examples of how the conceptual framework of conserved gene programs and multicellular movements can inform hypothesis generation spanning tissue contexts.

## Discussion

Recent advances in technology have allowed single cell transcriptional profiling of diverse tissues, leading to the identification of a plethora of cell types and subtypes in physiological and pathological tissue contexts. How those cell types are spatially and temporally organized within the tissue will help us understand the underlying dynamic nature of tissue homeostasis and pathology. Here, we established a high-dimensional spatiotemporal framework to study pairing of cell types during a physiologically complex process-namely during wound repair. In this setting, the concept that spatiotemporal correlation may indicated paired biology drove the identification of groups of cell types that together form larger cellular movements, wherein their emergence and/or disappearance is correlated and partially conserved in both skin repair and tumor growth.

Prior to our study several fibroblast-macrophage interactions have been described in health and disease, such as fibrosis and cancer^23^. The use of space time correlation analysis (STCA) takes this systematically one step further and identified four distinct fibroblast-macrophage cellular pairings during wound repair (**Figure 4F**), each characterized by a distinct space-time pattern during the repair process. The earliest pattern consisted of inflammatory Fibro_1, expressing the neutrophil-attracting chemokine *Cxcl5*^78^ and the early inflammatory gene *Ptx3*^79^. Accordingly, this ‘Early’ pattern was accompanied by neutrophils and monocytes (**Figure S4T**) whose accumulation in early wounds had been previously identified^3^ but not tied to this fibroblast population nor the co-appearance of an endothelial cell population (endothelial cell 2), highlighting the involvement of non-immune cells in early wound repair. In later stages, the ‘Late-Ex’ pattern of Fibro_5 and Mono_Mac_MHCII_Mgl2 cells was accompanied by several other cell subtypes, such as mast cells, MHCII^hi^ Mono_Macs, T cells, and keratinocyte subsets. Together, this provides an example of a multicellular pattern composed of a diverse range of cell identities, akin to the recent finding of interacting stellate cells, hepatocytes, endothelial cells, and liver-infiltrating monocytes together coordinating a local niche for monocyte-to-macrophage differentiation^6^. These observations highlight the importance of considering cell function and differentiation in the context of their subtissular location and surrounding cell environment.

By combining NMF-based decomposition with spatiotemporal data, we moved beyond conventional cell subset identification, and identified co-occurring gene programs that provide candidates for reciprocal regulation between cell types. We present this approach as a framework for identifying cell-cell communication pathways and their downstream effects on gene expression and as a means to identify pathways to target in order to advance transitions in biological systems. In our conceptual framework, factors or gene programs represent a functional module activated in a cell type due to response to external stimuli which in turn may be derived from factors activated in other cell types. In this way our work frames wound healing as coordinated, interlinked gene programs (‘movements’) across diverse cell types. In particular, we put forth a model of reciprocally acting fibroblast and macrophage gene programs, whereby emergence of gene programs in one cell type ratchets the phenotype of another cell type, in turn driving a new program’s emergence (**Figure 5D**). One example we highlight is Oncostatin M, previously known to induce *Serpine1* and *Il33* upregulation^80,81^. Using NMF analysis combined with spatiotemporal correlations across cell types, we identified a host of other previously undescribed genes (e.g. *Glul, Mt2, Acsl4*) in fibroblasts upregulated by Oncostatin M.

Using gene program analysis, we were also able to predict the suite of genes induced by progranulin (PRGN) treatment on fibroblasts, going from the known genes (*Acta2*) to the unknown (*Tpm2, Tcim, Nt5dc2*)^75,82^. In our dataset, Mono_Macs are the primary cell type expressing *Grn*, but given that granulin is expressed throughout the Mono_Mac population and does not contribute to any identified factor specifically, it remains unclear how its effects are restricted to the Int-In D3 pattern. Possibly the presence of another signal—perhaps a protease that cleaves progranulin into individual active peptides ^83^—licenses granulin’s effects on fibroblasts or earlier signals before day 3 prime fibroblasts to respond to granulin signaling. ^83^

Similarly, our movement identification suggested periostin, which emerges later in wound healing in fibroblasts, as a candidate to induce the genes defined in factor-22 in Mono_Macs. This factor includes *Gpnmb*, *Lipa*, and *Trem2*. Previous reports have mainly described the ability of periostin to promote adhesion and migration of macrophages in tumor settings^84,85^. Given the therapeutic implications of *Trem2* expression in tumor associated macrophages, the identification of a possible molecular etiology is of significant interest for tumor immunology^86–88^. Notably, we observe from tissue staining that periostin is found in fibrillar structures in the tumor, consistent with its description as a matricellular protein^89^. Although we verified a handful of cell-cell interactions here, our dataset highlights a multitude of movements and other gene program pairings, serving as a potential resource to other investigators seeking to infer cell-cell crosstalk mechanisms. Finally, although we focused on verification of fibroblast-macrophage interactions, we generalized our computational approach to all other identified cell types, immune and non-immune. Doing so allowed us to identify the major groupings of spatiotemporally correlated gene programs in cell types as disparate as neutrophils, endothelial cells, and keratinocytes, with each grouping displaying a unique space-time pattern (**Figures S5H, S5I, and Movie S5**). Similarly, to the ‘hubs’ of gene programs described in Pelka et al. these groupings of gene programs could represent spatially co-localized, functional units of cell organization in tissue^69^. Additionally, such groupings could underlie the findings that tumor microenvironments tend to adopt defined compositional ‘Archetypes’^90^, wherein each archetype might draw upon these groups of gene programs differentially. Keeping in mind the caveat of mismatch between mRNA and protein levels, the gene programs identified here, and their shared spatiotemporal profiles, will inform future studies to identify which correlations are indicative of true causation through cell-cell crosstalk^91^; indeed combination with curated databases such as NicheNet’s weighted networks between ligand and target genes could allow for tracing back the signaling events that activate a given gene program^12^.

Moving beyond skin wound healing, we posit that gene program analysis can serve as a powerful tool as we move towards integrative studies across tissue types and disease models. For example, NMF analysis was recently employed to identify shared gene programs associated with cell cycle, EMT, and senescence across many cancer cell lines^35^. We identified paired gene programs in the Mono_Mac populations from two tumor models and our wound healing dataset. We also note that factor pairings are not perfect, with some genes showing little contribution in one setting versus major contribution in another. We theorize this could indicate either a purely coincidental co-expression in one setting or an additional layer of epigenetic regulation. High dropout rates of certain genes in one dataset versus another due to differential sequencing depth could also be a cause. We also note the presence of many factors without a good counterpart. These factors could represent artifacts introduced in the processing pipeline such as a collection of housekeeping genes moving together in cells with lower UMI counts or could be representative of truly tissue specific programs. This methodology can only be strengthened by application to ever more diverse datasets to extract gene programs with representation across contexts. For example, during fibrosis whereby excessive ECM deposition begins to adversely affect organ function^92^, many of the same cell-cell interactions that drive regeneration become dysregulated, e.g. persisting inappropriately^23,93–95^. Thus, identifying the space-time profiles of conserved gene programs and between fibroses, wound healing, and tumorigenesis could identify where the dynamics of pathologies begin to diverge in state space from normal^31^.

As a proof of concept for this approach, we demonstrate that testable hypotheses can be generated from our correlative studies by proceeding in a two-step fashion, first translating tumor Mono/Mac factors to wound healing Mono/Mac factors (**Figure 6B**), then using identified multicellular movements to translate WH M/M factors to correlated WH factors in other cell types (**Figure S5H**). This analysis allowed us to predict increased periostin signaling to macrophages in B16F10 versus MC38 tumors and conversely, an enrichment for P-selectin-expressing endothelial cells (endo factor-8) in MC38 versus B16F10 tumor. The close association of MHCII^Hi^ macrophages with CD31 ^+^ vasculature has been reported previously^96^ but not specifically Selectin-P^+^ vasculature. Of interest, the endothelial factor-8 genes including *Selp, Sele, and Ackr1* also define a tumor associated high endothelial venule (TA-HEV) network found to be correlated with patient response to checkpoint blockade immunotherapy^97^. Given the relative unresponsiveness of the B16-F10 melanoma model versus the MC38 model^90^, we posit that the difference in density of endothelial cells expressing endo factor-8 results in functional differences in the state of the immune compartment. We anticipate these translation tables can be generalized to other tumor models and tissue pathologies such as fibrosis, which will allow to identify thus far unappreciated cell-cell interactions in those settings.

## Limitations of study

Our scRNAseq study identified several immune and non-immune cells and their dynamic change during the skin repair process. However, we noticed no capturing of adipocytes, neurons, or glial cells. Presumably the combination of our enrichment method of cells via fluorescent-activated cell sorting and the subsequent scRNAseq pipeline using the 10x Genomics platform led to loss of these and potentially other unidentified cell types. This precludes analysis of their potential involvement the presented cell patterns and gene factor networks. Previous studies have highlighted the importance of adipocytes in skin repair^42^, as well as the role of macrophage-neuron crosstalk skin homeostasis^98^.

The skin is colonized by a microbiome that breaches skin barrier upon injury. Members of the microbiome are sensed by local immune cells, initiating local immune responses. In carcinomas originating in barrier tissue, especially along the gastrointestinal tract (mouth, throat, esophageal, gastric, colon, rectal), skin when ulcerating, and lung, similar local immune responses can unfold upon barrier-breaching. But in an otherwise sterile environment during tumorigenesis, pathogen sensing would be less likely to influence cell-cell communication. Thus, commonalities discovered here using subcutaneous tumor models and skin wounding are conserved in both absence and presence of pathogen detection. This also raises the question of how other sources of internal organ damage, such as chemicals, influence crosstalk between immune and non-immune cells and how conserved these layers of communication are in both mice and humans.

## Acknowledgements

We would like to thank Drs. Hong-Erh Liang and Richard Locksley for the generous gift of the Arg1-reporter mouse. We also thank Drs. Chris McGinnis and Zev Gartner for the LMO’s and barcoded oligonucleotides and comments on manuscript. Additionally, we would like to thank members of the Krummel lab for scientific discussion and comments on the manuscript. This work was supported by funds from NIH R01CA197363. K.H.H. is supported by the American Cancer Society Postdoctoral Fellowship (#133078-PF-19-222-01-LIB) and N.F.K. is supported by NIH T32 5T32CA108462-17. Flow cytometry was performed at the UCSF Parnassus Flow CoLab, RRID:SCR_018206. We also thank the Center for Advanced Technology at UCSF for sequencing support.

## Author Contributions

Conceptualization, K.H.H., N.F.K. and M.F.K.; Methodology, K.H.H. and N.F.K.; Investigation, K.H.H., N.F.K. and T.C.; Formal Analysis, K.H.H., N.F.K. and T.C.; Visualization, K.H.H., N.F.K.; Writing – Original Draft, K.H.H. and N.F.K.; Writing – Review & Editing, K.H.H., N.F.K., and M.F.K.; Funding Acquisition, M.F.K.; Resources, M.F.K.; Supervision, M.F.K., K.H.H.

## Data Availability

Raw Fastq files and count matrices from scRNAseq experiment are available at NCBI GEO with the following accession number: GSE204777.

