## Supplementary material for "Space-Time Mapping Identifies Concerted Multicellular Patterns and Gene Programs in Healing Wounds and their Conservation in Cancers": Methods

**Animals**
All mice were housed in an American Association for the Accreditation of Laboratory Animal Care (AALAC)-accredited animal facility and maintained in specific pathogen-free conditions. All animal experiments were approved and performed in accordance with the Institutional Animal Care and Use Program protocol number AN184232. Wild-type female C57BL/6 mice between 8-10 weeks old were purchased from The Jackson Laboratory, *Arg1-tdTomato$^{CreERT2}$* x *R26R-EYFP* mice were a kind gift from Drs. Hong-Erh Liang and Richard Locksley (UCSF). All mice were housed at the University of California, San Francisco (UCSF) animal facility with typical light/dark cycles and standard chow. For tumor growth studies, MC38 colon cancer ($5x10^5$ cells / 50 µl) or B16-F10 melanoma cancer cells ($1x10^5$ cells / 50 ul) were transplanted into the subcutaneous region of the mouse flank. On day 14 after tumor challenge, when tumors reached a size/volume of approximately 0.5 $cm^3$, mice were sacrificed, tumors were excised and processed for downstream analysis.

**Cell lines**
B16-F10 and MC38 cells were purchased from ATCC and cultured at 37ºC in 5% $CO_2$ in DMEM supplemented with 10% FCS (Benchmark), 2 mM L-glutamine, 100 U/ml penicillin, and 100 µg/ml streptomycin.

**Full-thickness wounding**
All mice used for wounding experiments were between 7-10 weeks old (second telogen hair follicle phase). Two to four days before wounding, back skin was shaved and residual hair was removed using NAIR (Walgreens). On the day of wounding, mice were anesthetized with 3% isofluorane and subcutaneously injected with 50 µl of 0.25% bupivacaine and 50 µl of 50 µg/ml buprenorphine for analgesia. The back skin was then sterilized using a Betadine Solution swab stick (for scRNAseq and imaging experiments) or ChloraPrep (BD Biosciences) (for Mass Cytometry experiments to avoid iodine contamination during sample acquisition). Four full-thickness wounds were generated with a 4 mm biopsy punch. Wound diameter was measured with calipers.

**Tissue Processing**
At time of analysis, mice were euthanized, back skin was dissected from mice, excess scapular and inguinal fat was removed, and wounds plus adjacent tissue was excised using an 8 mm biopsy punch. An 8 mm biopsy punch was used to collect unwounded back skin as a day 0 or unwounded control. Four to six 8 mm biopsy punches were pooled from unwounded back skin to collect sufficient cells for downstream analysis. On ice, tissue was finely minced with scissors and then placed in a 2 ml tube containing 1 ml of digestion medium (2 mg/ml collagenase XI, 0.5 mg/ml hyaluronidase, 0.1 mg/ml DNase in RPMI with 10% FCS, 2 mM L-glutamine, 100 U/ml penicillin, 100 µg/ml streptomycin, 50 µM beta-mercaptoethanol). The tube was placed horizontally in a bacterial shaker for 45 min at 37ºC and 225 rpm. The sample was then filtered through a 100 µm filter and washed with 10 ml cold RPMI. The generated single cell suspension was then processed for further depending on analysis method.

**Mass Cytometry**
*Antibody conjugate generation*
All mass cytometry antibodies are listed in the Key Resource Table. Primary conjugates of mass cytometry antibodies were prepared using the MaxPAR antibody conjugation kit (Fluidigm) according to the manufacturer's instructions. After labeling, antibodies were diluted in Candor PBS Antibody Stabilization solution (Candor Bioscience GmbH, Wangen, Germany) containing 0.02% NaN3 at 0.1-0.3 mg/ml. Antibody conjugates were stored at 4ºC. Each antibody clone and lot was titrated to optimal staining concentrations.

*Cell Preparation*
After single cell generation, cells were resuspended at $1x10^6$ cells / ml PBS with 5 mM EDTA. An equal volume of PBS with 5 mM EDTA plus 50 µM Cisplatin (Enzo Life Sciences, Farmingdale, NY) was added and incubated for 60 s before quenching with an equal volume of PBS with 5 mM EDTA plus 0.5% BSA. Cells were centrifuged at 500 g for 5 min at 4ºC. Washed cells were resuspended in PBS with 5 mM EDTA and fixed with 2.67% PFA for 10 min at RT. Fixation was quenched by adding 10x volume of PBS with 5 mM EDTA plus 0.5% BSA. Cells were centrifuged at 500 g for 5 min at 4ºC and resuspended in PBS with 0.5% BSA and 10% DMSO before frozen and stored at -80ºC until barcoding.
*Mass-Tag Cellular Barcoding*
After thawing stored samples at RT, up to $1x10^6$ cells from each mouse were barcoded with distinct combinations of stable Pd isotopes in PBS with 0.02% saponin as described before[1]. After incubation at RT for 15 min on a shaker at 90 rpm, cells were pelleted at 600 g for 5 min at 4ºC and washed two more times with cell staining media (CSM: PBS with 0.5% BSA and 0.02% NaN3). After the last wash, all samples were pooled into a single 15 ml tube.
*Mass Cytometry Staining and Acquisition*
After barcoding and pooling of samples, all cells were pelleted at 600 g for 5 min at 4ºC and resuspended in CSM containing a metal-labeled anti-CD16/32 antibody for 5 min at RT on shaker at 90 rpm to block Fc receptors. Extracellular cell markers were stained by adding a master mix of metal-labeled antibodies listed in the Key Resource Table. After incubation for 30 min at RT on a shaker at 90 rpm, cells were washed with 5 ml of CSM. For cell permeabilization prior to intracellular stain, pelleted cells were resuspended in 1 ml of pre-chilled 99% MeOH (Sigma Aldrich) and incubated for 10 min at 4ºC. Cells were washed twice in 5 ml of CSM prior to intracellular stain. For intracellular staining, pelleted cells were resuspended in the master mix of metal-labeled antibodies for intracellular targets and incubated for 30 min at RT on a shaker 90 rpm. Cells were washed with 5 ml of CSM and then resuspended in 2 ml PBS with 1.6% PFA and 0.55 µl 191/193Ir DNA Intercalator (Fluidigm). Cells were incubated overnight at 4ºC. Cells were then washed once with 12 ml CSM, once with 12 ml PBS, and once with 12 ml of $ddH_2O$ prior to dilution in $H_2O$ at about $3x10^6$ cells/ml containing normalization beads (see below), filtered through a 70 µm cell strainer and then analyzed on a Helios mass cytometer (Fluidigm, San Francisco, CA). We analyzed $3x10^4$-$2x10^5$ cells per sample.
*Mass Cytometry Bead Standard Data Normalization*
Data normalization was performed as previously described[2]. All mass cytometry files were normalized together using the mass cytometry data normalization algorithm[3], which uses the intensity values of a sliding window of the bead standards to correct for instrument fluctuations over time and between samples.
*Data Analysis*
After bead normalization and debarcoding using the Premessa package (https://github.com/ParkerICI/premessa), singlets were gated by Event Length and DNA content. Live cells were identified as Cisplatin-negative cells. Immune cells were identified as $CD45^+$ cells and downsampled to 3.000-10.000 cells per sample prior to export and concatenation in FlowJo (Treestar). After concatenation of all cells from each timepoint, cell clusters were identified using the Phenograph algorithm[4] using k=190 nearest neighbors. Mean intensity values of each marker per cluster were exported from FlowJo and imported into Morpheus (https://software.broadinstitute.org/morpheus/) to generate heatmaps. The identity of each cluster was determined based on expression of stained markers.

**Single cell RNA sequencing of skin wound-associated cells**
For spatiotemporal scRNAseq analysis, skin wounds were processed as described above to generate single cell suspensions, except that after excising the skin wound and adjacent tissue with an 8 mm biopsy punch, the excised skin wounds were further partitioned by successively

using a 2 mm, 4 mm, and 6 mm biopsy punch. Digestion of tissue proceeded as described above, except that no DNase I was used in the digestion media to avoid potential downstream cleaving of nucleotide barcodes. After washing and pelleting of cells at 500 g for 5 min at 4, samples were each resuspended in 100 µl of staining buffer (PBS plus 2% FCS and 2 mM EDTA) and 2 µl of purified anti-mouse CD16/32 (Fc Shield, clone 2.4G2, Tonbo Biosciences) was added to the sample. Samples were incubated for 15' on ice to block FC receptors. Following this incubation, anti-mouse CD45 Alexa Fluor 647 (clone 30-F11, Biolegend, 1:1000) was added and cells were incubated for an additional 20' on ice. Cells were washed using staining buffer, pelleted at 500 g for 5 min, and resuspended in staining buffer plus DAPI (1 µM). Cells were filtered on a 40 µm cell strainer right before being sorted on a BD FACSAria II cell sorter (BD Biosciences). $CD45^+$ and $CD45^-$ cells were collected separately in ice cold collection buffer (colorless RPMI supplemented with 10% FCS, 2 mM L-glutamine, 100 U/ml penicillin, 100 µg/ml streptomycin, 50 µM beta-mercaptoethanol). Across all 17 samples (= 4 timepoints multiplied by 4 areas plus one unwounded sample), 206 x $10^3$ $CD45^+$ and 263.7 x $10^3$ $CD45^-$ cells were collected. Cells of individual samples were pelleted at 500 g for 5 min at 4ºC. Supernatant was removed and cells were resuspended in 160 µl of colorless RPMI. Samples were then barcoded using lipid modified oligonucleotides (LMO) as in McGinnis et al.[5] before pooling. The pool was then split over 5 lanes of a 10X 3' NEXTGEM chip before encapsulation with a targeted cell number of 12,000 per lane. Following the MULTI-Seq library construction protocol, additive primer was spiked in during the cDNA amplification step as described in McGinnis et al[5], the supernatant was reserved following SPRI cleanup and separately amplified. Finally, libraries were pooled for sequencing using a 1:10 molar ratio of LMO : gene expression (GEX) libraries with approximately equal representation from each lane. Pooled samples were then sequenced using 1 lane of a S4 flowcell for a target of ~3B reads. GEX libraries had total read counts of (493M, 515M, 495M, 452M, 403M) for lanes (1,2,3,4,5) respectively, showing similar order of magnitude representation in read counts. The estimated sequencing saturation ranged from 58-64% (calculated via CellRanger), indicating sequencing depth did not vary appreciably between lanes. LMO libraries had total read counts of (37M, 39M, 38M, 36M, 32M) also showing stable representation from all 5 lanes.
Each individual well was demultiplexed using deMULTIplex package[5] to remove doublets and unlabeled cells. Confidently hashed cells were then carried forward for integration in Seurat v3. From there, cells with percent mitochondrial reads >25 percent and number of genes <200 were filtered out. Following an initial high-level clustering, we removed several clusters composed mostly of high mitochondrial % cells or low nUMI and a immune/non-immune doublet cluster. An initial high-level clustering and dimensional reduction was used to define the CD45+ and CD45- subsets. Once subsetted, each group was then further re-clustered to generate the 'final' CD45+/- datasets.

**Candidate identification of ligand-receptor interactions using CellChat**

For CellChat analysis[6], the M/M subset and fibroblast subsets were merged into a single Seurat object, then split into 5 Seurat objects based on the 'Day' metadata. Each Seurat object was then imported into a CellChat object using the 'RNA' assay. Signaling network likelihoods were calculated using CellChat's computeCommunProb function using the trimean method, using raw data, and with population size scaling. Stacked barchart was calculated using the rankNet function with genes ordered by day weighted average.

**Embedding a low-dimensional representation of samples using PhEMD**

For mass cytometry data, PhEMD was employed to generate a two- or three-dimensional embedding of all samples split by timepoint based on their immune cell profiles[7]. For scRNAseq data, a two-dimensional embedding of $CD45^+$ samples split by all space/timepoints based on their immune cell profiles was generated. Briefly, PhEMD first generates a reference map of cell subtypes, then uses Earth Mover's Distance (EMD) to compute pairwise dissimilarities between samples (incorporating sample-to-sample differences in cell fractions of each cell subtype as well as intrinsic dissimilarities between subtypes based on the cell subtype reference map), and finally applies a dimensionality reduction technique to the sample-to-sample distance matrix to generate a final embedding of samples. The Seurat implementation of 3D Uniform Manifold Approximation and Projection (UMAP) was used to map the cell-subtype space. Dissimilarity between each pair of cell subtypes was defined as the distance between the centroids (in UMAP space) of all cells assigned to the two respective subtypes. PHATE was applied to the EMD patient-to-patient distance matrix to generate the final 3D embedding of samples[8].

For PhEMD application to CyTOF data, the fully concatenated clean datasets were converted into feature-cell sparse matrices that Seurat could import, analogous to a scRNA-Seq dataset. The data was then transformed using the arcsinh transform and processed as described above.

**NMF Decomposition**

The CD45-/+ objects were subsetted according to broad cell type definitions (M/M, Neutrophils, DC's, Mast Cells, T, TNK, B and Fibroblast, Keratinocyte, Endothelial, Melanocyte, Dermal sheath papilla, vSM). The top 1250 most variable genes (depending on diversity of subset expression patterns) were selected using the 'vst' method in Seurat. Using the scaled RNA assay (non-centered) and subsetting out the most variable genes, we thus created a 1250xN (N = # of cells) expression matrix, then applied the non-smooth NMF algorithm (as described in NMF package[9]). We used a parameter sweep combined with the cophenetic metric to find the most stable number of factors using 50 iterations.

For gene weights per factor, we used the basis matrix output from the nsNMF and sorted based on the highest contributing genes. For plotting purposes, we normalized gene contributions across all factors to understand how specific a given gene was for a factor. We used the coefficient matrix as the 'expression' value for each cell for that factor. This value was used directly for average factor expression and feature plots.

When translating factors between tumor and wound healing, we used a Jaccard distance metric based on the top 20 contributing genes for each factor.

$$J_{20} = ((top\ 20\ genes\ factor\ n) \cap (top\ 20\ genes\ factor\ m)) / (top\ 20\ genes\ factor\ n \cup top\ 20\ genes\ factor\ m)$$

**Space/Time Tileplot of cell frequencies**

To depict the distribution of identified cell subsets in the scRNAseq data set within the wound over time, we devised a Space/Time Tileplot that is a 4x4 grid, where the x-axis is split into the four time points of sampling (day 1, day 3, day 7, and day 14) and the y-axis is split into the four areas of the sampled wound (2 mm wound center, 4 mm ring, 6 mm ring, 8 mm ring). This generates 16 space/timepoints. To plot the tileplot, we used the 'geom_tile' function from the ggplot2 R package. First, each cell cluster's percentage within a larger object (for example the 'Mono_Mac_1' cluster within the larger 'Mono_Mac' object) is calculated for each space/timepoint. These are then plotted on the 4x4 grid and color-coded based on their relative change compared to the unwounded state. Red indicates increase in subpopulation, white indicates the same percentage, blue indicates decrease in subpopulation compared to UW. Code is deposited and provided.

**Space/Time Correlation Analysis (STCA)**

To identify cell subsets in the scRNAseq data set that shared similar space/time profiles, we calculated the Pearson correlation coefficient, *R*, using the percentage for every space/timepoint between pairs of cell subsets. A Pearson correlation coefficient of *R* = 1 between a pair of cell subsets would indicate that they have the same space/time profile across the wound healing process. A Pearson correlation coefficient of *R* = -1 would indicate an negative correlation between the cell subsets, i.e. that the presence of the two cell subsets are inversely correlated across the wound. P-values were calculated using either Pearson or Spearman method (noted in legends). The Benjamini-Hochberg correction was applied for false discovery rate correction.

**Pseudotime Analysis in scRNAseq data set**

*Monocle analysis*

Raw counts from the individual cell-specific object were used to create a Monocle3[10–12] cell_data_set object, and the PCA and UMAP embeddings were imported directly from the Seurat object. Each cell-specific trajectory was inferred by reverse embedding the UMAP coordinates using the DDRTree method. The root cell states for the trajectory in the $MHCII^{low}$ and $MHCII^{hi}$ Mono_Mac objects were chosen based on which cell cluster was present on day 1 post-wounding. Relative pseudotime was obtained through a linear transformation relative to the cells with the lowest and highest pseudotimes ((1 - *min_pseudotime*) / *max_pseudotime*).

*Gene expression along pseudotime*

*Space/Time Tileplot of pseudotime*

Similar to the Space/Time Tileplot of cell frequencies described above, a tileplot for the $MHCII^{low}$ and $MHCII^{hi}$ Mono_Macs was created. The numbers in each tile represent the average pseudotime value of all cells found at that particular space/timepoint. 'D00' represents cells found in the unwounded skin and, therefore, is not split into different wound regions.

**Ce3D Tissue Clearing and Whole Mount Imaging**

*Tissue staining and clearing*

Wounds and surrounding tissue were excised from back skin using an 8 mm biopsy punch. Wound samples were cleared using the Ce3D clearing protocol[13] with minor modifications. First, wound samples were fixed in 4% PFA (16% PFA, Electron Microscopy Sciences, diluted in PBS) at 4ºC overnight on a horizontal shaker at 100 rpm by sandwiching them between two layers of cell strainer mesh to prevent sample curling. Subsequent steps were all performed protected from light. One wound sample was then transferred into a 2 ml tube and washed three times 30-60 min in 1 ml of wash buffer (PBS, 0.3% Triton X-100, 0.5% 1-thioglycerol) on a horizontal shaker at RT at 100 rpm. The inside of the 2 ml tube lid was plugged with a polydimethylsiloxane (PDMS) cut out to prevent the sample from being stuck during all subsequent washing, blocking, staining, and clearing steps. After washing, the sample was immersed in filtered blocking solution (PBS, 0.3% Triton X-100), 1% BSA, 1% normal mouse serum, and 1% normal serum of host species of used staining antibodies) overnight at 37ºC on a horizontal shaker at 100 rpm. The next day, the sample was stained in 1 ml blocking buffer supplemented with DAPI (1 ug/ml) for nuclear staining, anti-CD49f/ITGA6 AF488 (clone GoH3; 1:100) and anti-CD11b AF647 (clone M1/70; 1:100) antibodies. The sample was incubated for three days at 37ºC on a horizontal shaker at 100 rpm. After the staining incubation, the sample was transferred into a new 2 ml tube and washed once using 1 ml wash buffer for 8-12 h at 37ºC on a horizontal shaker at 100 rpm, and then three more times with 1 ml wash buffer for 8-12 h at RT on a horizontal shaker at 100 rpm. After washing, the sample was transferred into a new 2 ml tube for clearing by subsequently incubating the sample in 1 ml of 33%, 50%, 80%, or 100% Ce3D clearing solution (2.75 ml 40% N-methylacetamide, 4 g Histodenz, 5 µl Triton X-100, 25 µl 1-thioglycerol) diluted in wash buffer at RT on a horizontal shaker at 100 rpm. The 33%, 50%, and 80% Ce3D clearing steps were done for 1 h each and the 100% clearing step was done overnight. After clearing, the sample was mounted epidermal side facing down on a PDMS chamber that fit the sample in the middle. Ce3D clearing solution was used as the mounting medium.

*'Thick' section clearing:*

For 'thick' section (250-300 µm) clearing and imaging for both wounds and tumor samples, we started with whole tissue (wound or tumor) and fixed with 4% PFA at 4C overnight. This was followed by a progressive 15% to 30% w/v sucrose gradient then embedding and freezing in OCT. Thick sections were then made using a cryostat and placed into PBS to wash OCT residue away.

These sections were then carried through a the generalized Ce3D workflow[12]. Blocking and wash buffers same as above.

*'Thin' cryosection imaging:*

For thin (10 µm) sections, fresh tumor samples were embedded and frozen in OCT, then sectioned on the cryostat and transferred to slides. Samples were then fixed 2 hr 4C in 4% PFA before permeabilization for 15 min in 0.2% Triton X-100. Samples were then blocked using (PBS, 0.1% Triton X-100), 1% BSA, and 1% normal serum of host species of used staining antibodies) and stained in the same buffer. Following washes and incubation with 1 ug/mL DAPI for 5 min., slides were washed and coverslipped using VectaShield.

*Image acquisition and analysis*

All samples were imaged using a Leica SP8 laser scanning confocal microscope with a white light laser and 405 nm diode. For whole cleared wounds, a 16X 0.6NA (HC Fluotar L 16x/0.6 IMM CORR DLS, Leica) objective was used during acquisition. 'Thick' (250-300 µm) and 'thin' (10 µm) cryosections were imaged using a 20X 0.75NA (HC PL APO 20x/0.75 IMM CORR CS2, Leica) objective. After acquisition, individual tiled images were stitched together using the LAS X software (Leica) and then analyzed using the Imaris software suite (Bitplane).

*Imaris processing*

For large volume images in Figure 3, spots were created using the Imaris Spots function and inputting the DAPI nuclear stain signal. Parameters for Spots generation are listed in Supplementary Table S3. This first step is done to locate and identify individual cells within the imaged sample. Spherical spots here are created based on signal above a user-defined threshold value in the DAPI channel and based on the size of the DAPI object. Next, Surfaces for channels CD11b, Arg1-tdTomato, and CD206 were created based on the parameters listed in Supplementary Table S3 using the Imaris Surfaces function. Surface marker-positive cells were identified by their vicinity to a previously identified DAPI spot. Depending on surface marker (see Supplementary Table S3 for details), DAPI spots 5-20 $\mu$m close to the Surface were identified as $CD11b^+$, $Arg1^+$, and/or $CD206^+$ using the Find Spots Close to Surface function in Imaris. To identify $CD11b^+$ $Arg1^+$ or $CD11b^+$ $CD206^+$ double positive cells, the Colocalize Spots function selecting CD11b -and Arg1-positive spots or CD11b- and CD206-positive spots, respectively, with a threshold of =<1 $\mu$m was used in Imaris.

*Generate Distance transformation from center of wound and count cells*

To quantify the number of cells per area of wound in the Imaris software, first, a user-defined 'center' spot was selected in the wound that represents the center of the wound. This spot was duplicated to create a new spot object. Then, the duplicated 'center' spot was selected and using the 'Distance Transformation' tool under 'Tools -> Create new Channel' a new channel was generated, wherein the channel intensity across the sample reflects the distance from the previously selected center spot. Areas close to the center spot have a low intensity and areas distant from the center have a high intensity in the 'Distance Transformation' channel. To calculate how many cells/spots are located at what distance from the center of the wound, the 8 mm wound cutout used for imaging was subdivided into 8 circular bins (0-0.5 mm distance from center of wound; 0.5-1.0; 1.0-1.5; 1.5-2.0; 2.0-2.5; 2.5-3.0; 3.0-3.5; 3.5-4.0). Cells/spots with a cell marker of interest was selected, for example 'CD11b+' spots, and a new filter was created called 'Intensity Mean' of the 'Distance Transformation' channel. Selection of the maximum and minimum filter settings reflect which of the 8 circular bins all 'CD11b+' spots are counted. The same was done for all 'CD11b+ Arg1-tdTomato+' spots and the percentage of 'CD11b+ Arg-tdTomato+ of CD11b+' spots was calculated for each of the 8 circular bins and plotted. The same analysis was done for 'CD11b+ CD206+' spots.

**Ex vivo experiments of primary skin fibroblast and bone marrow-derived macrophages (BMDMs)**

*Primary skin fibroblast isolation*

6-12 week old C57BL/6 mice were euthanized and their ventral skin covering the thorax was shaved and residual hair were removed by NAIR application. The skin was excised and remaining subcutaneous fat was removed before the tissue was finely minced with scissors. The tissue was then transferred into a 2 ml tube containing 1 ml of digestion medium (2 mg/ml collagenase XI, 0.5 mg/ml hyaluronidase, 0.1 mg/ml DNase in RPMI with 10% FCS, 2 mM L-glutamine, 100 U/ml penicillin, 100 µg/ml streptomycin, 50 µM beta-mercaptoethanol). Tissue from one skin excision was split into 3 tubes. The tubes were placed horizontally in a bacterial shaker for 90 min at 37ºC and 225 rpm. After digestion, the samples were filtered on a 100 µm cell strainer and the digestion was quenched by adding 20 ml of ice cold RPMI. The single cell suspension was then seeded at a concentration of $2x10^4$ cells / $cm^2$ in fibroblast media (DMEM supplemented with 10% FCS (Benchmark), 2 mM L-glutamine, 50 µM beta-mercaptoethanol, 100 U/ml penicillin, 100 µg/ml streptomycin, and 250 ng/ml amphotericin B) at 37ºC in 5% $CO_2$. After 3 days or when culture reached 80-90% confluency, cells were collected, aliquoted, and stored in liquid nitrogen (passage 1). For subsequent co-culture experiments, cells were thawed and cultured in fibroblast media for one more passage before use in experiments.

*Generation of bone marrow-derived macrophages (BMDMs)*

6-12 week old C57BL/6 mice were euthanized and their femurs and tibiae were excised. Using PBS, the bone marrow was flushed from the long bones after cutting off the ends. After pelleting the bone marrow, the red blood cells were lysed using RBC lysis buffer (155 mM $NH_4Cl$, 12 mM $NaHCO_3$, 0.1 mM EDTA) for 5 min at RT. Cells were washed with DMEM and filtered through a 40 µm cell strainer before being seeded at $1x10^6$ cells / ml on a low-adherent cell culture dish in BMDM media (DMEM supplemented with 10% FCS (Benchmark), 2 mM L-glutamine, 50 µM beta-mercaptoethanol, 100 U/ml penicillin, 100 µg/ml streptomycin, and 20 ng/ml M-CSF). After 3 days in culture, cells were split 1:2 into new low-adherent cell culture dishes with fresh BMDM media. On day 6-7 of culture, BMDMs were harvested and used for experiments.

For Oncostatin M stimulation, PSF's were grown to 80% confluency, before switching to DMEM+1% FCS with 50 ng/mL of recombinant OSM (Biolegend) and incubated 24hrs at 37C before harvest for RNA extraction.

Conditioned BMDM media from $Grn^{-/-}$ was generated as follows: BMDM's were generated as described above from bone marrow from $Grn^{-/-}$ mice. On D7 following BMDM isolation and differentiation, the media was replaced with DMEM+1%FCS and harvested 24 hours following incubation. PSF's were grown till 80% confluent then media was replaced with $Grn^{-/-}$ BMDM CM with or without 1 ug/mL recombinant Progranulin (R&D). PSF's were incubated at 37C for 24hrs before harvest for RNA extraction.

For Periostin treatment, Collagen IV coated 6 well plates (Corning 354428) were coated with 1 ug/mL recombinant mouse Periostin (R&D 2955-F2-050) overnight at 4C. Plates were then washed twice with PBS before cell plating. BMDM's D8 post-isolation and differentiation were then plated at a density of 750k cells per well and briefly spun down at 200g for 15 seconds to force them into contact with the plate. BMDM's were then incubated 24 hrs at 37C before RNA extraction.

*Reverse transcription real-time quantitative PCR*

Following cell harvest and washing 1X cold PBS, cells were lysed and RNA extraction performed using the RNeasy kit from Qiagen. Roughly equal mass of RNA was then used for reverse transcription using the iScript Master Mix. The reaction mix was directly used for RT-qPCR with the SsoFast EvaGreen kit and read out on a Bio-Rad thermocycler with the following cycling protocol: [95C for 30s, 95C for 5s, 60C for 2s, repeat from step 2 44x, 65C to 95C melt curve with 0.5C increments at 5s each]. Primer sequences can be found in the key resources table. We confirmed that melt curves were consistent across samples and the non-template controls has no appreciable amplification. Cq was calculated at roughly ¼ the maximum signal and then corrected against 18s Cq before conversion to relative expression. Shown bar charts are representative of the mean+SE of 3 technical replicate wells.
