## Supplemental Figures for "Space-Time Mapping Identifies Concerted Multicellular Patterns and Gene Programs in Healing Wounds and their Conservation in Cancers"

**Supplementary Information**

**Supplementary Figure S1.**

(**A)** Diameter of wounds plotted over time of two separate experiments. Each dot represents the mean diameter of 4-12 wounds from 1-3 mice per timepoint. Error bars represent ±SD.

**(B)** Heatmap of marker expression across all identified cell clusters. Mean intensity of each marker is plotted as scaled value across column.

**(C)** Histogram plots of selected markers detected in Neutrophils_1 (black), Neutrophils_2 (red), and pre-Neutrophils (blue).

**(D)** Top: Gating strategy to identify $CD4^+$ $Foxp3^+$ T cells (red), $\gamma\delta^+$ T cells (black), $\gamma\delta^{mid}$ dermal T cells (blue), and $\gamma\delta^-$ $CD4^+$ T cells (green). Bottom: UMAP projection of all immune cells (gray) and gated cells from top highlighted by color.

**(E)** Line plot of $CD4^+$ T cell subsets plotted as percentage of all $CD45^+$ immune cells per timepoint of wound sampling. Dots represent mean values of n=3 mice per timepoint. Error bars represent ±SD.

**(F)** UMAP projection of immune cells detected during skin wound repair 'Experiment #2' based on a 40-marker panel.

**(G)** Line plots of all immune cell clusters identified in (C) plotted as percentage of all $CD45^+$ immune cells per timepoint of wound sampling (left y-axis). Dotted line represents wound diameter (right y-axis). Dots represent mean values of n=3 mice per timepoint. Error bars represent ±SD.

**(H)** Phenotypic earth mover's distance (PhEMD) diffusion map embedding of all samples collected during wound closure of 'Experiment #2'. Each dot represents the wound-resident immune cell composition of one mouse. Dots are color-coded by day and n=3 mice were collected per day. UW, unwounded. DC, diffusion coefficient.

**Supplementary Figure S2.**

**(A)** UMAP plot of $CD45^+$ immune cells found during skin repair.

**(B)** Bubble plot of differentially expressed genes across all identified clusters in (A).

**(C)** Bubble plot of differentially expressed genes across all Mono_Mac subpopulations.

**(D)** UMAP plot of all monocyte/macrophage cells. Violet color intensity indicates mRNA expression level of *H2-Ab1* (gene encoding structural component of MHCII complex) and *Nr4a3* within each cell.

**(E)** UMAP plot of $MHCII^{hi}$ Mono_Mac subset.

**(F)** Gene expression of select genes along pseudotime. D00 represents unwounded skin.

**(G)** Space-time tile plot representing the distribution of all cells in the $MHCII^{lo}$ subset according to their presence in pseudotime. Each square represents the average pseudotime value of all cells within that space-time point. Squares are color-coded according to the pseudotime value. Dark violet, low. Yellow, high. D00 represents the unwounded skin and has no wound areas.

**(H)** Pseudotime trajectory using Monocle 3 on $MHCII^{hi}$ subset starting at Mono_MHCII, progressing through Mono_Mac_MHCII cluster and branching off to the Mono_Mac_MHCII_Mgl2 cluster.

**(I)** Violin plot of all cells within $MHCII^{hi}$ Mono_Mac subset split by day post-wounding and plotted according to their presence in pseudotime. D00 represents unwounded skin.

**(J)** Violin plot split by the $MHCII^{hi}$ monocyte/macrophage subpopulations and plotted according to their distribution in pseudotime.

**(K)** Same as (G) for $MHCII^{hi}$ Mono_Mac subset.

Left: Line plots of all $MHCII^{lo}$ monocyte/macrophage subpopulations identified in the scRNAseq dataset during wound healing. Percentage of each subpopulation within all Mono_Mac cells is plotted by day post-wounding.

**(L)** Line plots of all Mono_Mac subpopulations identified in the scRNAseq dataset during wound healing. Percentage of each subpopulation within all Mono_Mac cells is plotted by day post-wounding.

**(M)** Space-time tile plots representing the Mono_2 and Mono_MHCII subpopulations.

**(N)** Left: Line plots of all CD45$^+$ subpopulations identified in the scRNAseq dataset during wound healing. Percentage of each subpopulation within all CD45$^+$ cells is plotted by day post-wounding. Right: Space-time tile plot representing the CD45$^+$ subpopulations.

**Supplementary Figure S3.**
**(A)** Top-down view of D3 post-wounding imaged sample. Fluorescent stain of CD49f/ITGA6 is shown in white. *Edge of wound with re-established epithelial basement membrane. See Movie S1 for 3D view.
**(B)** Top-down view of D7 post-wounding imaged sample. Fluorescent stain of CD49f/ITGA6 is shown in white. *Vasculature in non-wounded skin, **hair follicles, ***fascia, #severed nerve bundle. See Movie S2 for 3D view.
**(C)** Top: UMAP plot of all Mono_Mac clusters. Blue color intensity indicates mRNA expression level of *Arg1* within each cell. Bottom: UMAP plot of Mono_Mac subsets.
**(D)** Top-down view of stained wound sample from *Arg1*-tdTomato reporter mouse (top) day 3 and (bottom) day 7 post-wounding. Boxes highlight zoom-ins shown in (E). One of two representative images is shown for each day.
**(E)** Zoom-in of boxes from (D) displaying single z-stack. Images depict wound center region (i) and (iii) or wound distal region (ii) and (iv). CD11b$^+$ *Arg1*-tdTomato$^+$ double-positive cells are highlighted by an asterisk, which are predominantly found in the wound center on day 3. Bar, 50 μm.
**(F)** UMAP plot of all CD45$^+$ immune cells. Blue color intensity indicates mRNA expression level of *Mrc1* within each cell.
**(G)** Top-down view of stained wound sample from WT mouse (top) day 3 and (bottom) day 7 post-wounding. Boxes highlight zoom-ins shown in (H). One of two representative images is shown for each day.
**(H)** Zoom-in of boxes from (G) displaying single z-stack. Images depict wound center regions (i) and (iii) or wound distal regions (ii) and (iv). CD11b$^+$ CD206$^+$ double-positive cells are highlighted by an asterisk, which are only found in unwounded skin on day 3 and day 7, and on the wound edge on day 7. Bar, 50 μm.

**Supplementary Figure S4.**
**(A)** Bubble plot of differentially expressed genes across all identified CD45$^-$ non-immune cell clusters. Cluster labelled 'immune' expressed high levels of the MHCII invariant chain *Cd74* and was excluded from further analysis.
**(B-O)** Matching line plots and space-time tile plots of all CD45$^-$ non-fibroblast non-immune cells identified in the scRNAseq dataset during skin repair. Percentage within all CD45$^-$ non-immune cells is plotted by day and space post-wounding.
**(P)** Bubble plot of differentially expressed genes across all fibroblast subpopulations identified during wound skin repair.
**(Q)** Heatmap depicting Jaccard similarity of cluster-specific genes between fibroblast clusters identified in Buechler et al., 2021, and fibroblast subpopulations in present study. About 25% of all differentially expressed genes in Lrrc15 cluster from Buechler et al., 2021, and Fibro_4 cluster in present study were shared.
**(R)** Top: gene expression density plots of the inflammatory marker genes *Il6*, *Cxcl2*, and *Ccl2* described in 'inflammatory cancer-associated fibroblasts (iCAFs)'. Bottom: gene expression density plots of genes described in 'antigen-presenting cancer-associated fibroblasts (apCAFs)'[61,62]: *Cd74*, *H2-Ab1*, and *Saa3*.
**(S)** Pearson correlation matrix output of STCA comparing all identified CD45$^+$ immune and CD45$^-$ non-immune cell subpopulations. Dots shown represent statistically significant pairs ($p$-value<0.05) and color indicates Pearson's correlation coefficient.
**(T)** Table listing cell types and subpopulations according to STCA patterns. vSM, vascular smooth muscle.

**Supplementary Figure S5.**

**(A)** Heatmap showing gene weights for the top 8 contributing genes for each factor in the Mono_Mac subset. Values shown are normalized across the rows such that each gene has a maximal contribution of 1 towards a given factor.
**(B)** Same analysis as in (A) for the fibroblast subset.
**(C)** Plot showing strategy for choosing optimal number of factors for NMF decomposition. The factor number prior to a sharp downturn in cophenetic score (stability measure) was chosen.
**(D)** CellChat stacked bar plot showing relative information flow computed within the Mono_Mac. and Fibroblast merged datasets split by timepoint (D00, D1, D3, D7 and D14 following wounding)
**(E)** CellChat hierarchy plots to depict 'likelihood' of cell-cell interaction based on expression levels of the Periostin signaling elements (*Postn* and *Itgav/Itgb3*). The fibroblast and Mono_Macrophage objects were merged, then split based on timepoint. These individual 'timepoint' objects were used as the starting point for the CellChat workflow. The diameter of circles represent relative abundance of the cell cluster at the timepoint while the thickness of the line represents relative likelihood of interaction.
**(F)** Correlation matrix between the space-time profiles of average levels of MM (=Mono_Mac) factors and the frequencies of CD45 positive cell clusters, excluding Mono_Mac. Values shown representative of Pearson's correlation coefficient and significance testing performed using Pearson's test with a Benjamini-Hochberg multiple comparisons correction. * $p<0.05$ **$p<0.005$
**(G)** Correlation matrix between the space time profiles of average levels of M_M factors and the frequencies of CD45 negative cell clusters. Values shown representative of Pearson's correlation coefficient and significance testing performed using Pearson's test with a BH multiple comparisons correction. * $p<0.05$ **$p<0.005$
**(H)** Correlation matrix showing the space-time correlation between 114 factors generated from each of the coarse-grained cell type definitions identified in Figures S2A and 4A. Dots shown represent statistically significant pairs (p-value<0.05) and color indicates Pearson's correlation coefficient. Abbreviations: endo, endothelial; dc, dendritic cell; MM, Mono_Mac; fibro, fibroblast; kerat, keratinocyte; tnk, T and NK cell; t, T cell; mast, mast cell; dsp, dermal sheath papilla; vsm, vascular smooth muscle; Neut, neutrophil; melano, melanocyte; b, B cell.
**(I)** Tile plots showing average factor expression over space-time for the 114 identified gene programs in Figure S5H. Factors are ordered via hierarchical clustering in the correlation matrix with the bands identifying factors within the same clusters delineated on the correlation matrix. Several clusters matching the previously defined ST patterns (Early/Int-In/Late-In/Late-Ex/Edge) are highlighted with matching colors. Red tiles denote increase relative to the unwounded state while blue tiles denote decrease relative to unwounded.

**Supplementary Figure S6.**
**(A)** UMAP dimensional reduction on integrated MC38 and B16F10 Mono_Mac datasets (n=3859 cells)
**(B)** Heatmap showing gene weights for the top 8 contributing genes for each factor in the Mono_Mac combined B16F10 and MC38 dataset. Values shown are normalized across the rows such that each gene has a maximal contribution of 1 towards a given factor.
**(C)** Plot showing strategy for choosing optimal number of factors for NMF decomposition. The factor number prior to a sharp downturn in cophenetic score (stability measure) was chosen.
**(D-I)** Scatter plots for selected tumor/WH factor pairs for **(D)** Tumor factor-4 vs WH factor-11, **(E)** Tumor factor-5 vs WH factor-12, **(F)** Tumor factor-7 vs WH factor-15, **(G)** Tumor factor-10 and WH factor-8, **(H)** Tumor factor-11 and WH factor-14, and **(I)** Tumor factor-12 and WH factor-1 with the gene weight contributions plotted as calculated from the basis matrix in the NMF output (see Figure S5A for WH factors and S6B for tumor factors). Slope represents x=y line and dotted lines represent the weight for the $20^{th}$ highest gene contribution in either factor. The $Jaccard_{20}$ index is shown and thus reflects the frequency of points in quadrant I over quadrants I,II and IV. For pairings in **D-I**, top shared genes in the upper right quadrant were put through Enrichr to find overrepresented cellular processes with the top result by p-value listed. Full Enrichr output can be found in the extended data.

**Supplementary Figure S7.**
**(A)** POSTN$^+$ surfaces generated via Imaris overlaid on top of IF images from B16F10 and MC38 tumors. Magenta dots denote DAPI$^+$ spots in close association (within 3 μm) of a CD11b$^+$ surface (defined as CD11b$^+$ Cells) and yellow signal denotes the distance transform from POSTN surfaces. Images representative of 3 separate tumor samples per type.
**(B)** 3D projections of cleared tissue imaging from 250 μm thick tumor sections from B16F10 and MC38 tumors stained as indicated. Images representative of 4 tumor samples per type.
**(C)** Histograms indicating the distances of MHCII$^+$ spots and MHCII$^-$ spots to the nearest Selectin-P$^+$ or Selectin-P$^-$ surface in the MC38 and B16F10 models. Dashed line indicates the median. Histograms representative of 4 independent replicates (4 separate tumors).

**Supplementary Movie S1.**
Clearing-enhanced (Ce3D) skin wound D3 post-wounding of wild-type C57BL/6 mouse. Original wound size, 4 mm. Movie starts with internal-facing view of wound and DAPI (blue), ITGA6/CD49b (white), CD11b (green), and CD206 (red) channels on. Individual channels are turned on and off as wound image rotates along horizontal axis. ITGA6/CD49b staining (white) highlights re-established epithelial basement membrane, vasculature structures in non-wounded skin, hair follicles, and fascia.

**Supplementary Movie S2.**
Clearing-enhanced (Ce3D) skin wound D7 post-wounding of wild-type C57BL/6 mouse. Original wound size, 4 mm. Movie starts with internal-facing view of wound and DAPI (blue), ITGA6/CD49b (white), CD11b (green), and CD206 (red) channels on. Individual channels are turned on and off as wound image rotates along horizontal axis. ITGA6/CD49b staining (white) highlights re-established epithelial basement membrane, vasculature structures in non-wounded skin, hair follicles, fascia, and severed nerve bundles.

**Supplementary Movie S3.**
Clearing-enhanced (Ce3D) skin wound D3 post-wounding of *Arg1*-tdTomato reporter mouse. Original wound size, 4 mm. Movie starts with internal-facing view of wound and DAPI (blue), ITGA6/CD49b (white), CD11b (green), and *Arg1*-tdTomato (red) channels on. Individual channels are turned on and off as wound image rotates along horizontal axis. ITGA6/CD49b staining (white) highlights re-established epithelial basement membrane, vasculature structures in non-wounded skin, hair follicles, and fascia.

**Supplementary Movie S4.**
Clearing-enhanced (Ce3D) skin wound D7 post-wounding of *Arg1*-tdTomato reporter mouse. Original wound size, 4 mm. Movie starts with external-facing view of wound and DAPI (blue), ITGA6/CD49b (white), CD11b (green), and *Arg1*-tdTomato (red) channels on. Individual channels are turned on and off as wound image rotates along horizontal axis.

**Supplementary Movie S5.**
Movie showing spatiotemporal changes in factor 'expression' level across all 114 identified factors. Concentric rings denote the 2,4,6,8mm and UW respectively moving from center outwards. Bar height represents level of mean factor expression at each space-time coordinate normalized to the maximum value across all 17 space-time coordinates. Bars color-coded based on cell type the factor belongs to. Brackets on the edge denote the groupings of factors based on hierarchical clustering in Supplementary Figure 5H. Several frames between each timepoint were generated using interpolation to allow for smoothing of the transitions.

**Supplementary Table 1.**

Top 20 contributing genes by gene weight from basis matrix shown for each of the 114 identified factors along with their gene weight towards that factor.

**Supplementary Table 2.**

Genes in the upper right quadrant (I) of the tumor by wound healing M_M factor scatterplots (Figure 6C-E and Supplementary Figure SD-I) were used for GO analysis. Table shows a truncated output from each query for each pair with $J_{20} > 0.25$

**Supplementary Table 3.**

Parameters used for surface and spot generation using Imaris software.

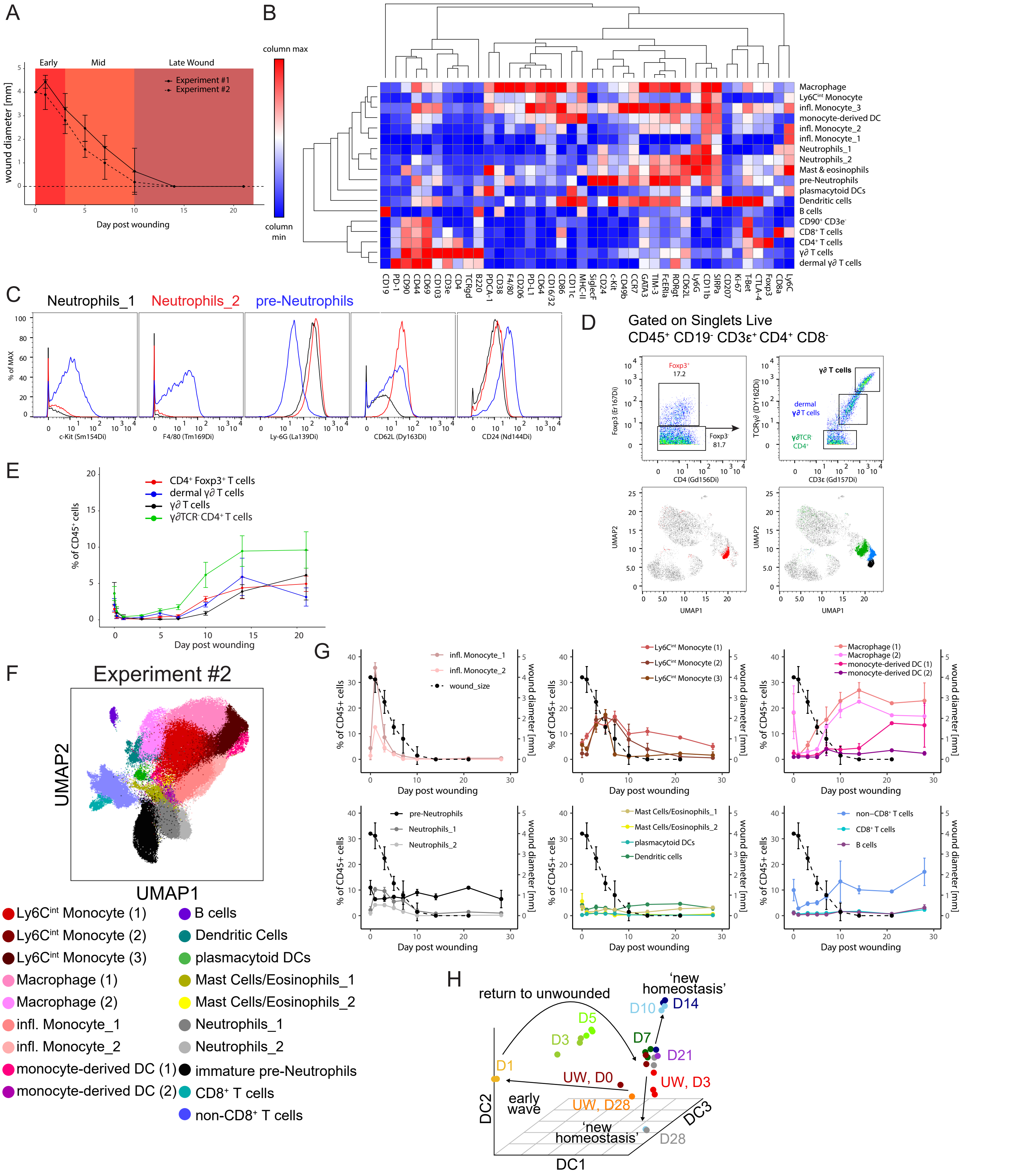
A
B
C
D
E
F
G
H
Early
Mid
Late Wound
Experiment #1
Experiment #2
wound diameter [mm]
Day post wounding
column max
column min
Macrophage
Ly6C^int Monocyte
infl. Monocyte_3
monocyte-derived DC
infl. Monocyte_2
infl. Monocyte_1
Neutrophils_1
Neutrophils_2
Mast & eosinophils
pre-Neutrophils
plasmacytoid DCs
Dendritic cells
B cells
CD90+ CD3e-
CD8+ T cells
CD4+ T cells
γ∂ T cells
dermal γ∂ T cells
CD19 PD-1 CD90 CD44 CD69 CD103 CD3e CD4 TCRgd B220 PDCA-1 CD38 F4/80 CD206 PD-L1 CD64 CD16/32 CD86 CD11c MHC-II SiglecF CD24 c-Kit CD49b CCR7 GATA3 TIM-3 FcERIa RORgt CD62L Ly6G CD11b SIRPa CD207 Ki-67 T-Bet CTLA-4 Foxp3 CD8a Ly6C
Neutrophils_1
Neutrophils_2
pre-Neutrophils
% of MAX
c-Kit (Sm154Di)
F4/80 (Tm169Di)
Ly-6G (La139Di)
CD62L (Dy163Di)
CD24 (Nd144Di)
Gated on Singlets Live
CD45+ CD19- CD3ε+ CD4+ CD8-
Foxp3+
17.2
Foxp3
81.7
Foxp3 (Er167Di)
CD4 (Gd156Di)
γ∂ T cells
dermal γ∂ T cells
γ∂TCR- CD4+
TCRγ∂ (DY162Di)
CD3ε (Gd157Di)
UMAP2
UMAP1
CD4+ Foxp3+ T cells
dermal γ∂ T cells
γ∂ T cells
γ∂TCR- CD4+ T cells
% of CD45+ cells
Day post wounding
Experiment #2
UMAP2
UMAP1
Ly6C^int Monocyte (1)
Ly6C^int Monocyte (2)
Ly6C^int Monocyte (3)
Macrophage (1)
Macrophage (2)
infl. Monocyte_1
infl. Monocyte_2
monocyte-derived DC (1)
monocyte-derived DC (2)
B cells
Dendritic Cells
plasmacytoid DCs
Mast Cells/Eosinophils_1
Mast Cells/Eosinophils_2
Neutrophils_1
Neutrophils_2
immature pre-Neutrophils
CD8+ T cells
non-CD8+ T cells
infl. Monocyte_1
infl. Monocyte_2
wound_size
Ly6C^int Monocyte (1)
Ly6C^int Monocyte (2)
Ly6C^int Monocyte (3)
Macrophage (1)
Macrophage (2)
monocyte-derived DC (1)
monocyte-derived DC (2)
pre-Neutrophils
Neutrophils_1
Neutrophils_2
Mast Cells/Eosinophils_1
Mast Cells/Eosinophils_2
plasmacytoid DCs
Dendritic cells
non−CD8+ T cells
CD8+ T cells
B cells
% of CD45+ cells
wound diameter [mm]
Day post wounding
return to unwounded
'new homeostasis'
D1
D3
D5
D7
D10
D14
D21
UW, D0
UW, D3
UW, D28
early wave
'new homeostasis'
D28
DC1
DC2
DC3

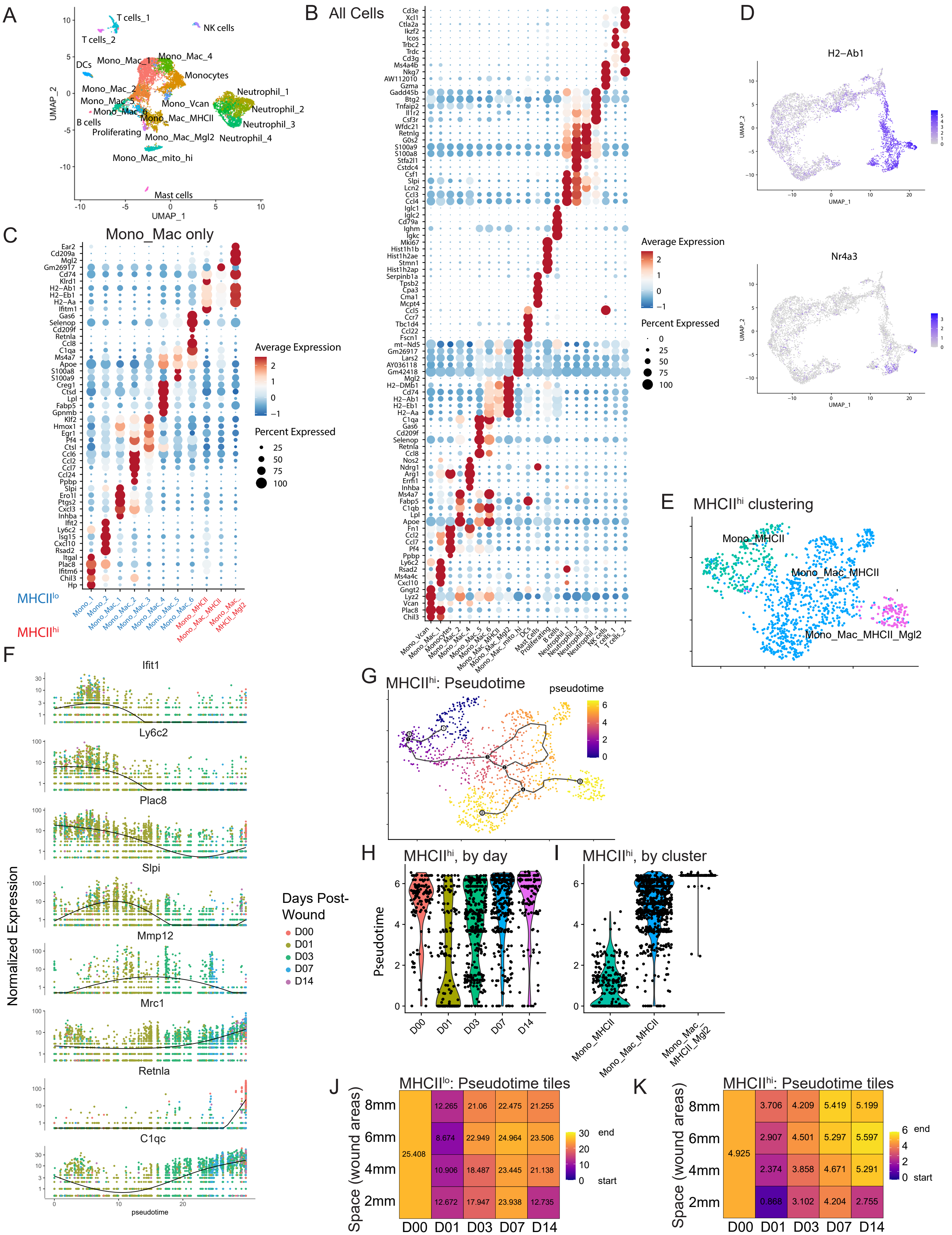


Supplementary Figure 2

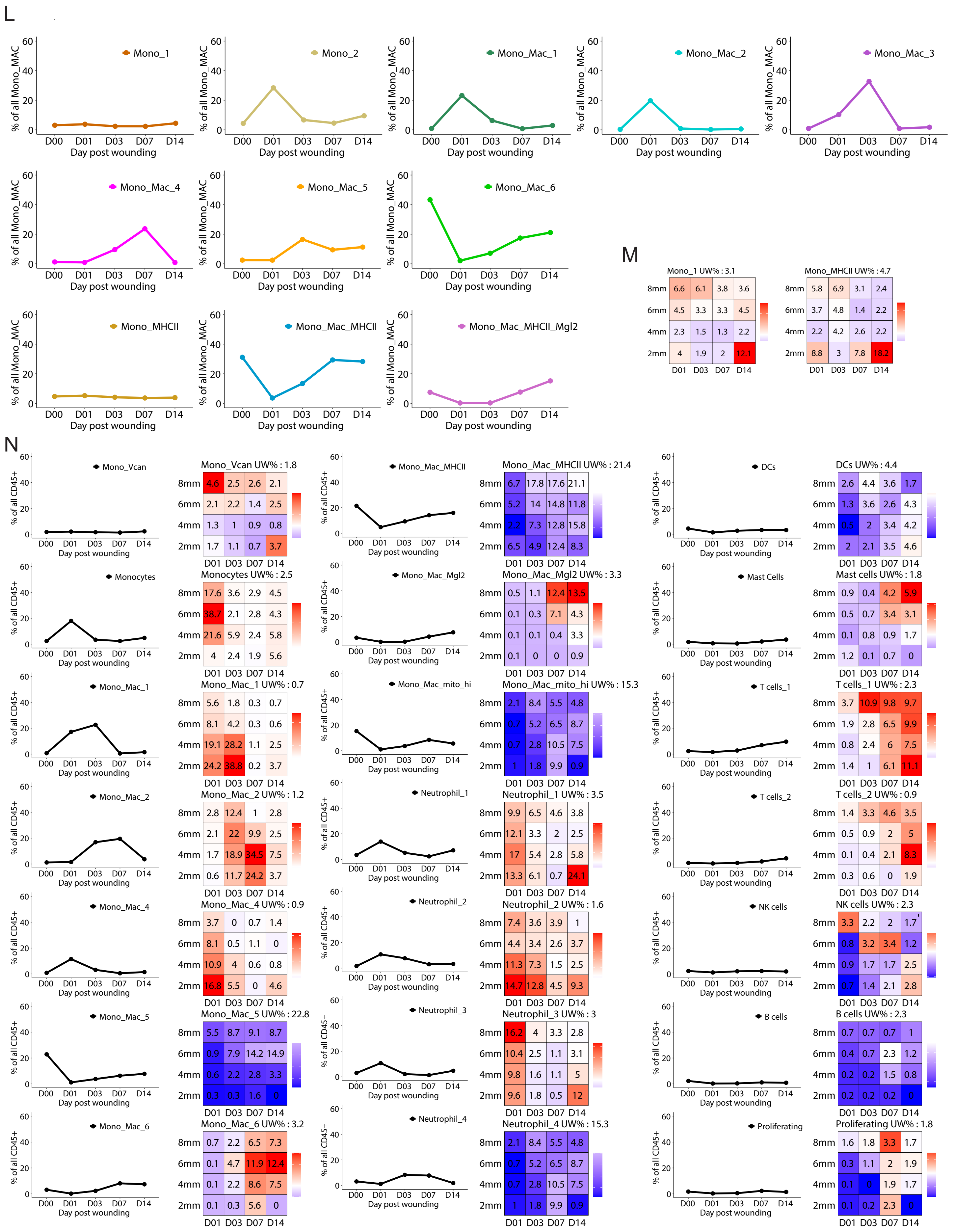


Supplementary Figure 2

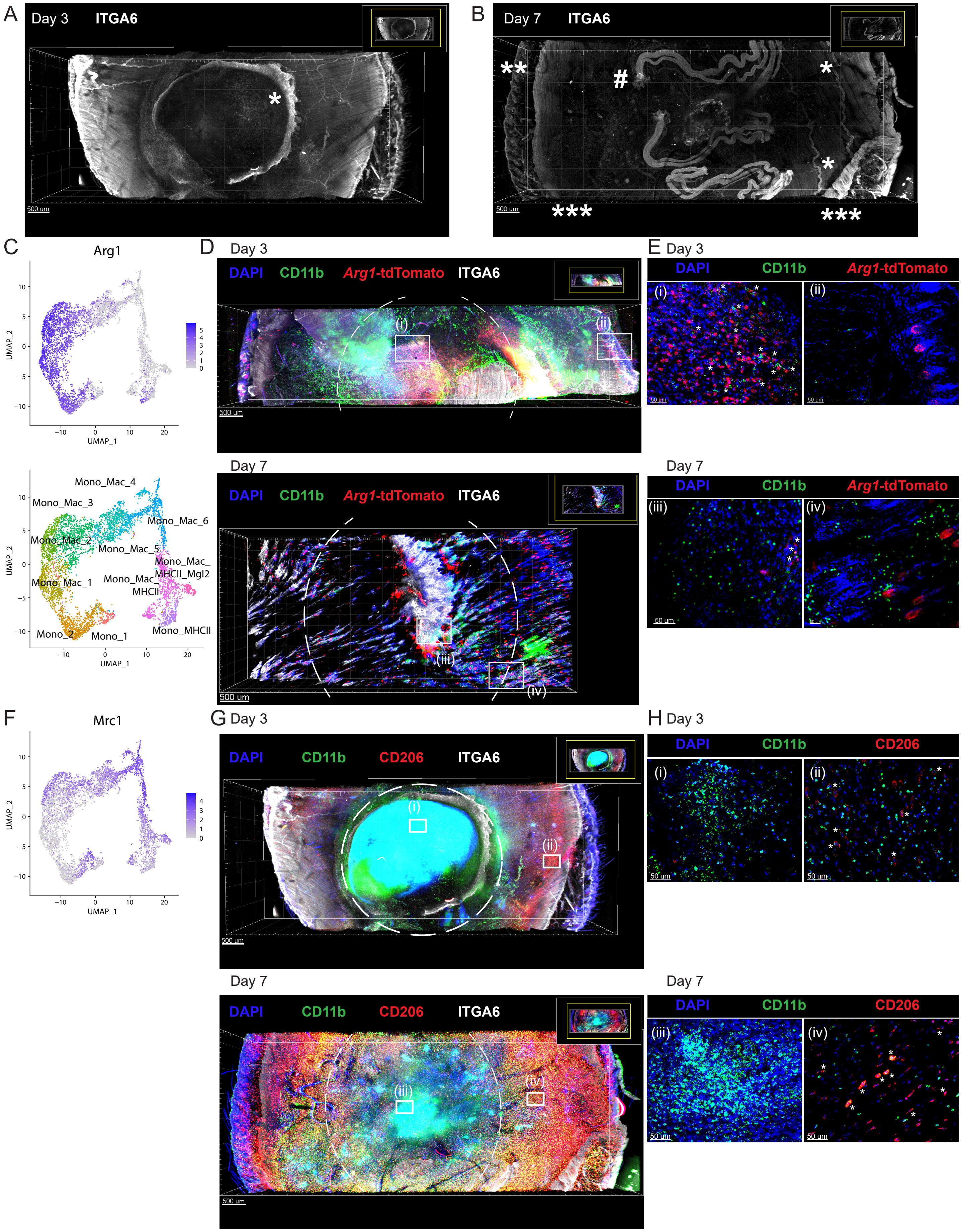


Supplementary Figure 3

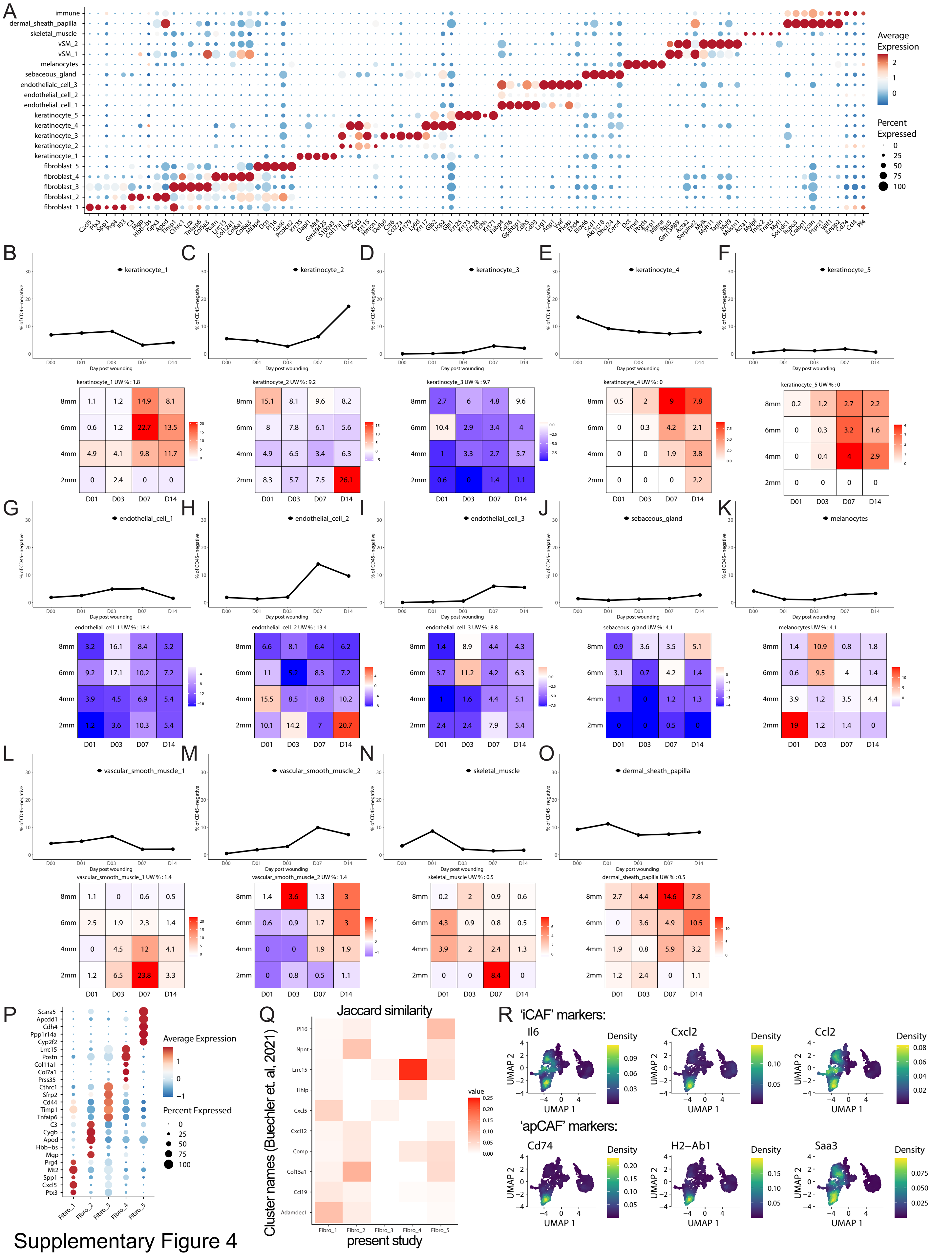


Supplementary Figure 4

S

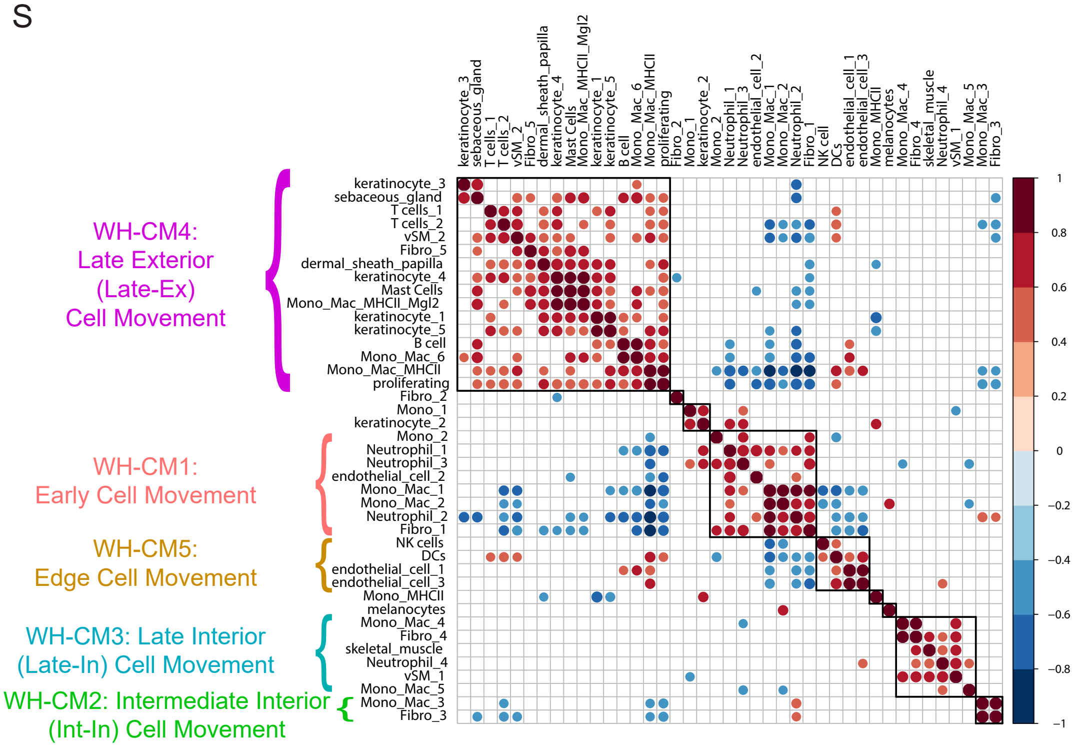


T

Pattern

Cell Type

| | WH-CM1: Early | WH-CM2: Int-In, Intermediate Interior | WH-CM3: Late-In, Late Interior | WH-CM4: Late-Ex, Late Exterior | WH-CM5: Edge | Other |
|---|---|---|---|---|---|---|
| Neutrophils | + Neutrophil_1,2,3 | - | + Neutrophil_4 | - | - | - |
| Mono_Mac | + Mono_2, Mono_Mac_1,2 | + Mono_Mac_3 | + Mono_Mac_4,5 | + Mono_Mac_6, Mono_Mac_MHCII, Mono_Mac_MHCII_Mgl2 | - | + Mono_1, Mono_MHCII |
| Fibroblasts | + Fibro_1 | + Fibro_3 | + Fibro_4 | + Fibro_5 | - | + Fibro_2 |
| Keratinocytes | - | - | - | + keratinocytes_1,3,4,5 | - | + keratinocytes_2 |
| Endothelial | + endothelial_cell_2 | - | - | - | + endothelial_cell_1,3 | - |
| Mast cells | - | - | - | + | - | - |
| Dendritic cells | - | - | - | - | + | - |
| lymphoid | - | - | - | + B cell, T cells_1,2 | + NK | - |
| sebaceous gland | - | - | - | + | - | - |
| dermal sheath papilla | - | - | - | + | - | - |
| muscle | - | - | + vSM_1, skeletal muscle | + vSM_2 | - | - |
| melanocytes | - | - | - | - | - | + melanocytes |

Supplementary Figure 4

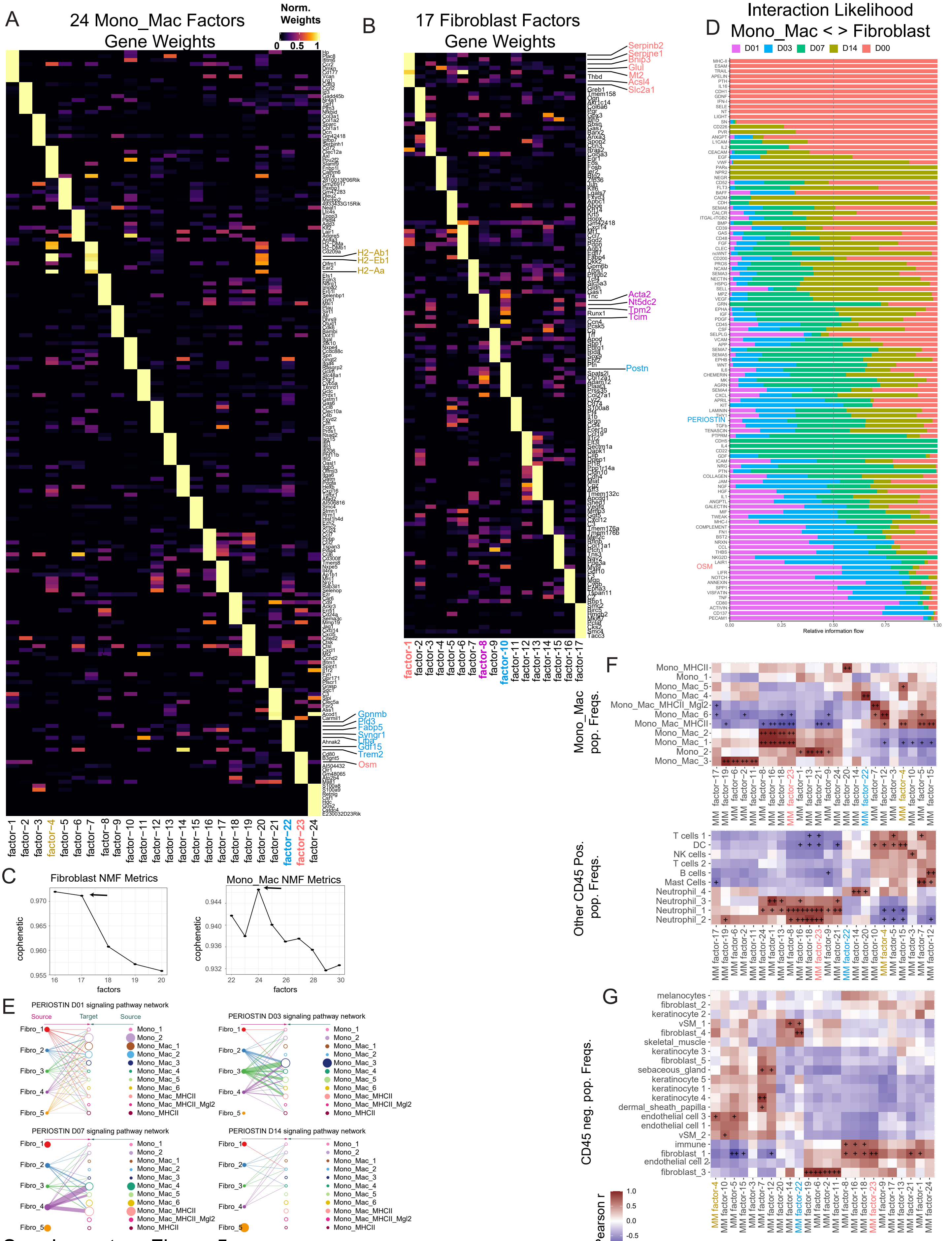


Supplementary Figure 5

H

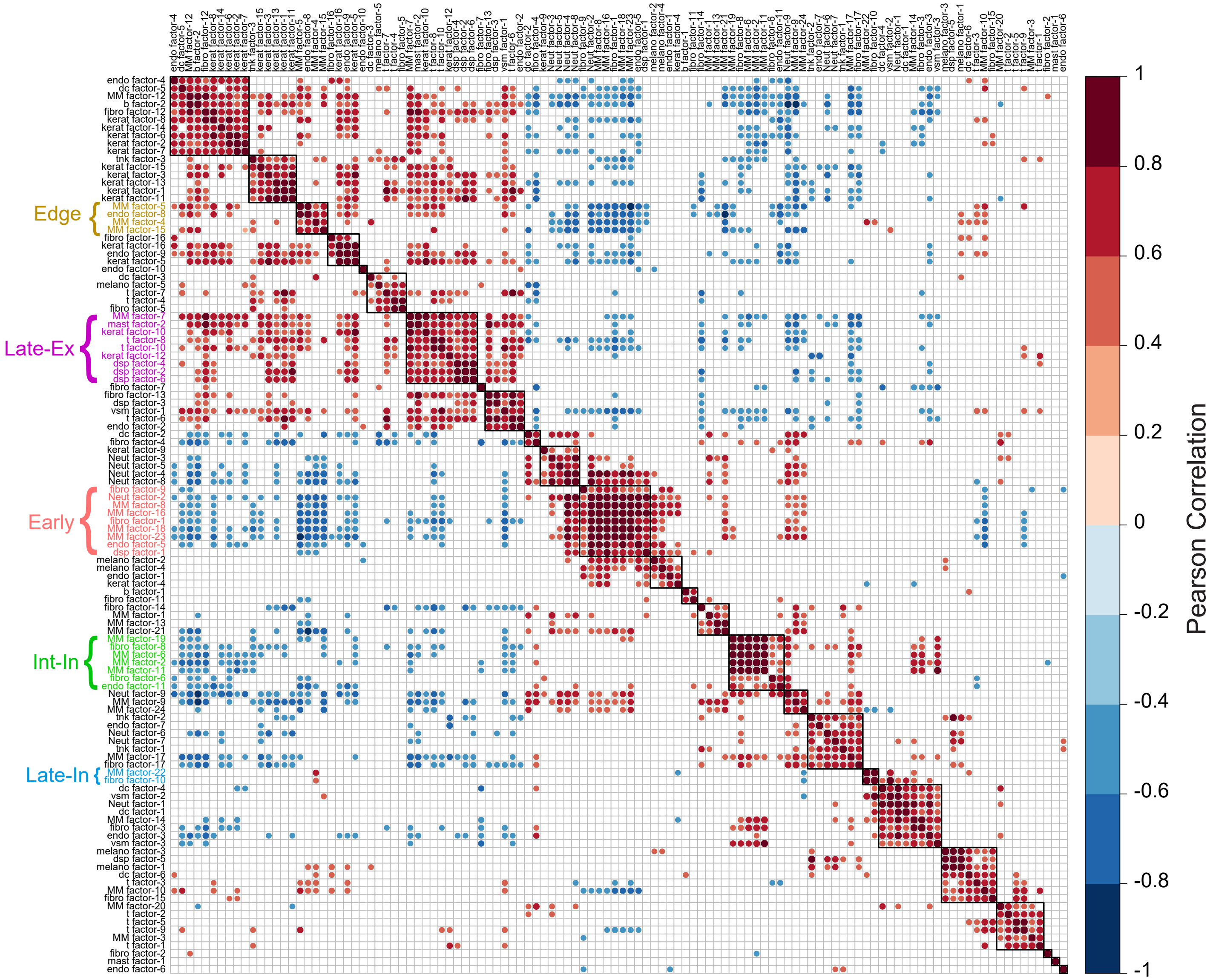


Supplementary Figure 5

I

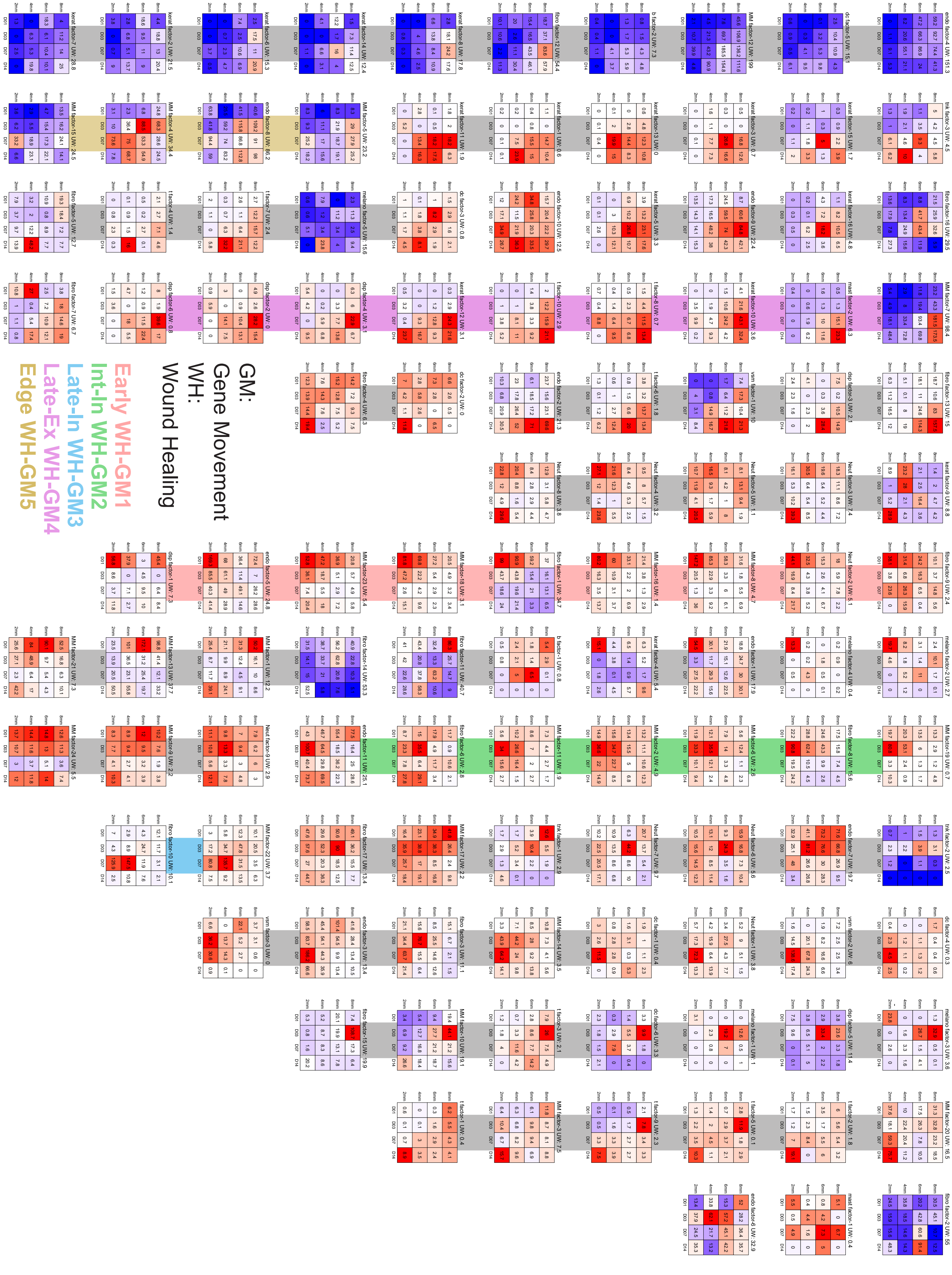


Supplementary Figure 5

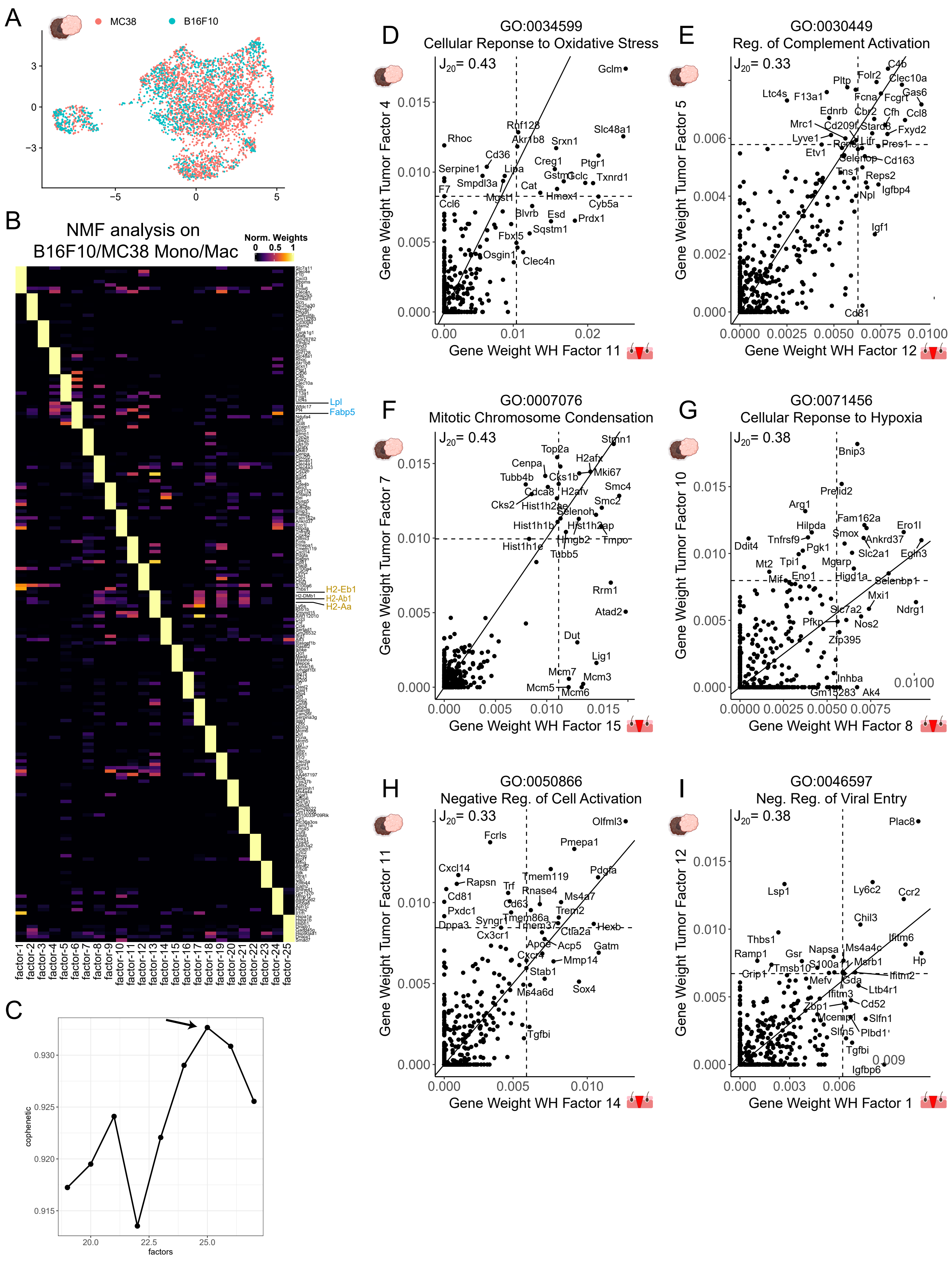


Supplementary Figure 6

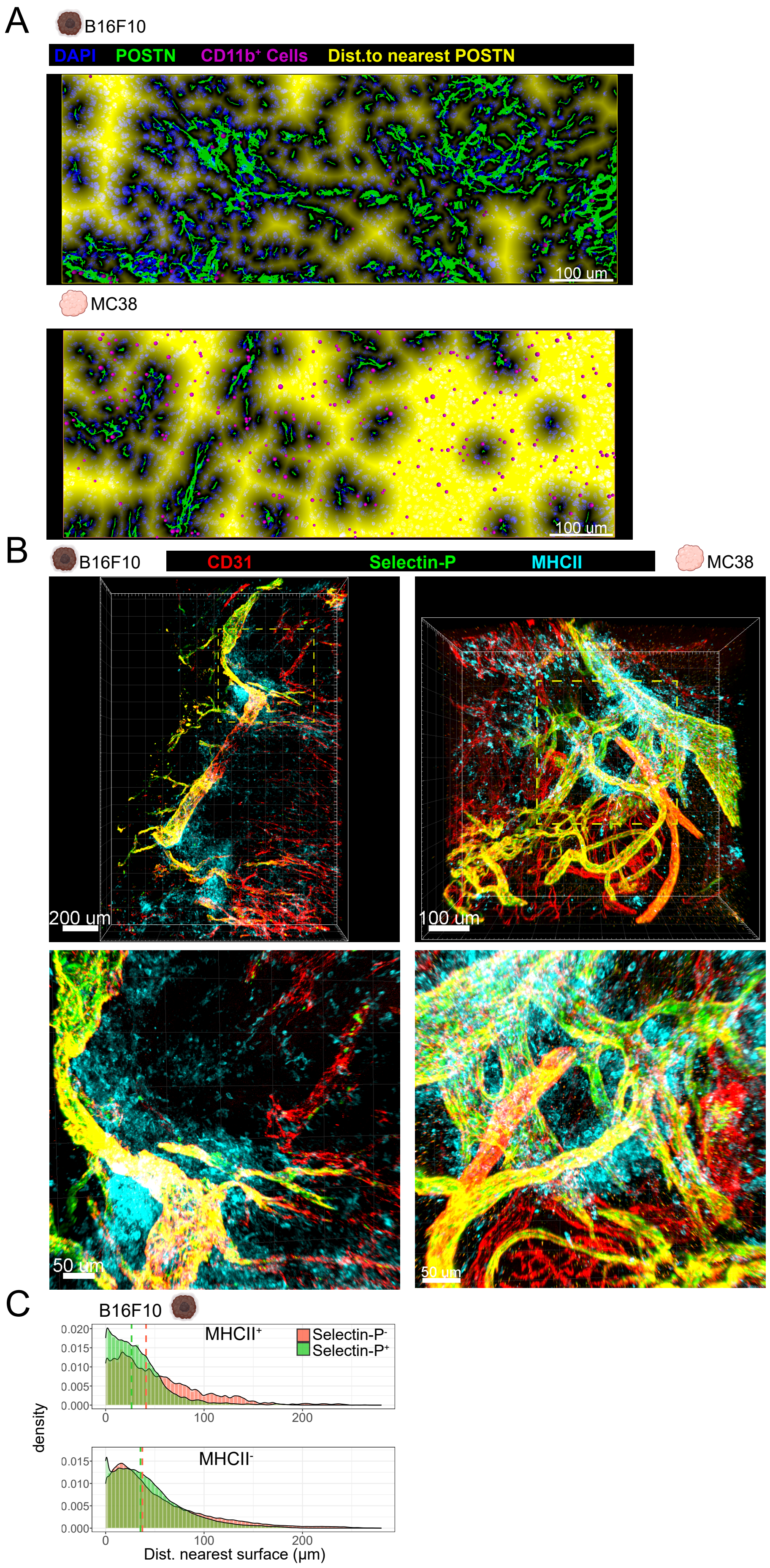


Supplementary Figure 7
