## Supplemental Table 1 for "Space-Time Mapping Identifies Concerted Multicellular Patterns and Gene Programs in Healing Wounds and their Conservation in Cancers"

| M_M factor-1 genes | M_M factor-1 weight | M_M factor-2 genes | M_M factor-2 weight | M_M factor-3 genes | M_M factor-3 weight | M_M factor-4 genes | M_M factor-4 weight | M_M factor-5 genes | M_M factor-5 weight | M_M factor-6 genes | M_M factor-6 weight |
| --- | --- | --- | --- | --- | --- | --- | --- | --- | --- | --- | --- |
| Hp | 0.010968 | Cd83 | 0.017718 | Col3a1 | 0.034269 | Cd72 | 0.011909 | 2810013P06Rik | 0.019583 | Ltc4s | 0.017004 |
| Plac8 | 0.010781 | Ccrl2 | 0.015186 | Col1a2 | 0.033362 | Clec12a | 0.010817 | Gm26917 | 0.017962 | Tppp3 | 0.014371 |
| Ifitm6 | 0.010004 | Id3 | 0.014934 | Sparc | 0.032426 | Axl | 0.009492 | Paxbp1 | 0.0171 | Padi4 | 0.013732 |
| Ccr2 | 0.009901 | Gadd45b | 0.014468 | Col1a1 | 0.031775 | Pou2f2 | 0.009213 | Gm47283 | 0.016 | Add3 | 0.012826 |
| Dmkn | 0.009442 | Nr4a1 | 0.013697 | Dcn | 0.023896 | Slamf9 | 0.00907 | Dleu2 | 0.014764 | Klf2 | 0.011319 |
| Cd177 | 0.009117 | Tgif1 | 0.013103 | Gm42418 | 0.021139 | Fnbp1l | 0.008821 | Mycbp2 | 0.01423 | Lair1 | 0.009968 |
| Vcan | 0.009095 | Pim3 | 0.012335 | Igfbp7 | 0.019184 | Calhm6 | 0.008686 | 4933433G15Rik | 0.014178 | Adgre5 | 0.009507 |
| Lrg1 | 0.008783 | Nfkbid | 0.012229 | Serpinh1 | 0.016516 | Cd74 | 0.008545 | Neat1 | 0.01351 | Anxa3 | 0.009208 |
| Igfbp6 | 0.008722 | Dusp2 | 0.011601 | Fstl1 | 0.0152 | H2-Ab1 | 0.008412 | Gm36486 | 0.012836 | Ncf1 | 0.00885 |
| Sirpb1c | 0.008184 | Btg2 | 0.011306 | S100a9 | 0.015159 | Tmem176b | 0.008356 | Lncpint | 0.012774 | Crip1 | 0.00877 |
| Ly6c2 | 0.008018 | Socs3 | 0.010954 | Rarres2 | 0.013969 | H2-Aa | 0.008211 | Mpp7 | 0.012293 | Smpdl3a | 0.008669 |
| Sell | 0.007861 | Kdm6b | 0.010909 | Col5a1 | 0.013717 | Pld4 | 0.008097 | Golgb1 | 0.01227 | Gas7 | 0.008605 |
| Nrg1 | 0.007622 | Atf3 | 0.010884 | Lgals7 | 0.013698 | H2-Eb1 | 0.008032 | 4933421O10Rik | 0.012206 | Naaa | 0.008573 |
| Slfn1 | 0.007599 | Nfil3 | 0.010773 | Col4a1 | 0.013246 | Ly86 | 0.008005 | Lars2 | 0.011777 | Gda | 0.00837 |
| Chil3 | 0.007276 | Egr1 | 0.010697 | Col6a1 | 0.013177 | Pltp | 0.007703 | Hspa1a | 0.011148 | Gpx1 | 0.008317 |
| Smim40 | 0.0072 | Ppp1r15a | 0.010661 | Cavin1 | 0.013036 | Ckb | 0.007696 | B930036N10Rik | 0.011032 | Tnfsf13 | 0.00825 |
| Ltb4r1 | 0.007168 | Csrnp1 | 0.010572 | Gpx3 | 0.012499 | Cx3cr1 | 0.007691 | Trim44 | 0.01067 | Lmna | 0.008089 |
| Ifitm2 | 0.006963 | Ier2 | 0.010555 | Aebp1 | 0.012492 | Aif1 | 0.007569 | 2900060B14Rik | 0.010607 | Slc28a2 | 0.00795 |
| Tgfbi | 0.006779 | Marcksl1 | 0.010403 | Mmp2 | 0.012295 | Cd79b | 0.007489 | Utrn | 0.010483 | Lsp1 | 0.007822 |
| Cd52 | 0.006708 | Tagap | 0.009976 | Gm26917 | 0.011429 | Clec4a3 | 0.007382 | Jun | 0.010483 | Ccl6 | 0.00758 |
| M_M factor-7 genes | M_M factor-7 weight | M_M factor-8 genes | M_M factor-8 weight | M_M factor-9 genes | M_M factor-9 weight | M_M factor-10 genes | M_M factor-10 weight | M_M factor-11 genes | M_M factor-11 weight | M_M factor-12 genes | M_M factor-12 weight |
| H2-DMa | 0.008367 | Ets1 | 0.010788 | Plau | 0.02181 | Itgal | 0.016702 | Gclm | 0.026282 | Gas6 | 0.009659 |
| H2-DMb1 | 0.008083 | Egln3 | 0.01041 | Sirt1 | 0.018098 | Stk10 | 0.012087 | Slc48a1 | 0.025937 | Ccl8 | 0.008786 |
| Cd209a | 0.008008 | Ndrg1 | 0.010098 | Atr | 0.014687 | Nxpe4 | 0.010871 | Ptgr1 | 0.02235 | Clec10a | 0.008621 |
| H2-Ab1 | 0.007825 | Impa2 | 0.009847 | Dhrs9 | 0.014482 | Ccdc88c | 0.010346 | Cyb5a | 0.022339 | C4b | 0.007878 |
| H2-Eb1 | 0.007669 | Ero1l | 0.009393 | Otud1 | 0.013902 | Spn | 0.009862 | Txnrd1 | 0.021561 | Fxyd2 | 0.007849 |
| Olfm1 | 0.007489 | Selenbp1 | 0.008528 | Cdk6 | 0.013689 | Gngt2 | 0.009145 | Gclc | 0.020429 | Cfh | 0.007724 |
| Ear2 | 0.007489 | Gys1 | 0.007771 | Bambi | 0.013632 | Itga4 | 0.009064 | Prdx1 | 0.018924 | Fcgrt | 0.007503 |
| H2-Aa | 0.007487 | Mxi1 | 0.007401 | Dot1l | 0.013516 | Rasgrp2 | 0.008946 | Gstm1 | 0.01731 | Pros1 | 0.007371 |
| Mgl2 | 0.00746 | Ankrd37 | 0.007262 | Dppa3 | 0.013514 | Adgre5 | 0.008808 | Hmox1 | 0.016342 | Igfbp4 | 0.00737 |
| Cd74 | 0.007204 | Fam162a | 0.00714 | Anxa1 | 0.013127 | Eno3 | 0.008566 | Srxn1 | 0.016214 | Folr2 | 0.00727 |
| Lmo1 | 0.007142 | Slc2a1 | 0.007096 | Myo1e | 0.012757 | Fgr | 0.008504 | Creg1 | 0.016013 | Igf1 | 0.007179 |
| Ramp1 | 0.007133 | Hk1 | 0.007009 | Asph | 0.012598 | Fam49a | 0.008456 | Esd | 0.015465 | Cbr2 | 0.007145 |
| Jaml | 0.007105 | Pgm1 | 0.007007 | Arf2 | 0.01212 | Cytip | 0.007968 | Abcc1 | 0.014414 | Stard8 | 0.007039 |
| Cd300lg | 0.006982 | Egln1 | 0.006978 | Rgs3 | 0.012052 | Ldlrad3 | 0.007963 | Cat | 0.01391 | Npl | 0.006777 |
| Vrk1 | 0.006864 | Nos2 | 0.006948 | Tsc22d3 | 0.01097 | Ceacam1 | 0.007925 | Blvrb | 0.012738 | Reps2 | 0.006705 |
| Cbfa2t3 | 0.006797 | Bnip3 | 0.006733 | Hsp90aa1 | 0.010798 | Samsn1 | 0.007603 | Sqstm1 | 0.012364 | Cd163 | 0.006644 |
| Mpp6 | 0.006765 | Ak4 | 0.006721 | Jund | 0.01009 | Pilrb2 | 0.007421 | Adgrl2 | 0.012056 | Cd81 | 0.006521 |
| Ank | 0.006625 | P4ha2 | 0.00662 | Ckap4 | 0.010013 | Pou2f2 | 0.007291 | Clec4n | 0.011441 | Tns1 | 0.006517 |
| Ciita | 0.006554 | Higd1a | 0.006533 | Snapc1 | 0.010006 | Adgre4 | 0.007255 | Rnf128 | 0.010717 | Lifr | 0.006498 |
| Lsp1 | 0.006506 | Mgarp | 0.006431 | Dpt | 0.009826 | St3gal5 | 0.007084 | Osgin1 | 0.010603 | Slco2b1 | 0.006386 |
| M_M factor-13 genes | M_M factor-13 weight | M_M factor-14 genes | M_M factor-14 weight | M_M factor-15 genes | M_M factor-15 weight | M_M factor-16 genes | M_M factor-16 weight | M_M factor-17 genes | M_M factor-17 weight | M_M factor-18 genes | M_M factor-18 weight |
| Rsad2 | 0.012069 | Itgb5 | 0.013205 | Atad2 | 0.017704 | Ccl24 | 0.012649 | Tmem8 | 0.014555 | Ezr | 0.008829 |
| Isg15 | 0.01204 | Olfml3 | 0.013185 | AI506816 | 0.017403 | Ccl7 | 0.011244 | Nxpe5 | 0.011567 | Car6 | 0.008645 |
| Ifit1 | 0.011741 | Itga6 | 0.012064 | Smc4 | 0.017074 | Ppbp | 0.010937 | Il4ra | 0.011319 | Cd9 | 0.007729 |
| Ifit3 | 0.01165 | Gatm | 0.011223 | Stmn1 | 0.016538 | Ccl2 | 0.010218 | Ap1b1 | 0.010372 | Ackr3 | 0.00771 |
| Ifi206 | 0.011527 | Pdgfa | 0.011179 | Rrm1 | 0.016251 | Tspan3 | 0.009991 | Mrc1 | 0.010369 | Errfi1 | 0.007479 |
| Phf11b | 0.011295 | Hexb | 0.010865 | Hist1h4d | 0.015461 | Pdia4 | 0.009163 | Nrp1 | 0.009934 | Cd24a | 0.007404 |
| Ifit2 | 0.010919 | Cxcl16 | 0.010683 | Ezh2 | 0.015449 | Ccl6 | 0.008783 | Rab3il1 | 0.009865 | Sema3c | 0.007223 |
| Oasl1 | 0.010829 | Tgfbr1 | 0.010059 | Smc2 | 0.015372 | Cd300lf | 0.008655 | Selenop | 0.009532 | Mmp19 | 0.007035 |
| Ifi209 | 0.010718 | Sox4 | 0.009793 | Tmpo | 0.015313 | Chil3 | 0.008264 | Rnase4 | 0.009216 | Cdk5r1 | 0.006766 |
| Usp18 | 0.010619 | F11r | 0.009782 | Lig1 | 0.014852 | Arg1 | 0.007917 | Ccl12 | 0.009155 | Hopx | 0.006693 |
| Ifi47 | 0.01041 | AU020206 | 0.009709 | Hist1h2ap | 0.014812 | 4930430E12Rik | 0.007667 | Cbr2 | 0.009065 | AA467197 | 0.006549 |
| Ifi213 | 0.010252 | Pmepa1 | 0.009452 | H2afx | 0.014267 | Lrp12 | 0.00752 | Stab1 | 0.008842 | Ncf1 | 0.006525 |
| Ms4a4c | 0.01014 | Lpcat2 | 0.008593 | Mcm3 | 0.013556 | E230016K23Rik | 0.007334 | Itsn1 | 0.008719 | Glipr2 | 0.006516 |
| Herc6 | 0.010124 | Ms4a7 | 0.008489 | Mcm6 | 0.013411 | Pf4 | 0.007105 | Gja1 | 0.008568 | Kcnn4 | 0.006425 |
| Phf11d | 0.010113 | Trem2 | 0.008323 | Mki67 | 0.013173 | Ccl9 | 0.00602 | Snx2 | 0.008098 | Serpine1 | 0.006361 |
| Cmpk2 | 0.010025 | Ctla2a | 0.008275 | Hmgb2 | 0.013131 | Timp1 | 0.005866 | Rgl1 | 0.00799 | Cd109 | 0.006258 |
| Gbp2 | 0.009969 | Mmp14 | 0.007939 | Dut | 0.012987 | Fnip2 | 0.005592 | Ap2a2 | 0.007915 | Rgcc | 0.006188 |
| Oas3 | 0.009934 | Apoc2 | 0.007868 | Mcm7 | 0.012164 | Slc7a8 | 0.005568 | Dab2 | 0.007784 | Adamtsl5 | 0.006179 |
| Ifi211 | 0.009452 | Cpd | 0.007757 | Mcm5 | 0.012133 | Ednrb | 0.005528 | Wwp1 | 0.007748 | Sgms1 | 0.006169 |
| Isg20 | 0.009413 | Tmem119 | 0.007739 | Tubb5 | 0.011875 | Vat1 | 0.005498 | Adgre1 | 0.007575 | Thbd | 0.006095 |
| M_M factor-19 genes | M_M factor-19 weight | M_M factor-20 genes | M_M factor-20 weight | M_M factor-21 genes | M_M factor-21 weight | M_M factor-22 genes | M_M factor-22 weight | M_M factor-23 genes | M_M factor-23 weight | M_M factor-24 genes | M_M factor-24 weight |
| Jag1 | 0.010752 | Ccnd2 | 0.011271 | Sdc1 | 0.009655 | Gpnmb | 0.014467 | Cd80 | 0.011064 | S100a8 | 0.024427 |
| Cxcl14 | 0.008784 | Ifitm1 | 0.010119 | C3 | 0.009135 | Pld3 | 0.012894 | B3gnt5 | 0.010668 | S100a9 | 0.023436 |
| Cxcl5 | 0.008618 | Spint1 | 0.009859 | Slpi | 0.009104 | Fabp5 | 0.011683 | Osm | 0.010202 | Retnlg | 0.016516 |
| Cited2 | 0.008559 | Il1r2 | 0.008573 | Clec5a | 0.00851 | Syngr1 | 0.011202 | AI504432 | 0.00963 | Csf1 | 0.014989 |
| Ctsk | 0.008075 | Fyn | 0.008244 | Fpr2 | 0.008276 | Lipa | 0.011154 | Olr1 | 0.009233 | Hdc | 0.014814 |
| Ctsl | 0.007965 | Gpr171 | 0.008166 | Ass1 | 0.008158 | Ahnak2 | 0.010766 | Gm48065 | 0.009085 | G0s2 | 0.014764 |
| Cxcl1 | 0.0078 | Plscr1 | 0.008016 | Acod1 | 0.007864 | Gdf15 | 0.010265 | Atp2b4 | 0.008107 | Cstdc4 | 0.012461 |
| Mt2 | 0.007666 | Grasp | 0.007835 | Carmil1 | 0.007764 | Trem2 | 0.008893 | Malt1 | 0.007996 | E230032D23Rik | 0.01242 |
| Cd36 | 0.007619 | Klrd1 | 0.007607 | Ptges | 0.007369 | Lpl | 0.008884 | Rab11fip1 | 0.007856 | Dgat1 | 0.012369 |
| Abca1 | 0.007522 | Ramp3 | 0.007501 | Sod2 | 0.007037 | Vat1 | 0.008831 | Slc7a11 | 0.007811 | Hcar2 | 0.011183 |
| Pf4 | 0.007464 | Avpi1 | 0.006955 | Fpr1 | 0.006718 | Anpep | 0.008469 | Dmxl2 | 0.007803 | Wfdc21 | 0.01117 |
| Myc | 0.007312 | Nr4a3 | 0.006817 | Tnip1 | 0.006644 | Cd63 | 0.008337 | Crem | 0.007517 | Ccl3 | 0.010986 |
| Arg1 | 0.00714 | Napsa | 0.006488 | Mmp14 | 0.006626 | Nceh1 | 0.008138 | Tgm2 | 0.007409 | Lcn2 | 0.009358 |
| Gadd45a | 0.007133 | Syngr2 | 0.006478 | Il1rn | 0.00655 | Mfsd12 | 0.00809 | Pde4d | 0.00709 | Ankrd33b | 0.009044 |
| Plpp3 | 0.006998 | Tspan13 | 0.006466 | Cav1 | 0.006536 | Atp6v0d2 | 0.007957 | Pde4b | 0.007021 | Igf1r | 0.008928 |
| Slc27a1 | 0.00664 | Traf1 | 0.006433 | Met | 0.00644 | Blnk | 0.007833 | Plk3 | 0.006956 | Ccl4 | 0.008662 |
| Sgk1 | 0.006583 | Klrk1 | 0.00641 | Il7r | 0.006429 | Angptl4 | 0.007697 | Gem | 0.006866 | Il1b | 0.008145 |
| Hmox1 | 0.00649 | Cytip | 0.006306 | Slc25a37 | 0.006368 | Serpinb6a | 0.007682 | Egr2 | 0.006863 | Acod1 | 0.008056 |
| Adrb2 | 0.006347 | Bcl2a1d | 0.006257 | Pcdh7 | 0.006321 | Creg1 | 0.007226 | Nfat5 | 0.006762 | Samsn1 | 0.007765 |
| Pgk1 | 0.006341 | H2-Ab1 | 0.006158 | Mirt2 | 0.006292 | Hist1h2bc | 0.007016 | Dusp5 | 0.006642 | Sh2d3c | 0.007717 |

| fibro factor-1 genes | fibro factor-1 weight | fibro factor-2 genes | fibro factor-2 weight | fibro factor-3 genes | fibro factor-3 weight | fibro factor-4 genes | fibro factor-4 weight | fibro factor-5 genes | fibro factor-5 weight | fibro factor-6 genes | fibro factor-6 weight |
| --- | --- | --- | --- | --- | --- | --- | --- | --- | --- | --- | --- |
| Serpinb2 | 0.011316 | Greb1 | 0.011412 | Sbsn | 0.014107 | Egr1 | 0.036707 | Lgals7 | 0.017012 | Cxcl14 | 0.013461 |
| Serpine1 | 0.010675 | Tmem158 | 0.010658 | Gas7 | 0.013959 | Fos | 0.036203 | Fxyd3 | 0.013212 | Mt1 | 0.012391 |
| Bnip3 | 0.010079 | Ogn | 0.01051 | Barx2 | 0.012483 | Fosb | 0.034848 | Apoc1 | 0.012847 | Ccl7 | 0.012019 |
| Glul | 0.009948 | Akr1c14 | 0.010053 | Anxa3 | 0.012339 | Ier2 | 0.032845 | Apoe | 0.012201 | Sod2 | 0.011462 |
| Mt2 | 0.009218 | Col6a6 | 0.009973 | Spon2 | 0.01033 | Btg2 | 0.031554 | S100a9 | 0.011664 | Pdpn | 0.011142 |
| Thbd | 0.009191 | Pgr | 0.009736 | Ccn3 | 0.01029 | Zfp36 | 0.031314 | Krt14 | 0.011622 | Aqp1 | 0.010803 |
| Acsl4 | 0.009148 | Gpx3 | 0.008711 | Rras2 | 0.009851 | Jun | 0.031042 | Krt5 | 0.010456 | Egfl7 | 0.010593 |
| Slc2a1 | 0.008993 | Itih5 | 0.008621 | Col5a3 | 0.009815 | Klf6 | 0.02623 | Cxcl2 | 0.010377 | Fabp4 | 0.010549 |
| Gfpt2 | 0.008901 | Mettl7a1 | 0.008431 | Tubb2b | 0.009723 | Ccn1 | 0.024326 | Hopx | 0.01018 | Mt2 | 0.010437 |
| Ptgis | 0.008752 | Ace | 0.008377 | Sfrp2 | 0.009426 | Dusp1 | 0.022171 | Gm42418 | 0.010007 | Abi3bp | 0.0103 |
| Apln | 0.008666 | Col14a1 | 0.008311 | Tppp3 | 0.009325 | Atf3 | 0.021215 | Net1 | 0.009713 | Timp1 | 0.009883 |
| Cxcl5 | 0.008663 | Ramp2 | 0.008213 | Creb5 | 0.009264 | Dnajb1 | 0.020836 | Dsp | 0.009701 | Mbd1 | 0.009849 |
| Uap1 | 0.008641 | Ngfr | 0.008141 | Hspb1 | 0.009121 | Gm34455 | 0.019533 | Fabp5 | 0.009558 | Mif | 0.009429 |
| Vegfa | 0.008555 | Celf4 | 0.007932 | Lrrn4cl | 0.008964 | Nfkbia | 0.019203 | Chka | 0.0088 | Rasl11a | 0.00915 |
| Fam162a | 0.008513 | Irf4 | 0.00785 | Pdlim1 | 0.00882 | Neat1 | 0.019192 | Mt1 | 0.008426 | Tspan13 | 0.008868 |
| Rdh10 | 0.008412 | Lrrn4cl | 0.007818 | Ackr3 | 0.008707 | Nfkbiz | 0.018908 | Fth1 | 0.008355 | Sdc4 | 0.008746 |
| Il33 | 0.00832 | Adam23 | 0.007494 | Cthrc1 | 0.00852 | Has1 | 0.016943 | Ly6d | 0.008272 | Tm4sf1 | 0.008609 |
| Slc39a14 | 0.0083 | Npy1r | 0.007475 | Glipr2 | 0.008391 | Nr4a1 | 0.016414 | S100a8 | 0.008266 | Pecam1 | 0.00851 |
| Rnd1 | 0.008219 | Prss23 | 0.00725 | Prss23 | 0.008233 | Pim1 | 0.016383 | Dapl1 | 0.008182 | Fth1 | 0.008486 |
| Sox11 | 0.008203 | Hmcn2 | 0.00709 | Wnt2 | 0.008081 | Gem | 0.01631 | Krt35 | 0.008173 | Ptx3 | 0.008412 |
| fibro factor-7 genes | fibro factor-7 weight | fibro factor-8 genes | fibro factor-8 weight | fibro factor-9 genes | fibro factor-9 weight | fibro factor-10 genes | fibro factor-10 weight | fibro factor-11 genes | fibro factor-11 weight | fibro factor-12 genes | fibro factor-12 weight |
| Dkk2 | 0.015258 | Tnc | 0.009826 | Cp | 0.012791 | Ptn | 0.012271 | Cxcl2 | 0.013198 | Ccl19 | 0.012372 |
| Gpm6b | 0.01431 | Acta2 | 0.009779 | Trf | 0.012183 | Postn | 0.011688 | Lyz2 | 0.013002 | Il1r2 | 0.010902 |
| Trps1 | 0.01316 | Nt5dc2 | 0.009661 | Apod | 0.011508 | Nrep | 0.010758 | Cd74 | 0.012351 | Flt3l | 0.010289 |
| Phldb2 | 0.012356 | Tpm2 | 0.009339 | Spp1 | 0.01144 | Spats2l | 0.010325 | S100a8 | 0.011847 | Sectm1a | 0.009826 |
| Tcf4 | 0.012327 | Runx1 | 0.009247 | Pdrg1 | 0.01047 | Col12a1 | 0.010056 | Pf4 | 0.011836 | Dapk1 | 0.009789 |
| Slc5a3 | 0.011564 | Tcim | 0.009109 | Rida | 0.010359 | Adam12 | 0.009952 | Il1b | 0.011692 | Cilp | 0.009731 |
| Gldn | 0.010668 | Ccn4 | 0.008811 | Sox9 | 0.009929 | Plaat3 | 0.009786 | S100a9 | 0.01149 | Dpep1 | 0.00972 |
| Nrep | 0.010627 | Pcsk5 | 0.008716 | Ebf2 | 0.009881 | Prss35 | 0.009726 | Srgn | 0.011218 | Pi16 | 0.009452 |
| Gas1 | 0.010433 | Crlf1 | 0.00852 | Cdkn2b | 0.009842 | Col27a1 | 0.009412 | Ccl4 | 0.011216 | Ecm2 | 0.009396 |
| Btbd3 | 0.010292 | Enpp1 | 0.008478 | Nr2f2 | 0.009477 | Mdk | 0.0093 | Fcer1g | 0.011182 | Mfap4 | 0.009086 |
| Enpp2 | 0.010235 | Cthrc1 | 0.008362 | Serpine2 | 0.009299 | H19 | 0.009202 | Arg1 | 0.010882 | Ltbp1 | 0.008786 |
| Syne1 | 0.00993 | Col5a3 | 0.007886 | Smim41 | 0.00896 | Lrrc15 | 0.008973 | Ctss | 0.010879 | Atp1a2 | 0.00873 |
| Coch | 0.009579 | Nr4a2 | 0.007822 | Gpc3 | 0.008889 | Gpc1 | 0.008937 | Tyrobp | 0.010703 | Bmp4 | 0.008587 |
| Slit2 | 0.009101 | Lox | 0.007806 | Col18a1 | 0.008412 | Cdh11 | 0.008713 | H2-Ab1 | 0.010094 | Sema3c | 0.00837 |
| Col4a1 | 0.008803 | Col12a1 | 0.007773 | Ptch1 | 0.008396 | Col7a1 | 0.00851 | Ccl6 | 0.010085 | Cd55 | 0.008307 |
| Nav3 | 0.008764 | Mmp19 | 0.007634 | Cldn1 | 0.008385 | Mmp11 | 0.008472 | H2-Aa | 0.009467 | Serpina3n | 0.008104 |
| Fbln5 | 0.008736 | Csrp2 | 0.007567 | Col28a1 | 0.008357 | Matn2 | 0.008451 | Wfdc17 | 0.009407 | Gas6 | 0.00809 |
| Igsf10 | 0.008657 | Lrrc32 | 0.007443 | Rasgrp2 | 0.008277 | Spon1 | 0.00842 | Laptm5 | 0.009346 | AW112010 | 0.008033 |
| Cpxm2 | 0.008573 | Timp1 | 0.007302 | Col8a1 | 0.008145 | Brinp3 | 0.008334 | H2-Eb1 | 0.009146 | Pcolce2 | 0.008033 |
| F2r | 0.00842 | Tnfrsf12a | 0.007263 | P2ry14 | 0.008025 | Aspn | 0.008067 | Ccl3 | 0.009081 | Htra3 | 0.008027 |
| fibro factor-13 genes | fibro factor-13 weight | fibro factor-14 genes | fibro factor-14 weight | fibro factor-15 genes | fibro factor-15 weight | fibro factor-16 genes | fibro factor-16 weight | fibro factor-17 genes | fibro factor-17 weight | | |
| Ppp1r14a | 0.010096 | Sned1 | 0.012659 | Mef2c | 0.011906 | Gdf10 | 0.014919 | Smc2 | 0.014658 |  |  |
| Cldn10 | 0.01001 | Vegfd | 0.012525 | Rflnb | 0.010488 | F3 | 0.014529 | Birc5 | 0.013637 |  |  |
| Cdh4 | 0.009907 | Mmp3 | 0.012364 | Col11a1 | 0.010427 | Mgp | 0.013103 | Hmgb2 | 0.013458 |  |  |
| Miat | 0.009704 | Ggt5 | 0.012059 | Ptch1 | 0.010072 | Cygb | 0.012285 | Mki67 | 0.013242 |  |  |
| Cpz | 0.009662 | Cxcl12 | 0.011984 | Tns3 | 0.010061 | Epha3 | 0.011854 | Pclaf | 0.013208 |  |  |
| Aff3 | 0.009651 | C3 | 0.011643 | Nav2 | 0.009768 | Tspan11 | 0.01151 | Cks2 | 0.013059 |  |  |
| Tmem132c | 0.009427 | Tmem176a | 0.011558 | Pde3a | 0.009562 | C2 | 0.011482 | Smc4 | 0.012635 |  |  |
| Apcdd1 | 0.009412 | Tmem176b | 0.011221 | Myl9 | 0.009458 | Rbp1 | 0.011452 | Tacc3 | 0.012577 |  |  |
| Sema3b | 0.009263 | C4b | 0.011202 | Mgst3 | 0.009098 | Ccdc3 | 0.011179 | Rrm1 | 0.012225 |  |  |
| Cyp2f2 | 0.009214 | Cyp7b1 | 0.011035 | Stmn2 | 0.009042 | Cfh | 0.010543 | Cenpa | 0.012159 |  |  |
| Wnt5a | 0.009185 | Steap4 | 0.010931 | Slc6a6 | 0.008913 | Nr2f2 | 0.009958 | Cdca8 | 0.01208 |  |  |
| Oxtr | 0.009067 | Serpina3n | 0.010834 | Sox18 | 0.008903 | Fmo2 | 0.009868 | Stmn1 | 0.012038 |  |  |
| Pla2g2e | 0.00902 | Cfb | 0.01072 | Wif1 | 0.008816 | Meox2 | 0.009492 | Tpx2 | 0.01195 |  |  |
| Scara5 | 0.008944 | Cpxm1 | 0.010674 | Bpgm | 0.008553 | Zim1 | 0.009346 | Anln | 0.011948 |  |  |
| Pi15 | 0.008943 | Cyp1b1 | 0.010291 | Ntrk3 | 0.008344 | Chrdl1 | 0.009259 | Cdca3 | 0.011569 |  |  |
| Pcolce2 | 0.008927 | Plac8 | 0.009915 | Alx4 | 0.008305 | Igfbp4 | 0.009092 | Top2a | 0.011564 |  |  |
| Aldh3a1 | 0.008809 | Ifit1 | 0.009512 | Fgl2 | 0.008298 | Pltp | 0.008979 | Racgap1 | 0.011543 |  |  |
| Slc27a6 | 0.008688 | Lgi2 | 0.009138 | Sms | 0.008274 | Peg3 | 0.008906 | H2afx | 0.011514 |  |  |
| Igfbp5 | 0.00857 | Scn7a | 0.009126 | Adamts18 | 0.00825 | Dact1 | 0.008794 | Cdk1 | 0.011373 |  |  |
| Qpct | 0.008506 | Slfn5 | 0.009076 | F2r | 0.008145 | Id4 | 0.008651 | Diaph3 | 0.011367 |  |  |

| Kerat factor-1 genes | Kerat factor-1 weight | Kerat factor-2 genes | Kerat factor-2 weight | Kerat factor-3 genes | Kerat factor-3 weight | Kerat factor-4 genes | Kerat factor-4 weight | Kerat factor-5 genes | Kerat factor-5 weight | Kerat factor-6 genes | Kerat factor-6 weight |
| --- | --- | --- | --- | --- | --- | --- | --- | --- | --- | --- | --- |
| Kitl | 0.010871 | Cst6 | 0.007771 | Dapl1 | 0.009138 | Krt10 | 0.012039 | Mki67 | 0.010169 | Akr1c18 | 0.007876 |
| Gli3 | 0.00997 | Defb6 | 0.00766 | Them4 | 0.008659 | Il6ra | 0.00774 | Top2a | 0.009673 | Srebf1 | 0.007041 |
| Slc16a10 | 0.009699 | Krt79 | 0.007485 | Celsr1 | 0.00837 | Stfa3 | 0.007296 | Cks2 | 0.009636 | Oxtr | 0.007003 |
| Prr5l | 0.007836 | Slc43a2 | 0.006838 | Lap3 | 0.007725 | Slco3a1 | 0.006892 | Cenpa | 0.009408 | Ar | 0.006874 |
| Prdm1 | 0.007821 | Alox12e | 0.006635 | Mtcl1 | 0.00741 | Hlf | 0.006602 | Cdca8 | 0.009402 | Avpr1a | 0.006741 |
| Slc7a5 | 0.007791 | Pdzk1ip1 | 0.0065 | Ptpre | 0.007173 | Chl1 | 0.006445 | Incenp | 0.009288 | Tgfbr3 | 0.006702 |
| Foxq1 | 0.007714 | Skint3 | 0.006185 | Gm49425 | 0.007099 | Lgals3 | 0.006313 | Cenpf | 0.009264 | Igfbp2 | 0.00669 |
| Krt75 | 0.007562 | Fbxo32 | 0.00616 | Cybrd1 | 0.007083 | Serpinb2 | 0.006202 | Birc5 | 0.009053 | Fgfbp1 | 0.006601 |
| 4930523C07Rik | 0.00715 | Nebl | 0.006134 | Tagln3 | 0.00707 | Ndufa4l2 | 0.00616 | Lockd | 0.009052 | Steap4 | 0.006527 |
| Tspan3 | 0.007126 | Klk7 | 0.006114 | Krt36 | 0.006857 | S100a11 | 0.00615 | Cdca3 | 0.008854 | F730043M19Rik | 0.006491 |
| Zfhx3 | 0.007104 | Cstb | 0.005831 | Blmh | 0.006815 | Krt1 | 0.006064 | Prc1 | 0.008685 | Psph | 0.006211 |
| Aldh1a3 | 0.00707 | Susd2 | 0.005822 | Pxylp1 | 0.006746 | Limch1 | 0.00599 | Kif20b | 0.008527 | Plin2 | 0.006167 |
| Gpnmb | 0.007068 | Ly6g6c | 0.005821 | Ttll7 | 0.006744 | Trim7 | 0.005939 | Knl1 | 0.008439 | Dag1 | 0.006145 |
| Dct | 0.007037 | Atp6v1c2 | 0.005682 | Selenbp1 | 0.006715 | Neat1 | 0.005905 | Racgap1 | 0.008437 | Ldhb | 0.006106 |
| Cyfip2 | 0.006975 | Plet1 | 0.00556 | Bicdl2 | 0.006701 | Sema3c | 0.005748 | Ncapd2 | 0.008414 | Prxl2a | 0.006101 |
| Gpc3 | 0.006502 | Pgap1 | 0.005489 | Alox12 | 0.006566 | Dmkn | 0.005684 | Smc2 | 0.008311 | Lama4 | 0.006093 |
| Krt23 | 0.006258 | Elovl7 | 0.005455 | Krt86 | 0.006512 | Krtdap | 0.005643 | Cenpe | 0.008305 | Slc16a1 | 0.006054 |
| Jakmip2 | 0.006177 | Bbox1 | 0.00545 | Hspa2 | 0.006492 | Oas1f | 0.005619 | Tpx2 | 0.008305 | Rgcc | 0.006032 |
| Tgm6 | 0.006064 | Psapl1 | 0.005334 | Fam25c | 0.006489 | Urah | 0.005534 | H2afx | 0.008303 | Hmgcs2 | 0.005985 |
| Syt7 | 0.006046 | Defb1 | 0.005276 | Trps1 | 0.00646 | Lgals7 | 0.005476 | Tacc3 | 0.008299 | Acsbg1 | 0.005914 |
| Kerat factor-7 genes | Kerat factor-7 weight | Kerat factor-8 genes | Kerat factor-8 weight | Kerat factor-9 genes | Kerat factor-9 weight | Kerat factor-10 genes | Kerat factor-10 weight | Kerat factor-11 genes | Kerat factor-11 weight | Kerat factor-12 genes | Kerat factor-12 weight |
| Ifitm3 | 0.007733 | Gm973 | 0.007951 | Col1a1 | 0.01266 | Col4a1 | 0.006069 | Jag1 | 0.009682 | Kalrn | 0.007192 |
| Fst | 0.007364 | Postn | 0.00735 | Col3a1 | 0.012311 | Timp3 | 0.005893 | St14 | 0.009359 | Dclk1 | 0.006835 |
| H2-Q7 | 0.006893 | Hmcn1 | 0.006968 | Vim | 0.012286 | Fndc1 | 0.005842 | Sat1 | 0.008437 | Mgat4a | 0.006652 |
| Sostdc1 | 0.006882 | Sdk2 | 0.006367 | Col1a2 | 0.011631 | Col4a2 | 0.005636 | Notch1 | 0.008027 | Insig2 | 0.006578 |
| Ccl27a | 0.006389 | Rflnb | 0.006337 | Srgn | 0.011319 | Fblim1 | 0.005554 | Stk17b | 0.007952 | Pthlh | 0.006534 |
| Lgals9 | 0.006351 | Fzd2 | 0.006068 | Cd74 | 0.011188 | Cbs | 0.005488 | Rnaset2a | 0.007845 | Fam102b | 0.006523 |
| Ahr | 0.006256 | Lhx2 | 0.005976 | Cxcl2 | 0.010821 | Specc1 | 0.005436 | Epha2 | 0.007829 | Myof | 0.006479 |
| H2-Q4 | 0.006088 | Ank | 0.005912 | S100a9 | 0.009896 | Col16a1 | 0.00515 | Spint1 | 0.007697 | Ece1 | 0.006177 |
| Adgrl2 | 0.005968 | Itprid2 | 0.005803 | Fcer1g | 0.009763 | Slc7a1 | 0.005148 | Wnt10b | 0.007653 | Pfn2 | 0.005982 |
| Ackr4 | 0.005954 | Dkk3 | 0.005787 | H2-Aa | 0.009589 | Igfbp7 | 0.005097 | Pitrm1 | 0.007616 | Efna5 | 0.00589 |
| Aldh3a1 | 0.005904 | Adgrg6 | 0.005779 | Pf4 | 0.009384 | Angptl4 | 0.00506 | Pcdh19 | 0.007489 | Lypd3 | 0.005821 |
| Adh7 | 0.005835 | Fgfr1 | 0.005649 | H2-Ab1 | 0.009375 | Myl9 | 0.005046 | Rnaset2b | 0.007298 | Ccnd1 | 0.005797 |
| Txnip | 0.005823 | Ism1 | 0.005469 | Il1b | 0.008901 | Col18a1 | 0.004986 | Sox21 | 0.007111 | Pdzrn3 | 0.00578 |
| Psmb9 | 0.005773 | Ptn | 0.00545 | Tyrobp | 0.008851 | Cpxm1 | 0.004875 | Sec11c | 0.006915 | AW011738 | 0.005764 |
| Arl4c | 0.005719 | Filip1l | 0.005424 | S100a8 | 0.008806 | Cbr2 | 0.004816 | Hoxc13 | 0.006894 | Sdc1 | 0.005722 |
| Cdh13 | 0.005688 | Phlda1 | 0.005382 | H2-Eb1 | 0.008557 | Gnmt | 0.004731 | Krt35 | 0.0066 | Myo1b | 0.005433 |
| Krt15 | 0.005659 | Cited2 | 0.005371 | Ctss | 0.008494 | Lgr5 | 0.00473 | Rexo2 | 0.006598 | Cmah | 0.005327 |
| H2-Q6 | 0.005635 | Col7a1 | 0.005326 | Lyz2 | 0.008402 | Slc39a14 | 0.004708 | Eif2s2 | 0.006325 | Sbsn | 0.005325 |
| Psmb8 | 0.00557 | Shisa2 | 0.005271 | Cd52 | 0.008333 | Itga1 | 0.004697 | Unc5b | 0.006207 | Iffo2 | 0.005271 |
| Cxadr | 0.005454 | Col17a1 | 0.005265 | S100a4 | 0.008205 | Il11ra1 | 0.004594 | Plekhg1 | 0.006124 | Slc35g1 | 0.005246 |
| Kerat factor-13 genes | Kerat factor-13 weight | Kerat factor-14 genes | Kerat factor-14 weight | Kerat factor-15 genes | Kerat factor-15 weight | Kerat factor-16 genes | Kerat factor-16 weight | | |  |  |
| Krt25 | 0.00953 | Elovl6 | 0.009187 | Ier2 | 0.023867 | Pcna | 0.015294 |  |  |  |  |
| Krt28 | 0.009311 | Tmem109 | 0.008272 | Jun | 0.023708 | Mcm4 | 0.013903 |  |  |  |  |
| Paqr5 | 0.008903 | Secisbp2l | 0.008254 | Egr1 | 0.023305 | Hells | 0.013785 |  |  |  |  |
| Taf13 | 0.008766 | Dhrs7 | 0.007759 | Hspa1a | 0.023161 | Mcm6 | 0.013393 |  |  |  |  |
| Gata3 | 0.00825 | Dhcr24 | 0.007655 | Junb | 0.022883 | Mcm3 | 0.01208 |  |  |  |  |
| Dsc1 | 0.007941 | Slc25a1 | 0.00765 | Hspa1b | 0.02029 | Dtl | 0.011891 |  |  |  |  |
| Krt71 | 0.007939 | Paqr7 | 0.007578 | Jund | 0.019237 | Ung | 0.011315 |  |  |  |  |
| Krt73 | 0.007829 | Acss2 | 0.007575 | Btg2 | 0.018809 | Mcm5 | 0.01114 |  |  |  |  |
| Csgalnact1 | 0.007826 | Hacd2 | 0.007475 | Fos | 0.0174 | Dut | 0.010916 |  |  |  |  |
| Arhgap29 | 0.007603 | Scd1 | 0.00731 | Mif | 0.016221 | Lig1 | 0.010545 |  |  |  |  |
| Prnp | 0.007598 | Lpcat3 | 0.00722 | Ier3 | 0.016112 | Atad5 | 0.010398 |  |  |  |  |
| Krt27 | 0.007549 | Far2 | 0.00718 | Nfkbia | 0.015935 | Atad2 | 0.01039 |  |  |  |  |
| Cdh6 | 0.0075 | Apmap | 0.007064 | Zfp36 | 0.015556 | Nsd2 | 0.010032 |  |  |  |  |
| Ppfibp1 | 0.007441 | Cidea | 0.00686 | Urah | 0.014749 | Syce2 | 0.009712 |  |  |  |  |
| Dok4 | 0.007364 | Fitm2 | 0.006827 | Tspo | 0.01356 | Gmnn | 0.009432 |  |  |  |  |
| Wdr47 | 0.007272 | Hsd17b12 | 0.006766 | Klf6 | 0.013179 | Uhrf1 | 0.008898 |  |  |  |  |
| Nudt13 | 0.007231 | Pdss1 | 0.006675 | Fam162a | 0.013147 | Siva1 | 0.008408 |  |  |  |  |
| Gm49708 | 0.007196 | Echdc1 | 0.006653 | Fosb | 0.01262 | Stmn1 | 0.008321 |  |  |  |  |
| Nkd2 | 0.007129 | Cers4 | 0.006535 | Myl6 | 0.012489 | Ccne2 | 0.008255 |  |  |  |  |
| Notch2 | 0.007043 | Aacs | 0.006512 | Rhob | 0.011579 | Cbx5 | 0.00822 |  |  |  |  |

| Endo factor-1 genes | Endo factor-1 weight | Endo factor-2 genes | Endo factor-2 weight | Endo factor-3 genes | Endo factor-3 weight | Endo factor-4 genes | Endo factor-4 weight | Endo factor-5 genes | Endo factor-5 weight | Endo factor-6 genes | Endo factor-6 weight |
| --- | --- | --- | --- | --- | --- | --- | --- | --- | --- | --- | --- |
| Ptpn14 | 0.007651 | Slc39a10 | 0.009327 | Mcam | 0.007361 | Tcf15 | 0.008553 | Col3a1 | 0.011178 | Sema3g | 0.009363 |
| Fxyd6 | 0.007497 | Pcp4l1 | 0.008624 | Mest | 0.006785 | Lpl | 0.007543 | Col1a1 | 0.010567 | Stmn2 | 0.008924 |
| Add3 | 0.007457 | Slc30a1 | 0.007057 | Prnp | 0.006301 | Ablim3 | 0.007148 | Col1a2 | 0.010193 | Fbln5 | 0.007547 |
| Ahnak2 | 0.007358 | Mir100hg | 0.006905 | Col18a1 | 0.006133 | Fmo2 | 0.006766 | Dcn | 0.007599 | Gja4 | 0.007485 |
| Hmcn1 | 0.007247 | H2-Q6 | 0.006778 | Fscn1 | 0.00611 | C1qtnf9 | 0.006153 | Gsn | 0.007373 | Cdk19 | 0.007074 |
| Tmod2 | 0.007105 | Pltp | 0.006739 | Kit | 0.006099 | Fmo1 | 0.006082 | Lgals1 | 0.007338 | Jag1 | 0.006796 |
| Klhl4 | 0.006646 | Ahr | 0.006597 | Chst1 | 0.006002 | Afdn | 0.005999 | Meg3 | 0.007052 | Aggf1 | 0.006794 |
| Nsg1 | 0.006607 | Iigp1 | 0.006077 | Nid2 | 0.005746 | Itga1 | 0.005955 | Col5a2 | 0.006815 | Efnb2 | 0.006711 |
| Stab1 | 0.006456 | Sned1 | 0.005925 | Col4a1 | 0.005561 | Ccdc85a | 0.00587 | Col6a3 | 0.006624 | Atp13a3 | 0.006612 |
| Ntn1 | 0.006424 | Hoxd9 | 0.005846 | Col4a2 | 0.005522 | Gpihbp1 | 0.005779 | Timp1 | 0.006585 | Azin1 | 0.006541 |
| Cpne2 | 0.006258 | Lrat | 0.005708 | Map1b | 0.005466 | Nrp1 | 0.005743 | Col5a1 | 0.006515 | Amd1 | 0.006321 |
| Prkcz | 0.00592 | Fam13c | 0.005636 | Mmp14 | 0.005461 | Tspan13 | 0.005688 | Gpx3 | 0.00642 | Clu | 0.006305 |
| Palm | 0.005903 | Apcdd1 | 0.005634 | Vwa1 | 0.005426 | Slc28a2 | 0.005668 | Acta2 | 0.006407 | Atp2a3 | 0.006157 |
| Flt4 | 0.005822 | Nfat5 | 0.005625 | Rbp1 | 0.005324 | Timp4 | 0.005566 | Rgs5 | 0.00633 | Bmx | 0.006085 |
| Arrdc3 | 0.005819 | Smad1 | 0.005612 | Prnd | 0.005312 | Cd36 | 0.005552 | Postn | 0.006142 | Glul | 0.005992 |
| Itga9 | 0.005802 | Itih5 | 0.005603 | N4bp3 | 0.005308 | Cavin2 | 0.005493 | Pcolce | 0.006137 | Egfl8 | 0.005876 |
| Cp | 0.005796 | Sparcl1 | 0.005307 | Apln | 0.005218 | Lims2 | 0.005435 | Col5a3 | 0.00609 | Jag2 | 0.005841 |
| Ackr3 | 0.005744 | Eogt | 0.005293 | Cd276 | 0.005209 | Sdc3 | 0.005299 | Col6a2 | 0.006068 | Lmo2 | 0.005837 |
| Slc41a1 | 0.005653 | Nin | 0.005282 | Adamts4 | 0.005164 | Aqp7 | 0.005292 | Tpm2 | 0.006048 | Cldn5 | 0.005757 |
| Gpm6a | 0.005638 | Gpcpd1 | 0.005201 | Sox4 | 0.005083 | Pcdh19 | 0.005213 | Cd63 | 0.005986 | Ptprr | 0.005739 |
| Endo factor-7 genes | Endo factor-7 weight | Endo factor-8 genes | Endo factor-8 weight | Endo factor-9 genes | Endo factor-9 weight | Endo factor-10 genes | Endo factor-10 weight | Endo factor-11 genes | Endo factor-11 weight | | |
| Stmn1 | 0.009603 | Selp | 0.007182 | Rbp7 | 0.009067 | Junb | 0.012482 | Cxcl2 | 0.012479 |  |  |
| Hmgb2 | 0.009144 | Vwf | 0.006726 | Btnl9 | 0.008452 | Jund | 0.011889 | S100a9 | 0.009598 |  |  |
| Lmnb1 | 0.008376 | Ptgs1 | 0.006725 | Gask1b | 0.007213 | Jun | 0.01133 | Cd74 | 0.009531 |  |  |
| Cks2 | 0.007875 | Lrg1 | 0.006634 | Cxcl12 | 0.006864 | Hspb1 | 0.01108 | Ccl4 | 0.009036 |  |  |
| Smc2 | 0.007797 | Il6st | 0.006632 | Mgll | 0.006673 | Ier3 | 0.009102 | S100a8 | 0.008962 |  |  |
| Mki67 | 0.007657 | Ctnnal1 | 0.006424 | Angptl4 | 0.006477 | Cebpd | 0.009043 | Lyz2 | 0.008893 |  |  |
| Cenpa | 0.007634 | Slco2a1 | 0.006411 | Gpihbp1 | 0.006371 | Socs3 | 0.00898 | Spp1 | 0.008575 |  |  |
| Hmgn2 | 0.007106 | Ackr1 | 0.00638 | Pcdh17 | 0.006314 | 2410006H16Rik | 0.008697 | Fcer1g | 0.008272 |  |  |
| Top2a | 0.007044 | Ehd4 | 0.006231 | Cd36 | 0.006222 | Ier2 | 0.008639 | Ctss | 0.007834 |  |  |
| Lig1 | 0.006972 | Chp2 | 0.006065 | Cd300lg | 0.00606 | Gm10076 | 0.008594 | Pf4 | 0.007823 |  |  |
| H2afx | 0.006932 | Ehd3 | 0.006008 | Plpp1 | 0.006057 | Gapdh | 0.008294 | Il1b | 0.007722 |  |  |
| Rrm1 | 0.006796 | Aqp1 | 0.005824 | Kctd12b | 0.00603 | Eif5a | 0.008122 | Cd52 | 0.007699 |  |  |
| Incenp | 0.006727 | Rasa4 | 0.005745 | Plpp3 | 0.005939 | Cycs | 0.00778 | Cxcl3 | 0.007679 |  |  |
| Ncapd2 | 0.006639 | Pdia5 | 0.005654 | Dysf | 0.005869 | Ifitm2 | 0.00771 | Tyrobp | 0.007586 |  |  |
| Ccdc34 | 0.006595 | Spint2 | 0.005643 | Rgcc | 0.005806 | Mif | 0.007701 | H2-Aa | 0.007316 |  |  |
| Cdk1 | 0.006594 | Cysltr1 | 0.005615 | Kdr | 0.005724 | Ppp1r15a | 0.007635 | Ccl3 | 0.007309 |  |  |
| Cks1b | 0.006567 | Cmah | 0.005552 | Efr3b | 0.005695 | Ran | 0.007454 | H2-Ab1 | 0.007275 |  |  |
| Selenoh | 0.006553 | Sncg | 0.005541 | Thbd | 0.005543 | Il6 | 0.007441 | Arg1 | 0.00726 |  |  |
| Birc5 | 0.006537 | Il1r1 | 0.00552 | Mef2c | 0.005336 | Btg2 | 0.007369 | Wfdc17 | 0.006866 |  |  |
| Smc4 | 0.006504 | Dnm3 | 0.005484 | Ccnd1 | 0.005259 | Fos | 0.007348 | Ptgs2 | 0.006827 |  |  |

| T factor-1 genes | T factor-1 weight | T factor-2 genes | T factor-2 weight | T factor-3 genes | T factor-3 weight | T factor-4 genes | T factor-4 weight | T factor-5 genes | T factor-5 weight | T factor-6 genes | T factor-6 weight |
| --- | --- | --- | --- | --- | --- | --- | --- | --- | --- | --- | --- |
| Mki67 | 0.008697 | Tmem176a | 0.006847 | C1qa | 0.005287 | Sparc | 0.007552 | Cxcl2 | 0.008766 | Nfkbid | 0.006128 |
| Stmn1 | 0.008414 | Serpinb1a | 0.006698 | Grn | 0.004984 | Col1a2 | 0.006876 | Thbs1 | 0.007982 | Neurl3 | 0.005989 |
| Incenp | 0.008226 | Cd163l1 | 0.006583 | Lrp1 | 0.004892 | Dcn | 0.006547 | Il1b | 0.007404 | Tnfaip3 | 0.005903 |
| Top2a | 0.008095 | Cxcr6 | 0.006362 | Mrc1 | 0.004864 | Apoe | 0.006209 | S100a8 | 0.007025 | Furin | 0.00573 |
| Cks2 | 0.007974 | Actn2 | 0.00636 | Aif1 | 0.004863 | Mt2 | 0.006068 | S100a9 | 0.007013 | Zfp36l2 | 0.00557 |
| Tubb4b | 0.007871 | Blk | 0.006295 | F13a1 | 0.004785 | Mt1 | 0.006036 | Ptgs2 | 0.006987 | Nr4a3 | 0.00542 |
| Hmgn2 | 0.007775 | Slc7a6os | 0.006116 | Csf1r | 0.004764 | Igfbp7 | 0.005558 | Ccl3 | 0.00689 | Ar | 0.005299 |
| Lig1 | 0.007631 | Tmem176b | 0.006103 | C1qb | 0.004704 | Timp1 | 0.005485 | Clec4e | 0.006659 | Itk | 0.005043 |
| Smc2 | 0.0076 | Aqp3 | 0.005965 | Unc93b1 | 0.004619 | Fabp4 | 0.005471 | Ier3 | 0.006607 | Cd40lg | 0.004988 |
| Dut | 0.007417 | Il18r1 | 0.005779 | Stab1 | 0.004614 | Lgals7 | 0.005457 | Slpi | 0.006545 | Gm15472 | 0.004852 |
| Mcm3 | 0.007257 | Mmp25 | 0.005741 | Cybb | 0.004605 | Serpinh1 | 0.005397 | Ccl4 | 0.006532 | Gadd45b | 0.004846 |
| Tubb5 | 0.007249 | Abi3bp | 0.005505 | Tmem106a | 0.004584 | Fstl1 | 0.005384 | Il1rn | 0.006384 | Gata3 | 0.004823 |
| Cks1b | 0.007234 | Rorc | 0.005347 | Mafb | 0.004566 | Col1a1 | 0.005327 | Acod1 | 0.006358 | Uhrf2 | 0.004802 |
| Racgap1 | 0.007232 | Il7r | 0.005237 | Mpeg1 | 0.004519 | Aebp1 | 0.005264 | Cxcl3 | 0.006187 | Nr4a1 | 0.004727 |
| Hist1h2ap | 0.007139 | Tmem64 | 0.005233 | Zeb2 | 0.004488 | Cald1 | 0.005182 | Arg1 | 0.006145 | Hlf | 0.00468 |
| Tuba1b | 0.007044 | Jaml | 0.005146 | Lgmn | 0.004485 | S100a9 | 0.005049 | Spp1 | 0.006072 | Il2 | 0.004587 |
| H2afx | 0.007018 | Ltb4r1 | 0.005124 | Sdc3 | 0.004469 | Ctsb | 0.004984 | Plaur | 0.005844 | Elmsan1 | 0.004557 |
| Hmgb2 | 0.006976 | Dap | 0.005039 | C1qc | 0.004382 | Serpinf1 | 0.004968 | Nlrp3 | 0.005771 | Dusp5 | 0.004472 |
| Lockd | 0.006915 | St3gal6 | 0.004881 | Lyz2 | 0.004373 | Gja1 | 0.004944 | Cebpb | 0.005761 | Bhlhe40 | 0.004435 |
| Ccdc34 | 0.00689 | Ramp1 | 0.004856 | Fcgr1 | 0.004351 | Nfix | 0.004905 | Osm | 0.005756 | Itpkb | 0.004369 |
| T factor-7 genes | T factor-7 weight | T factor-8 genes | T factor-8 weight | T factor-9 genes | T factor-9 weight | T factor-10 genes | T factor-10 weight | |  |  |  |
| Cd7 | 0.007075 | Ctla4 | 0.008315 | Ifi47 | 0.00923 | H2-Eb1 | 0.00755 |  |  |  |  |
| Ctla2b | 0.00675 | Ikzf2 | 0.008232 | Gimap3 | 0.008669 | H2-Aa | 0.006799 |  |  |  |  |
| Nkg7 | 0.006652 | Tnfrsf4 | 0.007999 | Gimap6 | 0.008242 | H2-DMa | 0.006769 |  |  |  |  |
| Prf1 | 0.005934 | Arl5a | 0.007798 | Ms4a6b | 0.007298 | H2-Ab1 | 0.006661 |  |  |  |  |
| Xcl1 | 0.005903 | Foxp3 | 0.007037 | Gimap4 | 0.007288 | H2-DMb1 | 0.006582 |  |  |  |  |
| Prkacb | 0.005834 | Il2ra | 0.006757 | Ms4a4b | 0.007021 | Napsa | 0.006498 |  |  |  |  |
| Avil | 0.005792 | Itgav | 0.006636 | S1pr1 | 0.006988 | Ciita | 0.006464 |  |  |  |  |
| Ctla2a | 0.005753 | Il1rl1 | 0.006491 | Gm8369 | 0.006926 | Cbfa2t3 | 0.006439 |  |  |  |  |
| Ctsw | 0.005727 | Raph1 | 0.006083 | Gimap1 | 0.006744 | Cd74 | 0.006412 |  |  |  |  |
| Atp1b1 | 0.005571 | Tnfrsf18 | 0.005996 | AW112010 | 0.006584 | Olfm1 | 0.00585 |  |  |  |  |
| Arap3 | 0.00552 | Areg | 0.005975 | Gimap5 | 0.006501 | Ifi30 | 0.005709 |  |  |  |  |
| Pitpnm2 | 0.005506 | Icos | 0.005684 | Satb1 | 0.006102 | Naga | 0.005386 |  |  |  |  |
| Usp24 | 0.005466 | Gzmb | 0.005599 | Gm26917 | 0.005871 | Cd86 | 0.005333 |  |  |  |  |
| Fermt2 | 0.005405 | Tnfrsf9 | 0.005535 | Zbp1 | 0.005854 | Plbd1 | 0.005287 |  |  |  |  |
| Col27a1 | 0.005229 | Gpr55 | 0.005526 | Ly6a | 0.005706 | Cd209a | 0.00513 |  |  |  |  |
| Nrgn | 0.0052 | Hopx | 0.005492 | Lef1 | 0.0057 | Pak1 | 0.005058 |  |  |  |  |
| Trat1 | 0.005177 | Cd27 | 0.005418 | Trac | 0.005451 | Syngr2 | 0.004901 |  |  |  |  |
| Cntn1 | 0.005172 | Odc1 | 0.00541 | Slfn2 | 0.005353 | Prcp | 0.004863 |  |  |  |  |
| Spry2 | 0.005006 | Neb | 0.005348 | Gbp4 | 0.00527 | Rnase6 | 0.004808 |  |  |  |  |
| Zfp318 | 0.005006 | Klrg1 | 0.00525 | Slfn1 | 0.005268 | Atf3 | 0.00478 |  |  |  |  |

| Neuts factor-1 genes | Neuts factor-1 weight | Neuts factor-2 genes | Neuts factor-2 weight | Neuts factor-3 genes | Neuts factor-3 weight | Neuts factor-4 genes | Neuts factor-4 weight | Neuts factor-5 genes | Neuts factor-5 weight | Neuts factor-6 genes | Neuts factor-6 weight |
| --- | --- | --- | --- | --- | --- | --- | --- | --- | --- | --- | --- |
| Gngt2 | 0.005827 | Dgat1 | 0.008943 | Rsad2 | 0.010744 | Fam162a | 0.008727 | Fnip2 | 0.007267 | Ahnak | 0.00599 |
| Clec4n | 0.005412 | Rdh12 | 0.008593 | Ifi204 | 0.009683 | Ero1l | 0.008704 | Slc31a2 | 0.006697 | Lgmn | 0.00597 |
| Cst3 | 0.004436 | G0s2 | 0.008248 | Oasl2 | 0.009368 | Bnip3 | 0.008255 | Atp6v1a | 0.006569 | Apoe | 0.005741 |
| Gadd45b | 0.004432 | Retnlg | 0.007928 | Parp14 | 0.009121 | Bsg | 0.00758 | Klhdc4 | 0.006166 | Gm26917 | 0.00572 |
| Mrpl52 | 0.004371 | Wfdc21 | 0.007694 | Isg15 | 0.008892 | Higd1a | 0.007407 | Npc1 | 0.006127 | Lrp1 | 0.005351 |
| Nr4a1 | 0.00436 | Glrx | 0.007496 | Slfn5 | 0.00876 | Gpi1 | 0.007039 | Amdhd2 | 0.00598 | Fn1 | 0.005264 |
| Snrpg | 0.004318 | Wfdc17 | 0.007408 | Irf7 | 0.008747 | Impa2 | 0.006995 | Syne1 | 0.005893 | C1qb | 0.005215 |
| Dynll1 | 0.004249 | Ccl6 | 0.007239 | Ifit1 | 0.008436 | Mif | 0.006972 | 1700017B05Rik | 0.005735 | Mafb | 0.005188 |
| Lst1 | 0.004059 | Isy1 | 0.007111 | Oasl1 | 0.008148 | Egln3 | 0.006702 | Osbpl8 | 0.005558 | Maf | 0.004958 |
| Il1r2 | 0.003963 | Plaur | 0.006736 | Stat1 | 0.008093 | Npepps | 0.006208 | Plgrkt | 0.00552 | Selenop | 0.004914 |
| Fgl2 | 0.003873 | Thbs1 | 0.006595 | Slfn8 | 0.008062 | Tpi1 | 0.006195 | Ccdc126 | 0.005506 | Crip1 | 0.00488 |
| Dennd4a | 0.003858 | Ifitm1 | 0.006571 | Ddx60 | 0.008044 | Ndufv3 | 0.006162 | Eea1 | 0.005239 | Pid1 | 0.004784 |
| Wnk1 | 0.003834 | Tgm2 | 0.006273 | Plac8 | 0.007927 | Sec61g | 0.006134 | Plbd1 | 0.005217 | Xist | 0.004765 |
| Gng10 | 0.003778 | Mrgpra2b | 0.006178 | Rtp4 | 0.007906 | Bnip3l | 0.005857 | Mreg | 0.005165 | Lgals1 | 0.004717 |
| Ccrl2 | 0.003776 | Fam107b | 0.006101 | Isg20 | 0.007834 | Manf | 0.00584 | Naa50 | 0.00515 | Stab1 | 0.004702 |
| Cdk2ap2 | 0.003759 | Ly6g | 0.005701 | Ddx58 | 0.007829 | Hk2 | 0.005747 | Ctsz | 0.005146 | Emp1 | 0.004573 |
| Gm2a | 0.003712 | Cxcl3 | 0.005463 | Gbp2 | 0.007775 | Egln1 | 0.005744 | Igf2r | 0.005112 | Mrc1 | 0.004521 |
| Nfkbid | 0.003696 | Osm | 0.005439 | Usp18 | 0.007753 | Pgp | 0.005741 | Gla | 0.005074 | Pf4 | 0.004482 |
| Pcna | 0.00368 | Il1r2 | 0.005426 | Rnf213 | 0.007396 | Fndc3a | 0.005736 | Cstb | 0.005066 | Aif1 | 0.004415 |
| Tnfaip2 | 0.003602 | Lcn2 | 0.005361 | 9930111J21Rik2 | 0.007291 | P4hb | 0.005703 | Lamp1 | 0.005009 | C1qa | 0.004363 |
| Neuts factor-7 genes | Neuts factor-7 weight | Neuts factor-8 genes | Neuts factor-8 weight | Neuts factor-9 genes | Neuts factor-9 weight | | |  |  |  |  |
| Ccpg1 | 0.007117 | Id1 | 0.006872 | Arf2 | 0.007716 |  |  |  |  |  |  |
| Hp | 0.005989 | Slc38a2 | 0.006543 | Metrnl | 0.007077 |  |  |  |  |  |  |
| Limd2 | 0.005903 | Fam107b | 0.005532 | Emp3 | 0.006688 |  |  |  |  |  |  |
| Tspan13 | 0.005881 | Spp1 | 0.00545 | Adgre5 | 0.006675 |  |  |  |  |  |  |
| Dck | 0.005542 | Ddit4 | 0.005208 | Tnf | 0.005954 |  |  |  |  |  |  |
| Anxa1 | 0.005477 | Ccl4 | 0.005017 | Ceacam1 | 0.005635 |  |  |  |  |  |  |
| Csf3r | 0.005317 | Brd2 | 0.004963 | Capg | 0.005529 |  |  |  |  |  |  |
| Nfam1 | 0.005204 | Cdkn1a | 0.004682 | Gpx1 | 0.005376 |  |  |  |  |  |  |
| Pglyrp1 | 0.005154 | Nfkb1 | 0.004606 | Ezr | 0.005235 |  |  |  |  |  |  |
| Tmem71 | 0.005139 | Tnfrsf12a | 0.004586 | Stk10 | 0.005229 |  |  |  |  |  |  |
| Akap13 | 0.005076 | Ifrd1 | 0.004513 | Ckap4 | 0.005217 |  |  |  |  |  |  |
| Fyb | 0.004967 | Tgfb1 | 0.004501 | Rhof | 0.00508 |  |  |  |  |  |  |
| Prr5l | 0.004917 | Gadd45g | 0.004401 | C3 | 0.005044 |  |  |  |  |  |  |
| Cytip | 0.004902 | Rgcc | 0.004314 | Rnh1 | 0.004968 |  |  |  |  |  |  |
| Hist1h2bc | 0.004896 | Rybp | 0.004283 | Lair1 | 0.004797 |  |  |  |  |  |  |
| Fos | 0.004855 | Fam71f2 | 0.004272 | Zbtb7a | 0.004615 |  |  |  |  |  |  |
| Mtus1 | 0.004805 | Rilpl2 | 0.004245 | Ncoa3 | 0.004592 |  |  |  |  |  |  |
| Rflnb | 0.004706 | Plaur | 0.004237 | AA467197 | 0.004557 |  |  |  |  |  |  |
| Rassf3 | 0.004681 | Csf3 | 0.004232 | Sema4d | 0.004552 |  |  |  |  |  |  |
| Nlrp12 | 0.004667 | Ncoa4 | 0.004192 | Icam1 | 0.004525 |  |  |  |  |  |  |
| DSP factor-1 genes | DSP factor-1 weight | DSP factor-2 genes | DSP factor-2 weight | DSP factor-3 genes | DSP factor-3 weight | DSP factor-4 genes | DSP factor-4 weight | DSP factor-5 genes | DSP factor-5 weight | DSP factor-6 genes | DSP factor-6 weight |
| Cxcl2 | 0.005036 | Zfhx4 | 0.004852 | Ogt | 0.004435 | Igfbp3 | 0.004402 | Sox18 | 0.00509 | Tmem100 | 0.003311 |
| S100a8 | 0.004507 | Meg3 | 0.004633 | Mir155hg | 0.004415 | Ogn | 0.004291 | Cd24a | 0.004649 | Stxbp6 | 0.003281 |
| Srgn | 0.004384 | Fam171b | 0.004424 | Grk3 | 0.004287 | Eln | 0.004098 | Ptch1 | 0.004226 | Galk2 | 0.003251 |
| Lyz2 | 0.004304 | Ptprd | 0.004353 | Gm20342 | 0.004122 | Myl9 | 0.004066 | Ctbp2 | 0.004149 | Cxcl14 | 0.003232 |
| Dusp1 | 0.004283 | Pappa2 | 0.00424 | Pnisr | 0.003932 | Tspan11 | 0.003931 | Tmem158 | 0.004096 | Fgf10 | 0.00319 |
| S100a9 | 0.004262 | Spock3 | 0.004102 | Ints6l | 0.003741 | Tagln | 0.003885 | Bmp7 | 0.004066 | Gm14226 | 0.003165 |
| Nfkbia | 0.00416 | Zcchc18 | 0.004043 | Kcnq1ot1 | 0.003698 | Mfap4 | 0.00377 | Sms | 0.004065 | Plagl1 | 0.003066 |
| Cd74 | 0.003975 | Sparcl1 | 0.003798 | Nfat5 | 0.003683 | Col12a1 | 0.00361 | Adamts15 | 0.004017 | Col14a1 | 0.003045 |
| Ccl4 | 0.003956 | Luzp2 | 0.003785 | Gm26917 | 0.003681 | Pmepa1 | 0.003552 | Notch1 | 0.003949 | Crabp2 | 0.003043 |
| Klf6 | 0.003903 | Sox2 | 0.003661 | Nav2 | 0.003642 | Smoc2 | 0.003545 | Sox11 | 0.003941 | Maged2 | 0.002945 |
| Pim1 | 0.003848 | Sfrp2 | 0.003528 | Nktr | 0.003641 | Klf4 | 0.003505 | Sdc3 | 0.003891 | Alpl | 0.00291 |
| Dcn | 0.003806 | Gm26771 | 0.003504 | Ralgapa2 | 0.003615 | Col7a1 | 0.003482 | Stmn1 | 0.003833 | Socs3 | 0.002877 |
| Il1b | 0.0038 | Kcnq3 | 0.003452 | AC149090.1 | 0.00361 | Slc6a6 | 0.003475 | Filip1l | 0.003808 | Prss12 | 0.002874 |
| Lgmn | 0.003689 | Kctd12 | 0.003426 | Zfp950 | 0.003595 | Crispld2 | 0.00347 | Lrrn1 | 0.003747 | Id3 | 0.002779 |
| Lgals3 | 0.003679 | Bambi | 0.003402 | Psd3 | 0.003594 | Dpep1 | 0.003422 | Ncald | 0.003743 | Ecrg4 | 0.002777 |
| Crip1 | 0.003665 | Unc5c | 0.003352 | Abca8a | 0.003522 | Foxd2os | 0.003394 | Jpt1 | 0.003736 | Gpx3 | 0.002755 |
| Fn1 | 0.003566 | Phf20l1 | 0.00335 | Smg1 | 0.003501 | Emp1 | 0.003379 | Prdm1 | 0.003589 | Sod3 | 0.002751 |
| Col1a1 | 0.003528 | Fst | 0.00332 | 4632427E13Rik | 0.00349 | Tpm2 | 0.003303 | S100a10 | 0.003537 | S100a1 | 0.002747 |
| Tyrobp | 0.003502 | S100b | 0.00331 | Nav3 | 0.003443 | Rasgrp2 | 0.003247 | Stat5a | 0.003528 | Map1b | 0.002743 |
| Junb | 0.003471 | Trim9 | 0.003253 | Clk1 | 0.003394 | Cnn2 | 0.003247 | Rgs12 | 0.003509 | Gstm1 | 0.002727 |

| DC factor-1 genes | DC factor-1 weight | DC factor-2 genes | DC factor-2 weight | DC factor-3 genes | DC factor-3 weight | DC factor-4 genes | DC factor-4 weight | DC factor-5 genes | DC factor-5 weight | DC factor-6 genes | DC factor-6 weight |
| --- | --- | --- | --- | --- | --- | --- | --- | --- | --- | --- | --- |
| Ass1 | 0.004575 | Clec4d | 0.003757 | Icosl | 0.00517 | Col1a2 | 0.005048 | Epsti1 | 0.005357 | Selenop | 0.005245 |
| Ramp3 | 0.004474 | Cd14 | 0.003461 | Prnp | 0.005107 | Col3a1 | 0.004547 | Net1 | 0.005207 | Apoe | 0.004938 |
| Tmem131 | 0.004382 | Arg1 | 0.003458 | Nostrin | 0.00505 | Dmxl1 | 0.004381 | Arhgap31 | 0.004867 | C1qb | 0.004852 |
| Ccnd2 | 0.004314 | Thbs1 | 0.00345 | Snrnp25 | 0.004904 | Tiam1 | 0.004288 | Pakap.1 | 0.004681 | Lyz2 | 0.00468 |
| Pvr | 0.004274 | Cxcl3 | 0.003365 | Tnfrsf18 | 0.004823 | Col1a1 | 0.004274 | Zmynd15 | 0.004653 | C1qa | 0.004671 |
| Helz2 | 0.004228 | Ccl4 | 0.003354 | Ptpn1 | 0.00468 | Cxcr4 | 0.004146 | Txndc17 | 0.004644 | Wwp1 | 0.004302 |
| Klrk1 | 0.003983 | Ets2 | 0.003335 | Chd3 | 0.004677 | Prkab1 | 0.003926 | Atxn1 | 0.004474 | Mrc1 | 0.004297 |
| Got1 | 0.003865 | Clec4e | 0.003214 | Cdc42ep3 | 0.004659 | Picalm | 0.003768 | AW112010 | 0.00443 | C1qc | 0.004221 |
| Nfkb1 | 0.003785 | Ccr1 | 0.003209 | Pold1 | 0.004656 | Lgals7 | 0.003767 | Apol7c | 0.004378 | Pltp | 0.004166 |
| Il2ra | 0.003755 | Bnip3 | 0.00318 | Zfand6 | 0.004484 | Lman2l | 0.003755 | Plaat3 | 0.004321 | Ehd4 | 0.004165 |
| AA467197 | 0.003693 | C5ar1 | 0.00317 | Fam107b | 0.004479 | Slc8a1 | 0.003735 | AY036118 | 0.004277 | Timp2 | 0.003993 |
| Acadm | 0.003635 | Msr1 | 0.00311 | Ly86 | 0.004396 | Skil | 0.003724 | Frmd4a | 0.0042 | Ctsd | 0.003916 |
| Nr4a3 | 0.003618 | Ero1l | 0.00308 | Gsto1 | 0.004286 | Kdm6b | 0.003667 | H2-M2 | 0.004198 | Lgmn | 0.003754 |
| Bhlhe40 | 0.003586 | Card19 | 0.003073 | Tcf7 | 0.004187 | Usp38 | 0.003663 | Nudt17 | 0.00416 | Nrp1 | 0.003741 |
| Cd83 | 0.003547 | Emilin2 | 0.003071 | Hmgn3 | 0.004157 | Sparc | 0.003569 | Zc3h12c | 0.004095 | Ctsb | 0.003727 |
| Foxp1 | 0.003538 | Cebpb | 0.003062 | Ccnd1 | 0.004148 | Nfkbid | 0.003563 | Mthfd2 | 0.004093 | Pld4 | 0.003704 |
| Ddt | 0.003481 | Snx18 | 0.003041 | Crispld2 | 0.004139 | Xpo4 | 0.003537 | Rabgap1l | 0.004073 | Unc93b1 | 0.003657 |
| Isg15 | 0.003431 | Cxcl2 | 0.003039 | Serpina3g | 0.00411 | S100a9 | 0.003534 | Rnf115 | 0.004049 | Grn | 0.003639 |
| Cd86 | 0.003426 | Hmox1 | 0.003025 | Gyg | 0.004082 | Adgre5 | 0.003524 | Lima1 | 0.004021 | Ptpn18 | 0.003589 |
| Grasp | 0.00339 | Vegfa | 0.002974 | Klrd1 | 0.004048 | Fgr | 0.003468 | Ccl5 | 0.004021 | Cyth4 | 0.003578 |

| Melano factor-1 genes | Melano factor-1 weight | Melano factor-2 genes | Melano factor-2 weight | Melano factor-3 genes | Melano factor-3 weight | Melano factor-4 genes | Melano factor-4 weight | Melano factor-5 genes | Melano factor-5 weight |
| --- | --- | --- | --- | --- | --- | --- | --- | --- | --- |
| Stmn1 | 0.005161 | Col1a1 | 0.005441 | Gpr143 | 0.004033 | Cdc42ep5 | 0.004647 | Enpp2 | 0.004079 |
| Smc2 | 0.004437 | Col1a2 | 0.005054 | Rab38 | 0.003909 | Prpf4 | 0.004596 | Col12a1 | 0.004044 |
| Hmgb2 | 0.004339 | S100a6 | 0.004878 | Mc1r | 0.003871 | Fbln5 | 0.004545 | Zbtb20 | 0.003991 |
| Lockd | 0.004141 | Col3a1 | 0.004608 | Slc24a5 | 0.003788 | Fopnl | 0.004542 | Txnip | 0.003901 |
| Rrm1 | 0.004073 | Cxcl2 | 0.004417 | Igsf8 | 0.003746 | Pmel | 0.004509 | Cdc14a | 0.003814 |
| Cks1b | 0.00403 | Ifitm3 | 0.004218 | Nudt16l1 | 0.003655 | Sub1 | 0.004501 | Bcl2 | 0.003799 |
| Lmnb1 | 0.003917 | Lyz2 | 0.004159 | Trpm1 | 0.003653 | Mgll | 0.004403 | Auts2 | 0.003793 |
| Cenpw | 0.003854 | Dusp1 | 0.00412 | Slc3a2 | 0.003642 | Cebpg | 0.004365 | Mylk | 0.003783 |
| Dut | 0.003838 | Nfkbia | 0.004079 | Nckap5l | 0.003626 | Sik1 | 0.004276 | Plekhm2 | 0.003755 |
| Rfc4 | 0.003755 | Tmsb4x | 0.003981 | Mlana | 0.003592 | Tmem64 | 0.004273 | Rsrp1 | 0.003554 |
| Mki67 | 0.003744 | Pf4 | 0.003962 | Aph1c | 0.003514 | Dmwd | 0.00415 | Adgrg1 | 0.003537 |
| Cenpe | 0.003704 | Ifi27l2a | 0.00378 | Gng11 | 0.003502 | Flot2 | 0.004149 | Rnase1 | 0.003492 |
| H2afx | 0.003647 | Crip1 | 0.003779 | Emc8 | 0.003404 | Adam10 | 0.004124 | Sptbn1 | 0.003479 |
| Tyms | 0.003632 | S100a9 | 0.003769 | Sec11c | 0.003389 | Cacul1 | 0.004121 | C4b | 0.003424 |
| Cenpa | 0.003599 | Cd74 | 0.003743 | Mlph | 0.003324 | Gnaq | 0.004051 | Pik3ip1 | 0.003418 |
| Birc5 | 0.003593 | Sparc | 0.003725 | Ctsz | 0.003298 | Znrf2 | 0.004027 | Appl2 | 0.003389 |
| Knstrn | 0.003559 | Pim1 | 0.003657 | Slc45a2 | 0.00328 | Tyr | 0.003989 | Sema5a | 0.00338 |
| Atad2 | 0.003483 | Cebpb | 0.003635 | Tspan10 | 0.003274 | Tyrp1 | 0.003974 | Cited1 | 0.003351 |
| Ube2s | 0.003452 | Lgmn | 0.003524 | Specc1 | 0.00319 | Socs4 | 0.003974 | Abca8a | 0.003349 |
| Spc24 | 0.003449 | Zfp36 | 0.00348 | Atp6v0a1 | 0.003182 | Bet1 | 0.003938 | Nav1 | 0.00334 |

| VSM factor-1 genes | VSM factor-1 weight | VSM factor-2 genes | VSM factor-2 weight | VSM factor-3 genes | VSM factor-3 weight |
| --- | --- | --- | --- | --- | --- |
| Myh11 | 0.003089 | Slfn5 | 0.002566 | Tubb4b | 0.003382 |
| Lmod1 | 0.002917 | Sept4 | 0.002481 | Ckap4 | 0.003148 |
| Cbx6 | 0.002909 | B2m | 0.002381 | Hmgb2 | 0.003147 |
| Ppp1r12b | 0.002909 | Tmem176b | 0.002378 | Hmgn2 | 0.003101 |
| Sncg | 0.002865 | Tmem176a | 0.002375 | Tubb6 | 0.00308 |
| Mustn1 | 0.002848 | Cyp4b1 | 0.002334 | Selenoh | 0.003076 |
| Bcam | 0.002842 | Clec2d | 0.002328 | Ube2s | 0.002961 |
| Dstn | 0.002784 | Adamts12 | 0.002317 | H2afz | 0.00295 |
| Igfbp5 | 0.002767 | Vstm4 | 0.002276 | AI506816 | 0.002902 |
| Pcp4l1 | 0.002683 | Meg3 | 0.002252 | Ran | 0.002858 |
| Wtip | 0.002654 | Ifitm1 | 0.002248 | Tmpo | 0.002849 |
| Sorbs2 | 0.002631 | Abcc9 | 0.002242 | Hjurp | 0.002839 |
| Syne1 | 0.002626 | Higd1b | 0.002242 | Dnmt1 | 0.002822 |
| Plpp3 | 0.002619 | Vtn | 0.002231 | Stmn1 | 0.002815 |
| Mob2 | 0.002605 | H2-D1 | 0.002223 | Tubb5 | 0.002781 |
| Crispld2 | 0.002591 | Serpine2 | 0.002204 | Tuba1b | 0.002742 |
| Lama5 | 0.002586 | Serping1 | 0.002196 | Tyms | 0.002696 |
| Fxyd1 | 0.002586 | Prrx1 | 0.002187 | Kdelr2 | 0.002688 |
| Rassf3 | 0.002573 | Tnfrsf21 | 0.002176 | Dut | 0.002682 |
| Emilin1 | 0.002552 | Ebf1 | 0.002156 | Nasp | 0.002639 |

| TNK factor-1 genes | TNK factor-1 weight | TNK factor-2 genes | TNK factor-2 weight | TNK factor-3 genes | TNK factor-3 weight |
| --- | --- | --- | --- | --- | --- |
| Grn | 0.003514 | Cxcl2 | 0.005512 | Klre1 | 0.004226 |
| Ctss | 0.003275 | S100a9 | 0.005193 | Bcl2 | 0.004027 |
| Maf | 0.003247 | S100a8 | 0.004912 | Ncr1 | 0.004018 |
| Lgmn | 0.003245 | S100a11 | 0.004879 | Nkg7 | 0.003902 |
| Ctsl | 0.003143 | Il1b | 0.00487 | Txk | 0.003897 |
| Snx2 | 0.003117 | Mcl1 | 0.004818 | Ms4a4b | 0.00381 |
| Unc93b1 | 0.003108 | Nfkbia | 0.004691 | Macf1 | 0.003778 |
| Ctsh | 0.003087 | Srgn | 0.004679 | Klrb1c | 0.003742 |
| Sirpa | 0.003081 | Cebpb | 0.004614 | Ctla2a | 0.003653 |
| Ctsb | 0.003031 | Ccl3 | 0.004596 | Klrd1 | 0.003644 |
| Gpx1 | 0.002988 | Clec4e | 0.004534 | Hist1h1e | 0.003591 |
| Aif1 | 0.002953 | Egr1 | 0.004468 | Xcl1 | 0.003576 |
| Cd68 | 0.002923 | Cd9 | 0.004418 | Arsb | 0.003551 |
| Csf1r | 0.002917 | Cxcl3 | 0.004292 | H2afz | 0.00351 |
| Lgals3 | 0.002915 | Ptgs2 | 0.004243 | Trbc1 | 0.003459 |
| Ctsc | 0.002908 | Clec4d | 0.004242 | AW112010 | 0.003454 |
| Lrp1 | 0.002886 | Cd14 | 0.004176 | Stat4 | 0.003405 |
| Dab2 | 0.002877 | Dusp1 | 0.004127 | Xist | 0.003382 |
| Ms4a6c | 0.002859 | Grina | 0.004113 | Prf1 | 0.003263 |
| Psap | 0.002852 | Fth1 | 0.004013 | Klrb1f | 0.003262 |

| Mast factor-1 genes | Mast factor-1 weight | Mast factor-2 genes | Mast factor-2 weight |
| --- | --- | --- | --- |
| Ly6e | 0.002562 | Cpa3 | 0.003731 |
| Lyz2 | 0.002508 | Cma1 | 0.003618 |
| Nfkbia | 0.002431 | Serpinb1a | 0.003394 |
| Cd52 | 0.002388 | Tpsb2 | 0.003344 |
| Ccl4 | 0.00237 | Kit | 0.003151 |
| Btg1 | 0.002282 | Hdc | 0.003106 |
| Ctsb | 0.002278 | Ndrg1 | 0.003063 |
| Alox5ap | 0.002269 | Hsp90aa1 | 0.003042 |
| Cd74 | 0.002242 | Dusp1 | 0.002969 |
| Coro1a | 0.002212 | Ubb | 0.002915 |
| Marcks | 0.00219 | Ly6a | 0.002899 |
| Ptpn18 | 0.00218 | Cited2 | 0.002881 |
| Ifitm3 | 0.002167 | Fos | 0.002877 |
| Actr3 | 0.00215 | Ccl2 | 0.00283 |
| Pim1 | 0.002147 | Crip1 | 0.002823 |
| Ctss | 0.002143 | Apoe | 0.002812 |
| Cebpb | 0.002126 | Jun | 0.002798 |
| Myh9 | 0.002121 | Slc6a4 | 0.002767 |
| Lgmn | 0.002103 | Hs6st2 | 0.002751 |
| Ifi27l2a | 0.002099 | Sec11c | 0.00273 |

| B factor-1 genes | B factor-1 weight | B factor-2 genes | B factor-2 weight |
| --- | --- | --- | --- |
| Akr1a1 | 0.002186 | Ighm | 0.003043 |
| Lamp1 | 0.002159 | Igkc | 0.002968 |
| Fcer1g | 0.002132 | H2-Ab1 | 0.002903 |
| Sdcbp | 0.002129 | H2-Aa | 0.00286 |
| Tyrobp | 0.002126 | Ly6d | 0.002859 |
| Cstb | 0.002087 | H2-Eb1 | 0.00281 |
| Anxa5 | 0.00207 | Iglc3 | 0.002767 |
| Fxyd5 | 0.002002 | Iglc2 | 0.002747 |
| Cebpb | 0.001998 | Ms4a1 | 0.002612 |
| Lgmn | 0.001991 | Gm42418 | 0.002591 |
| Lyz2 | 0.001985 | Xist | 0.002581 |
| Ctsb | 0.001978 | Ltb | 0.002536 |
| Grn | 0.001977 | Macf1 | 0.002504 |
| Fyb | 0.00197 | Ighd | 0.002433 |
| Anxa2 | 0.00195 | Hist1h1e | 0.002392 |
| Psap | 0.001948 | Dmxl1 | 0.002359 |
| Apoe | 0.001947 | Dennd4a | 0.002344 |
| Lilrb4a | 0.001943 | Fcmr | 0.002308 |
| Ifitm3 | 0.001938 | Mzb1 | 0.002289 |
| Litaf | 0.001928 | Klf2 | 0.002275 |
