## Supplemental Table 2 for "Space-Time Mapping Identifies Concerted Multicellular Patterns and Gene Programs in Healing Wounds and their Conservation in Cancers"

**Supplementary Table 2**

**Tumor Factor 6 and WH Factor 22**

| **Term** | **Overlap** | **P-value** | **Adjusted P-value** | **Old P-value** | **Old Adjusted P-value** | **Odds Ratio** | **Combined Score** | **Genes** |
| --- | --- | --- | --- | --- | --- | --- | --- | --- |
| neutrophil degranulation (GO:0043312) | 6/481 | 2.79E-07 | 3.45E-05 | 0 | 0 | 35.20962 | 531.4005 | VAT1;CD63;SYNGR1;FABP5;TIMP2;CTSD |
| neutrophil activation involved in immune response (GO:0002283) | 6/485 | 2.93E-07 | 3.45E-05 | 0 | 0 | 34.90844 | 525.1539 | VAT1;CD63;SYNGR1;FABP5;TIMP2;CTSD |
| neutrophil mediated immunity (GO:0002446) | 6/488 | 3.04E-07 | 3.45E-05 | 0 | 0 | 34.68583 | 520.5468 | VAT1;CD63;SYNGR1;FABP5;TIMP2;CTSD |
| regulation of sequestering of triglyceride (GO:0010889) | 2/15 | 4.08E-05 | 0.002646 | 0 | 0 | 279.3566 | 2823.716 | LPL;TREM2 |
| regulation of establishment of protein localization (GO:0070201) | 2/15 | 4.08E-05 | 0.002646 | 0 | 0 | 279.3566 | 2823.716 | TREM2;CTSD |
| regulation of cholesterol storage (GO:0010885) | 2/16 | 4.66E-05 | 0.002646 | 0 | 0 | 259.3896 | 2587.349 | LPL;TREM2 |
| triglyceride catabolic process (GO:0019433) | 2/23 | 9.79E-05 | 0.00477 | 0 | 0 | 172.8658 | 1595.797 | FABP5;LPL |
| regulation of cytokine production (GO:0001817) | 3/150 | 0.000112 | 0.00477 | 0 | 0 | 40.4898 | 368.3669 | GPNMB;TREM2;PLD3 |
| acylglycerol catabolic process (GO:0046464) | 2/35 | 0.000229 | 0.008686 | 0 | 0 | 109.9394 | 921.3629 | FABP5;LPL |
| regulation of chemokine production (GO:0032642) | 2/42 | 0.000331 | 0.010755 | 0 | 0 | 90.66818 | 726.5853 | LPL;TREM2 |
| regulation of cytokine production involved in inflammatory response (GO:1900015) | 2/43 | 0.000347 | 0.010755 | 0 | 0 | 88.45233 | 704.6477 | TREM2;PLD3 |
| triglyceride metabolic process (GO:0006641) | 2/55 | 0.000568 | 0.016141 | 0 | 0 | 68.38422 | 511.0599 | FABP5;LPL |
| positive regulation of MAPK cascade (GO:0043410) | 3/274 | 0.000657 | 0.017234 | 0 | 0 | 21.82583 | 159.9355 | GPNMB;GDF15;TREM2 |
| negative regulation of cellular catabolic process (GO:0031330) | 2/69 | 0.000893 | 0.021745 | 0 | 0 | 54.05699 | 379.5445 | TIMP2;TREM2 |
| positive regulation of protein phosphorylation (GO:0001934) | 3/371 | 0.001577 | 0.029872 | 0 | 0 | 15.99375 | 103.1958 | GPNMB;GDF15;TREM2 |
| regulation of interleukin-6 production (GO:0032675) | 2/110 | 0.002247 | 0.029872 | 0 | 0 | 33.46633 | 204.0777 | LPL;TREM2 |
| regulation of growth hormone receptor signaling pathway (GO:0060398) | 1/5 | 0.003246 | 0.029872 | 0 | 0 | 416.3125 | 2385.602 | GDF15 |
| cellular response to nutrient (GO:0031670) | 1/5 | 0.003246 | 0.029872 | 0 | 0 | 416.3125 | 2385.602 | LPL |
| regulation of hippocampal neuron apoptotic process (GO:0110089) | 1/5 | 0.003246 | 0.029872 | 0 | 0 | 416.3125 | 2385.602 | TREM2 |

**Tumor Factor 10 and WH Factor 8**

| **Term** | **Overlap** | **P-value** | **Adjusted P-value** | **Old P-value** | **Old Adjusted P-value** | **Odds Ratio** | **Combined Score** | **Genes** |
| --- | --- | --- | --- | --- | --- | --- | --- | --- |
| cellular response to hypoxia (GO:0071456) | 5/131 | 2.74E-09 | 2.9E-07 | 0 | 0 | 157.6508 | 3108.125 | EGLN3;FAM162A;BNIP3;MGARP;HIGD1A |
| cellular response to decreased oxygen levels (GO:0036294) | 4/69 | 2.68E-08 | 1.42E-06 | 0 | 0 | 204.359 | 3562.865 | EGLN3;FAM162A;BNIP3;MGARP |
| positive regulation of mitochondrion organization (GO:0010822) | 3/58 | 2.74E-06 | 9.67E-05 | 0 | 0 | 155.3377 | 1989.643 | FAM162A;BNIP3;MGARP |
| neuron apoptotic process (GO:0051402) | 2/21 | 4.7E-05 | 0.001246 | 0 | 0 | 262.7763 | 2618.598 | FAM162A;BNIP3 |
| positive regulation of release of cytochrome c from mitochondria (GO:0090200) | 2/25 | 6.71E-05 | 0.001334 | 0 | 0 | 217.0326 | 2085.578 | FAM162A;BNIP3 |
| mitochondrion organization (GO:0007005) | 3/175 | 7.55E-05 | 0.001334 | 0 | 0 | 49.3804 | 468.6684 | BNIP3;MGARP;HIGD1A |
| apoptotic process (GO:0006915) | 3/231 | 0.000172 | 0.002419 | 0 | 0 | 37.14662 | 322.0113 | EGLN3;FAM162A;BNIP3 |
| regulation of release of cytochrome c from mitochondria (GO:0090199) | 2/41 | 0.000183 | 0.002419 | 0 | 0 | 127.891 | 1100.918 | FAM162A;BNIP3 |
| regulation of apoptotic process (GO:0042981) | 4/742 | 0.00033 | 0.00362 | 0 | 0 | 17.39115 | 139.4168 | EGLN3;FAM162A;BNIP3;HIGD1A |
| mitochondrial transport (GO:0006839) | 2/56 | 0.000342 | 0.00362 | 0 | 0 | 92.2963 | 736.7103 | BNIP3;MGARP |
| regulation of transcription from RNA polymerase II promoter in response to hypoxia (GO:0061418) | 2/75 | 0.000612 | 0.005901 | 0 | 0 | 68.2089 | 504.6238 | EGLN3;HIGD1A |
| activation of cysteine-type endopeptidase activity involved in apoptotic process (GO:0006919) | 2/81 | 0.000714 | 0.006305 | 0 | 0 | 63.00949 | 456.4967 | EGLN3;FAM162A |
| regulation of transcription from RNA polymerase II promoter in response to stress (GO:0043618) | 2/87 | 0.000823 | 0.00671 | 0 | 0 | 58.54412 | 415.8217 | EGLN3;HIGD1A |
| positive regulation of cysteine-type endopeptidase activity involved in apoptotic process (GO:0043280) | 2/119 | 0.001531 | 0.011594 | 0 | 0 | 42.46368 | 275.2364 | EGLN3;FAM162A |
| disaccharide biosynthetic process (GO:0046351) | 1/5 | 0.002498 | 0.016547 | 0 | 0 | 555.1667 | 3326.771 | SLC2A1 |
| glucose import across plasma membrane (GO:0098708) | 1/5 | 0.002498 | 0.016547 | 0 | 0 | 555.1667 | 3326.771 | SLC2A1 |
| mitochondrial protein catabolic process (GO:0035694) | 1/6 | 0.002997 | 0.017647 | 0 | 0 | 444.1111 | 2580.412 | BNIP3 |
| hexose import across plasma membrane (GO:0140271) | 1/6 | 0.002997 | 0.017647 | 0 | 0 | 444.1111 | 2580.412 | SLC2A1 |
| polyamine biosynthetic process (GO:0006596) | 1/7 | 0.003495 | 0.018313 | 0 | 0 | 370.0741 | 2093.271 | SMOX |

**Tumor Factor 11 and WH Factor 14**

| **Term** | **Overlap** | **P-value** | **Adjusted P-value** | **Old P-value** | **Old Adjusted P-value** | **Odds Ratio** | **Combined Score** | **Genes** |
| --- | --- | --- | --- | --- | --- | --- | --- | --- |
| negative regulation of cell activation (GO:0050866) | 2/14 | 2.49E-05 | 0.006558 | 0 | 0 | 369.9444 | 3921.13 | PDGFA;TREM2 |
| regulation of protein phosphorylation (GO:0001932) | 3/266 | 0.000355 | 0.026726 | 0 | 0 | 28.12643 | 223.4462 | TMEM119;PDGFA;TREM2 |
| negative regulation of protein-containing complex assembly (GO:0031333) | 2/52 | 0.000359 | 0.026726 | 0 | 0 | 88.61778 | 702.8769 | PMEPA1;TREM2 |
| positive regulation of phosphatidylinositol 3-kinase signaling (GO:0014068) | 2/77 | 0.000787 | 0.026726 | 0 | 0 | 59.00444 | 421.7392 | PDGFA;TREM2 |
| positive regulation of protein phosphorylation (GO:0001934) | 3/371 | 0.000935 | 0.026726 | 0 | 0 | 19.99423 | 139.4542 | TMEM119;PDGFA;TREM2 |
| regulation of peptidyl-tyrosine phosphorylation (GO:0050730) | 2/92 | 0.001121 | 0.026726 | 0 | 0 | 49.13333 | 333.8105 | PDGFA;TREM2 |
| regulation of phosphatidylinositol 3-kinase signaling (GO:0014066) | 2/106 | 0.001483 | 0.026726 | 0 | 0 | 42.48932 | 276.7508 | PDGFA;TREM2 |
| regulation of glomerular mesangial cell proliferation (GO:0072124) | 1/5 | 0.002747 | 0.026726 | 0 | 0 | 499.625 | 2946.374 | PDGFA |
| regulation of hippocampal neuron apoptotic process (GO:0110089) | 1/5 | 0.002747 | 0.026726 | 0 | 0 | 499.625 | 2946.374 | TREM2 |
| chemokine (C-X-C motif) ligand 12 signaling pathway (GO:0038146) | 1/5 | 0.002747 | 0.026726 | 0 | 0 | 499.625 | 2946.374 | TREM2 |
| regulation of inward rectifier potassium channel activity (GO:1901979) | 1/5 | 0.002747 | 0.026726 | 0 | 0 | 499.625 | 2946.374 | TREM2 |
| regulation of astrocyte activation (GO:0061888) | 1/5 | 0.002747 | 0.026726 | 0 | 0 | 499.625 | 2946.374 | TREM2 |
| mRNA cleavage (GO:0006379) | 1/5 | 0.002747 | 0.026726 | 0 | 0 | 499.625 | 2946.374 | RNASE4 |
| regulation of phosphatidylinositol biosynthetic process (GO:0010511) | 1/5 | 0.002747 | 0.026726 | 0 | 0 | 499.625 | 2946.374 | PDGFA |
| negative regulation of astrocyte differentiation (GO:0048712) | 1/6 | 0.003296 | 0.026726 | 0 | 0 | 399.68 | 2284.211 | TREM2 |
| negative regulation of sequestering of triglyceride (GO:0010891) | 1/6 | 0.003296 | 0.026726 | 0 | 0 | 399.68 | 2284.211 | TREM2 |
| regulation of resting membrane potential (GO:0060075) | 1/6 | 0.003296 | 0.026726 | 0 | 0 | 399.68 | 2284.211 | TREM2 |
| positive regulation of osteoblast proliferation (GO:0033690) | 1/6 | 0.003296 | 0.026726 | 0 | 0 | 399.68 | 2284.211 | TMEM119 |
| negative regulation of toll-like receptor 2 signaling pathway (GO:0034136) | 1/6 | 0.003296 | 0.026726 | 0 | 0 | 399.68 | 2284.211 | TREM2 |

**Tumor Factor 12 and WH Factor 1**

| **Term** | **Overlap** | **P-value** | **Adjusted P-value** | **Old P-value** | **Old Adjusted P-value** | **Odds Ratio** | **Combined Score** | **Genes** |
| --- | --- | --- | --- | --- | --- | --- | --- | --- |
| negative regulation of viral entry into host cell (GO:0046597) | 2/17 | 5.27E-05 | 0.003582 | 0 | 0 | 242.0848 | 2384.527 | IFITM3;IFITM2 |
| response to interferon-alpha (GO:0035455) | 2/18 | 5.93E-05 | 0.003582 | 0 | 0 | 226.9432 | 2208.735 | IFITM3;IFITM2 |
| negative regulation of viral life cycle (GO:1903901) | 2/20 | 7.36E-05 | 0.003582 | 0 | 0 | 201.7071 | 1919.584 | IFITM3;IFITM2 |
| response to interferon-beta (GO:0035456) | 2/28 | 0.000146 | 0.00533 | 0 | 0 | 139.5874 | 1232.801 | IFITM3;IFITM2 |
| regulation of viral entry into host cell (GO:0046596) | 2/39 | 0.000285 | 0.008325 | 0 | 0 | 98.0344 | 800.2224 | IFITM3;IFITM2 |
| negative regulation of viral genome replication (GO:0045071) | 2/54 | 0.000548 | 0.012193 | 0 | 0 | 69.7028 | 523.4705 | IFITM3;IFITM2 |
| type I interferon signaling pathway (GO:0060337) | 2/65 | 0.000793 | 0.012193 | 0 | 0 | 57.50072 | 410.5592 | IFITM3;IFITM2 |
| cellular response to type I interferon (GO:0071357) | 2/65 | 0.000793 | 0.012193 | 0 | 0 | 57.50072 | 410.5592 | IFITM3;IFITM2 |
| regulation of viral genome replication (GO:0045069) | 2/67 | 0.000842 | 0.012193 | 0 | 0 | 55.72587 | 394.524 | IFITM3;IFITM2 |
| innate immune response (GO:0045087) | 3/302 | 0.000871 | 0.012193 | 0 | 0 | 19.75385 | 139.1744 | IFITM3;IFITM2;MSRB1 |
| negative regulation of viral process (GO:0048525) | 2/70 | 0.000919 | 0.012193 | 0 | 0 | 53.25936 | 372.4199 | IFITM3;IFITM2 |
| response to interferon-gamma (GO:0034341) | 2/80 | 0.001198 | 0.014572 | 0 | 0 | 46.40793 | 312.2029 | IFITM3;IFITM2 |
| positive regulation of cold-induced thermogenesis (GO:0120162) | 2/97 | 0.001754 | 0.019696 | 0 | 0 | 38.07081 | 241.5976 | PLAC8;CCR2 |
| positive regulation of metabolic process (GO:0009893) | 2/113 | 0.00237 | 0.023689 | 0 | 0 | 32.55692 | 196.8079 | PLAC8;CCR2 |
| defense response to symbiont (GO:0140546) | 2/124 | 0.002844 | 0.023689 | 0 | 0 | 29.60507 | 173.5594 | IFITM3;IFITM2 |
| monocyte extravasation (GO:0035696) | 1/5 | 0.003246 | 0.023689 | 0 | 0 | 416.3125 | 2385.602 | CCR2 |
| negative regulation of hydrogen peroxide metabolic process (GO:0010727) | 1/5 | 0.003246 | 0.023689 | 0 | 0 | 416.3125 | 2385.602 | HP |
| defense response to virus (GO:0051607) | 2/133 | 0.003263 | 0.023689 | 0 | 0 | 27.55864 | 157.776 | IFITM3;IFITM2 |
| neutrophil degranulation (GO:0043312) | 3/481 | 0.003303 | 0.023689 | 0 | 0 | 12.24414 | 69.94967 | PLAC8;HP;S100A11 |

**Tumor Factor 13 WH Factor 4**

| **GO biological process complete** | **Mus musculus - REFLIST (21997)** | **Client Text Box Input (11)** | **Client Text Box Input (expected)** | **Client Text Box Input (over/under)** | **Client Text Box Input (fold Enrichment)** | **Client Text Box Input (raw P-value)** | **Client Text Box Input (FDR)** |
| --- | --- | --- | --- | --- | --- | --- | --- |
| peptide antigen assembly with MHC class II protein complex (GO:0002503) | 15 | 5 | 0.01 | + | > 100 | 1.66E-13 | 4.36E-10 |
| MHC class II protein complex assembly (GO:0002399) | 15 | 5 | 0.01 | + | > 100 | 1.66E-13 | 3.74E-10 |
| peptide antigen assembly with MHC protein complex (GO:0002501) | 19 | 5 | 0.01 | + | > 100 | 4.55E-13 | 8.96E-10 |
| MHC protein complex assembly (GO:0002396) | 19 | 5 | 0.01 | + | > 100 | 4.55E-13 | 7.96E-10 |
| antigen processing and presentation of exogenous peptide antigen via MHC class II (GO:0019886) | 24 | 6 | 0.01 | + | > 100 | 1.73E-15 | 2.73E-11 |
| antigen processing and presentation of peptide antigen via MHC class II (GO:0002495) | 26 | 6 | 0.01 | + | > 100 | 2.64E-15 | 2.08E-11 |
| antigen processing and presentation of peptide or polysaccharide antigen via MHC class II (GO:0002504) | 27 | 6 | 0.01 | + | > 100 | 3.23E-15 | 1.7E-11 |
| positive regulation of antigen processing and presentation (GO:0002579) | 12 | 2 | 0.01 | + | > 100 | 2.06E-05 | 0.00637 |
| antigen processing and presentation of exogenous peptide antigen (GO:0002478) | 37 | 6 | 0.02 | + | > 100 | 1.77E-14 | 6.99E-11 |
| positive thymic T cell selection (GO:0045059) | 14 | 2 | 0.01 | + | > 100 | 2.72E-05 | 0.00808 |
| antigen processing and presentation of exogenous antigen (GO:0019884) | 44 | 6 | 0.02 | + | > 100 | 4.62E-14 | 1.46E-10 |
| immunoglobulin production involved in immunoglobulin-mediated immune response (GO:0002381) | 52 | 5 | 0.03 | + | > 100 | 4.45E-11 | 5.39E-08 |
| regulation of antigen processing and presentation (GO:0002577) | 21 | 2 | 0.01 | + | > 100 | 5.71E-05 | 0.0143 |
| chaperone cofactor-dependent protein refolding (GO:0051085) | 33 | 3 | 0.02 | + | > 100 | 6.57E-07 | 0.000288 |
| thymic T cell selection (GO:0045061) | 25 | 2 | 0.01 | + | > 100 | 7.92E-05 | 0.0186 |
| 'de novo' post-translational protein folding (GO:0051084) | 38 | 3 | 0.02 | + | > 100 | 9.8E-07 | 0.000407 |
| 'de novo' protein folding (GO:0006458) | 39 | 3 | 0.02 | + | > 100 | 1.06E-06 | 0.000416 |
| antigen processing and presentation of peptide antigen (GO:0048002) | 81 | 6 | 0.04 | + | > 100 | 1.46E-12 | 2.3E-09 |
| positive T cell selection (GO:0043368) | 36 | 2 | 0.02 | + | > 100 | 0.000158 | 0.0346 |

**Tumor Factor 16 and WH Factor 13**

| **Term** | **Overlap** | **P-value** | **Adjusted P-value** | **Old P-value** | **Old Adjusted P-value** | **Odds Ratio** | **Combined Score** | **Genes** |
| --- | --- | --- | --- | --- | --- | --- | --- | --- |
| type I interferon signaling pathway (GO:0060337) | 6/65 | 8.46E-13 | 4.65E-11 | 0 | 0 | 337.7797 | 9389.69 | RSAD2;OAS3;ISG15;IFIT1;IFIT3;IFIT2 |
| cellular response to type I interferon (GO:0071357) | 6/65 | 8.46E-13 | 4.65E-11 | 0 | 0 | 337.7797 | 9389.69 | RSAD2;OAS3;ISG15;IFIT1;IFIT3;IFIT2 |
| defense response to symbiont (GO:0140546) | 6/124 | 4.5E-11 | 1.65E-09 | 0 | 0 | 168.3898 | 4011.596 | RSAD2;OAS3;ISG15;IFIT1;IFIT3;IFIT2 |
| defense response to virus (GO:0051607) | 6/133 | 6.9E-11 | 1.9E-09 | 0 | 0 | 156.3858 | 3658.92 | RSAD2;OAS3;ISG15;IFIT1;IFIT3;IFIT2 |
| negative regulation of viral genome replication (GO:0045071) | 4/54 | 2.31E-08 | 5.08E-07 | 0 | 0 | 199.38 | 3505.681 | RSAD2;OAS3;ISG15;IFIT1 |
| regulation of viral genome replication (GO:0045069) | 4/67 | 5.58E-08 | 1.02E-06 | 0 | 0 | 158.1349 | 2641.137 | RSAD2;OAS3;ISG15;IFIT1 |
| negative regulation of viral process (GO:0048525) | 4/70 | 6.67E-08 | 1.05E-06 | 0 | 0 | 150.9242 | 2493.807 | RSAD2;OAS3;ISG15;IFIT1 |
| cytokine-mediated signaling pathway (GO:0019221) | 6/621 | 6.89E-07 | 9.47E-06 | 0 | 0 | 31.50081 | 446.9304 | RSAD2;OAS3;ISG15;IFIT1;IFIT3;IFIT2 |
| response to type I interferon (GO:0034340) | 2/9 | 1.19E-05 | 0.000145 | 0 | 0 | 570.8857 | 6475.556 | ISG15;IFIT1 |
| negative regulation of type I interferon-mediated signaling pathway (GO:0060339) | 2/16 | 3.94E-05 | 0.000434 | 0 | 0 | 285.3429 | 2893.761 | OAS3;ISG15 |
| regulation of type I interferon-mediated signaling pathway (GO:0060338) | 2/30 | 0.000142 | 0.001422 | 0 | 0 | 142.5714 | 1262.919 | OAS3;ISG15 |
| positive regulation of interferon-beta production (GO:0032728) | 2/36 | 0.000206 | 0.001884 | 0 | 0 | 117.3765 | 996.4997 | OAS3;ISG15 |
| negative regulation of innate immune response (GO:0045824) | 2/38 | 0.000229 | 0.00194 | 0 | 0 | 110.8444 | 928.9654 | OAS3;ISG15 |
| regulation of interferon-beta production (GO:0032648) | 2/49 | 0.000382 | 0.003002 | 0 | 0 | 84.85532 | 667.807 | OAS3;ISG15 |
| negative regulation of cytokine-mediated signaling pathway (GO:0001960) | 2/55 | 0.000481 | 0.003531 | 0 | 0 | 75.22642 | 574.6283 | OAS3;ISG15 |
| negative regulation of protein binding (GO:0032091) | 2/74 | 0.00087 | 0.005983 | 0 | 0 | 55.32222 | 389.8431 | IFIT1;IFIT2 |
| positive regulation of type I interferon production (GO:0032481) | 2/77 | 0.000942 | 0.006094 | 0 | 0 | 53.10133 | 369.9972 | OAS3;ISG15 |
| negative regulation of binding (GO:0051100) | 2/88 | 0.001228 | 0.007502 | 0 | 0 | 46.28372 | 310.2267 | IFIT1;IFIT2 |
| regulation of protein binding (GO:0043393) | 2/118 | 0.002192 | 0.012689 | 0 | 0 | 34.26207 | 209.79 | IFIT1;IFIT2 |

**Tumor Factor 4 and WH Factor 11**

| **Term** | **Overlap** | **P-value** | **Adjusted P-value** | **Old P-value** | **Old Adjusted P-value** | **Odds Ratio** | **Combined Score** |
| --- | --- | --- | --- | --- | --- | --- | --- |
| cellular response to oxidative stress (GO:0034599) | 4/125 | 6.92E-07 | 7.75E-05 | 0 | 0 | 82.09504 | 1164.369 |
| glutathione metabolic process (GO:0006749) | 3/43 | 2.01E-06 | 0.000113 | 0 | 0 | 166.2333 | 2180.621 |
| glutathione biosynthetic process (GO:0006750) | 2/11 | 1.81E-05 | 0.000426 | 0 | 0 | 443.9778 | 4848.17 |
| nonribosomal peptide biosynthetic process (GO:0019184) | 2/12 | 2.17E-05 | 0.000426 | 0 | 0 | 399.56 | 4290.418 |
| vascular process in circulatory system (GO:0003018) | 2/13 | 2.56E-05 | 0.000426 | 0 | 0 | 363.2182 | 3839.629 |
| regulation of tube diameter (GO:0035296) | 2/13 | 2.56E-05 | 0.000426 | 0 | 0 | 363.2182 | 3839.629 |
| cellular response to chemical stress (GO:0062197) | 3/101 | 2.66E-05 | 0.000426 | 0 | 0 | 67.65306 | 712.6872 |
| sulfur compound biosynthetic process (GO:0044272) | 3/113 | 3.72E-05 | 0.000521 | 0 | 0 | 60.23636 | 614.32 |
| prostanoid metabolic process (GO:0006692) | 2/18 | 5.02E-05 | 0.000625 | 0 | 0 | 249.65 | 2471.302 |
| cellular modified amino acid biosynthetic process (GO:0042398) | 2/20 | 6.23E-05 | 0.000698 | 0 | 0 | 221.8889 | 2148.582 |
| glutamate metabolic process (GO:0006536) | 2/22 | 7.57E-05 | 0.000771 | 0 | 0 | 199.68 | 1894.647 |
| prostaglandin metabolic process (GO:0006693) | 2/31 | 0.000152 | 0.001418 | 0 | 0 | 137.6483 | 1210.175 |
| blood vessel diameter maintenance (GO:0097746) | 2/35 | 0.000194 | 0.001673 | 0 | 0 | 120.9394 | 1033.621 |
| glutamine family amino acid metabolic process (GO:0009064) | 2/37 | 0.000217 | 0.001738 | 0 | 0 | 114.0171 | 961.6818 |
| response to hydrogen peroxide (GO:0042542) | 2/49 | 0.000382 | 0.002853 | 0 | 0 | 84.85532 | 667.807 |
| blood circulation (GO:0008015) | 2/51 | 0.000414 | 0.002898 | 0 | 0 | 81.38367 | 633.9615 |
| dicarboxylic acid metabolic process (GO:0043648) | 2/57 | 0.000517 | 0.003407 | 0 | 0 | 72.48364 | 548.5004 |
| response to reactive oxygen species (GO:0000302) | 2/59 | 0.000554 | 0.003447 | 0 | 0 | 69.93333 | 524.3825 |
| leukotriene B4 metabolic process (GO:0036102) | 1/5 | 0.002997 | 0.016781 | 0 | 0 | 454.1818 | 2638.914 |

**Tumor Factor 5 and WH Factor 12**

| **Term** | **Overlap** | **P-value** | **Adjusted P-value** | **Old P-value** | **Old Adjusted P-value** | **Odds Ratio** | **Combined Score** | **Genes** |
| --- | --- | --- | --- | --- | --- | --- | --- | --- |
| regulation of complement activation (GO:0030449) | 2/50 | 0.000218 | 0.004496 | 0 | 0 | 118.7083 | 1000.806 | C4B;CFH |
| regulation of immune effector process (GO:0002697) | 2/53 | 0.000245 | 0.004496 | 0 | 0 | 111.7087 | 928.7246 | C4B;CFH |
| regulation of humoral immune response (GO:0002920) | 2/54 | 0.000254 | 0.004496 | 0 | 0 | 109.5549 | 906.7098 | C4B;CFH |
| detection of molecule of bacterial origin (GO:0032490) | 1/7 | 0.003146 | 0.026788 | 0 | 0 | 416.3542 | 2398.853 | C4B |
| folic acid transport (GO:0015884) | 1/8 | 0.003595 | 0.026788 | 0 | 0 | 356.8571 | 2008.476 | FOLR2 |
| positive regulation of apoptotic cell clearance (GO:2000427) | 1/8 | 0.003595 | 0.026788 | 0 | 0 | 356.8571 | 2008.476 | C4B |
| regulation of apoptotic cell clearance (GO:2000425) | 1/8 | 0.003595 | 0.026788 | 0 | 0 | 356.8571 | 2008.476 | C4B |
| regulation of complement-dependent cytotoxicity (GO:1903659) | 1/9 | 0.004043 | 0.026788 | 0 | 0 | 312.2344 | 1720.615 | CFH |
| sodium ion export across plasma membrane (GO:0036376) | 1/13 | 0.005836 | 0.030275 | 0 | 0 | 208.1146 | 1070.484 | FXYD2 |
| positive regulation of sodium ion transmembrane transporter activity (GO:2000651) | 1/14 | 0.006284 | 0.030275 | 0 | 0 | 192.0962 | 973.8925 | FXYD2 |
| positive regulation of potassium ion transmembrane transporter activity (GO:1901018) | 1/14 | 0.006284 | 0.030275 | 0 | 0 | 192.0962 | 973.8925 | FXYD2 |
| regulation of cell killing (GO:0031341) | 1/17 | 0.007625 | 0.031088 | 0 | 0 | 156.0547 | 760.9634 | CFH |
| modified amino acid transport (GO:0072337) | 1/17 | 0.007625 | 0.031088 | 0 | 0 | 156.0547 | 760.9634 | FOLR2 |
| folic acid metabolic process (GO:0046655) | 1/19 | 0.008519 | 0.031679 | 0 | 0 | 138.7014 | 660.9724 | FOLR2 |
| folic acid-containing compound metabolic process (GO:0006760) | 1/20 | 0.008966 | 0.031679 | 0 | 0 | 131.3947 | 619.4396 | FOLR2 |
| regulation of sodium ion transmembrane transport (GO:1902305) | 1/28 | 0.012532 | 0.036118 | 0 | 0 | 92.42593 | 404.7762 | FXYD2 |
| amide transport (GO:0042886) | 1/28 | 0.012532 | 0.036118 | 0 | 0 | 92.42593 | 404.7762 | FOLR2 |
| actin filament polymerization (GO:0030041) | 1/29 | 0.012977 | 0.036118 | 0 | 0 | 89.12054 | 387.1908 | GAS7 |
| dicarboxylic acid transport (GO:0006835) | 1/29 | 0.012977 | 0.036118 | 0 | 0 | 89.12054 | 387.1908 | FOLR2 |

**Tumor Factor 7 and WH Factor 15**

| **Term** | **Overlap** | **P-value** | **Adjusted P-value** | **Old P-value** | **Old Adjusted P-value** | **Odds Ratio** | **Combined Score** | **Genes** |
| --- | --- | --- | --- | --- | --- | --- | --- | --- |
| mitotic chromosome condensation (GO:0007076) | 2/27 | 9.58E-05 | 0.009293 | 0 | 0 | 177.4578 | 1642.053 | SMC4;SMC2 |
| chromosome condensation (GO:0030261) | 2/45 | 0.000269 | 0.013035 | 0 | 0 | 103.0801 | 847.4912 | SMC4;SMC2 |
| positive regulation of binding (GO:0051099) | 2/90 | 0.001073 | 0.033336 | 0 | 0 | 50.25505 | 343.6227 | STMN1;HMGB2 |
| mitotic sister chromatid segregation (GO:0000070) | 2/102 | 0.001375 | 0.033336 | 0 | 0 | 44.19778 | 291.2425 | SMC4;SMC2 |
| regulation of guanyl-nucleotide exchange factor activity (GO:1905097) | 1/5 | 0.002747 | 0.046367 | 0 | 0 | 499.625 | 2946.374 | STMN1 |
| regulation of nuclease activity (GO:0032069) | 1/6 | 0.003296 | 0.046367 | 0 | 0 | 399.68 | 2284.211 | HMGB2 |
| DNA topological change (GO:0006265) | 1/9 | 0.00494 | 0.046367 | 0 | 0 | 249.7625 | 1326.334 | HMGB2 |
| positive regulation of megakaryocyte differentiation (GO:0045654) | 1/9 | 0.00494 | 0.046367 | 0 | 0 | 249.7625 | 1326.334 | HMGB2 |
| V(D)J recombination (GO:0033151) | 1/11 | 0.006035 | 0.046367 | 0 | 0 | 199.79 | 1020.969 | HMGB2 |
| hepatocyte growth factor receptor signaling pathway (GO:0048012) | 1/11 | 0.006035 | 0.046367 | 0 | 0 | 199.79 | 1020.969 | STMN1 |
| negative regulation of microtubule polymerization (GO:0031115) | 1/12 | 0.006582 | 0.046367 | 0 | 0 | 181.6182 | 912.3494 | STMN1 |
| regulation of chromatin organization (GO:1902275) | 1/14 | 0.007675 | 0.046367 | 0 | 0 | 153.6615 | 748.3003 | MKI67 |
| apoptotic DNA fragmentation (GO:0006309) | 1/16 | 0.008767 | 0.046367 | 0 | 0 | 133.16 | 630.7475 | HMGB2 |
| regulation of stem cell proliferation (GO:0072091) | 1/17 | 0.009313 | 0.046367 | 0 | 0 | 124.8313 | 583.7595 | HMGB2 |
| negative regulation of Rho protein signal transduction (GO:0035024) | 1/18 | 0.009858 | 0.046367 | 0 | 0 | 117.4824 | 542.7074 | STMN1 |
| microtubule depolymerization (GO:0007019) | 1/18 | 0.009858 | 0.046367 | 0 | 0 | 117.4824 | 542.7074 | STMN1 |
| DNA conformation change (GO:0071103) | 1/18 | 0.009858 | 0.046367 | 0 | 0 | 117.4824 | 542.7074 | HMGB2 |
| regulation of chromosome segregation (GO:0051983) | 1/18 | 0.009858 | 0.046367 | 0 | 0 | 117.4824 | 542.7074 | MKI67 |
| regulation of water loss via skin (GO:0033561) | 1/19 | 0.010403 | 0.046367 | 0 | 0 | 110.95 | 506.5602 | STMN1 |
