## Supplemental Table 3 for "Space-Time Mapping Identifies Concerted Multicellular Patterns and Gene Programs in Healing Wounds and their Conservation in Cancers"

**Supplementary Table S3**

| Figure | Surface / Spot | Surface / Spots name | Channel | Estimated XY Diameter | Estimated Z Diameter | Smoothing: Surface Grain Size | Background Subtraction: Diameter of largest sphere | Threshold: Manual Threshold Value | Threshold: Manual Threshold Value B | Threshold: Region Growing Estimated Diameter | Classify Spots/Seed points: Quality Threshold | Classify Surfaces: Number of Voxels | nuclei close to surface | CD11b colocated |
| --- | --- | --- | --- | --- | --- | --- | --- | --- | --- | --- | --- | --- | --- | --- |
| Figure 3, Day 3 | Spot | nuclei | DAPI | 8 µm | 30 µm |  | true |  |  |  | automatic |  |  |  |
| Arg1-tdTomato | Surface | Arg1 | tdTomato |  |  | 2.71 µm | 11 µm | 10 | 177.116 | 11 µm | >5.39 | >60 | 10 µm | <1 µm |
|  | Surface | CD11b | AF647 |  |  | 2.71 µm | 11 µm | 5 | 152.962 | 11 µm | >4.5 | >50 | 20 µm |  |
| Figure 3, Day 7 | Spot | nuclei | DAPI | 10 µm | 30 µm |  | true |  |  |  | >6.99 |  |  |  |
| Arg1-tdTomato | Surface | Arg1 | tdTomato |  |  | 2.71 µm | 10 µm | 21.7648 | 183.446 |  |  | >10 | 10 µm | <1 µm |
|  | Surface | CD11b | AF647 |  |  | 2.71 µm | 10 µm | 6 | 166.593 |  |  | >10 | 20 µm |  |
| Figure 3, Day 3 | Spot | nuclei | DAPI | 8 µm | 30 µm |  | true |  |  |  | automatic |  |  |  |
| CD206 | Surface | CD206 | AF647 |  |  | 2.71 µm | 12 µm | 7.11314 | 191.68 |  |  | >30 | 5 µm | <1 µm |
|  | Surface | CD11b | AF594 |  |  | 2.71 µm | 12 µm | 6.6 | 172.95 |  |  | >50 | 5 µm |  |
| Figure 3, Day 7 | Spot | nuclei | DAPI | 8 µm | 30 µm |  |  |  |  |  | >16.8 |  |  |  |
| CD206 | Surface | CD206 | AF647 |  |  | 2.71 µm | 11 µm | 12.8679 | 178.637 |  |  | >10 | 5 µm | <1 µm |
|  | Surface | CD11b | AF594 |  |  | 2.71 µm | 12 µm | 7.56092 | 184.342 |  |  | >10 | 5 µm |  |
